# Adaptive walks on high-dimensional fitness landscapes and seascapes with distance-dependent statistics

**DOI:** 10.1101/435669

**Authors:** Atish Agarwala, Daniel S. Fisher

## Abstract

The dynamics of evolution is intimately shaped by epistasis — interactions between genetic elements which cause the fitness-effect of combinations of mutations to be non-additive. Analyzing evolutionary dynamics that involves large numbers of epistatic mutations is intrinsically difficult. A crucial feature is that the fitness landscape in the vicinity of the current genome depends on the evolutionary history. A key step is thus developing models that enable study of the effects of past evolution on future evolution. In this work, we introduce a broad class of high-dimensional random fitness landscapes for which the correlations between fitnesses of genomes are a general function of genetic distance. Their Gaussian character allows for tractable computational as well as analytic understanding. We study the properties of these landscapes focusing on the simplest evolutionary process: random adaptive (uphill) walks. Conventional measures of “ruggedness” are shown to not much affect such adaptive walks. Instead, the long-distance statistics of epistasis cause all properties to be highly conditional on past evolution, determining the statistics of the local landscape (the distribution of fitness-effects of available mutations and combinations of these), as well as the global geometry of evolutionary trajectories. In order to further explore the effects of conditioning on past evolution, we model the effects of slowly changing environments. At long times, such fitness “seascapes” cause a statistical steady state with highly intermittent evolutionary dynamics: populations undergo bursts of rapid adaptation, interspersed with periods in which adaptive mutations are rare and the population waits for more new directions to be opened up by changes in the environment. Finally, we discuss prospects for studying more complex evolutionary dynamics and on broader classes of high-dimensional landscapes and seascapes.

## 1 Introduction

Evolution, even in the short term for “simple” asexual microbial populations with “simple” selective pressures, is a complex, non-linear dynamical process. One of the many sources of complexity is *epistasis* – the interactions between combinations of mutations. Most simply, pairs of mutations can have no benefit individually but benefits together, positive effects alone but deleterious ones together, and, once many mutations are involved, a plethora of other behaviors. These epistatic effects lead to correlations and anti-correlations in cooccurrence of mutations, which can have a dramatic effect on evolutionary trajectories. In trying to model epistasis, an important observation is that, generally, whether a mutation is beneficial or deleterious, and by how much, will depend on the both the environment and the genomic background, and hence the evolutionary history: any mutation that is unconditionally beneficial would have already occurred and fixed. Thus the details of epistasis will depend greatly on the context and one should hope for primarily statistical understanding.

At a basic level, we lack a theoretical understanding of the interplay between evolution and epistasis. There is a great deal of population genetic theory in cases where mutations are independent, and their contribution to the (log) fitness is additive [12, 18]. But, except in simple cases involving small numbers of mutations, little is understood generally about evolutionary dynamics in the presence of epistatic interactions. Many studies have focused on the statistical properties of epistasis by constructing and attempting to understand models of *fitness landscapes* – functions from genotype to fitness in a fixed environment. These are often conceptualized like physical landscapes with peaks and valleys [10], and theorists have endeavored to understand how properties of these landscapes constrain or enable evolution [49, 51, 55]. Some approaches have focused on quantitative “local” properties, such as using extreme value statistics to understand behaviors near fitness peaks [53]. Others have focused on quantifying the geometry of fitness landscapes, including distributions of local maxima and numbers of uphill paths between points [11, 33, 71]. There is often an emphasis placed on defining and quantifying measures of “ruggedness”, with the intuition that rugged landscapes constrain evolution by limiting the number of routes along which the fitness can continue to increase, leading to getting stuck at local maxima [10]. But ruggedness is not well defined in general, and, as we shall see, local measures of it can be misleading. Moreover, there is an intrinsic problem with focusing primarily on fitness landscapes themselves. The interplay between epistasis and evolution is essential; one cannot try to separately understand them as the properties of the landscape in the vicinity of the current genome are highly conditioned by the past evolution whether in a fixed or changing environment. The properties of epistatic fitness landscapes *conditioned* on past evolution will generally be very different than the properties of “typical” points or regions on such landscapes. Even if epistasis is some sense “weak”, over long times individuals will accumulate a large number of mutations [2] and thus conditioning on history will eventually become important.

A major conceptual difficulty is that fitness landscapes are very high dimensional: there are many potentially beneficial mutations and even more combinations of them. Yet intuition about landscapes and evolution in them is usually based on low-dimensional analogies. A large body of work in theoretical physics and probability theory has shown that geometry and dynamics in high-dimensional spaces, such as uphill paths on complex landscapes, can be very different than in their low dimensional analogues [7, 8].

Various experiments have provided some information about epistasis in natural systems. Extensive work has been done on fitness effects of combinations of pairs of knockouts in various microbes, in particular *S. cerevisiae* and *E. coli* [14, 30, 68]. However most knockouts are deleterious, and pairs of them at least as much so. But evolution is largely driven by beneficial mutations, thus it is not clear how much is learned from such experiments about statistical properties of epistasis that would affect evolutionary dynamics. High throughput studies of repeatedly mated yeast populations [5] have provided potentially valuable information, but even with some good combinations of genetic differences being produced, it is not clear what are useful measures of the features of epistasis that affect evolution. Furthermore, these measurements have thus far combined genomes that differ by large numbers of mutations, thus they are not directly relevant for the simpler problem of asexual evolution, our focus here, for which the landscape is explored locally with distant genomes having to be reached step by step.

In some relatively simple systems, a variety of *empirical fitness landscapes* — functions from genotype to measured fitness — have been analyzed and the consequences for evolution investigated theoretically. However, by their nature, these are restricted to exploring combinations of genomic changes on only a modest number of sites - typically in a single protein or RNA sequence [4, 33, 39, 71, 73]. While these studies can yield insights about specific evolutionary scenarios, such as antibiotic resistance caused by multiple mutations in a small number of proteins under strong, focused selective pressure, drawing general conclusions from these, especially about much higher dimensional landscapes, is problematic.

Some experiments have explored the interplay between epistasis and evolutionary dynamics by laboratory microbial evolution [58, 71, 74]. Much has been learned about the fitness effects of individual beneficial mutations [34, 38, 69], but thus far only limited amounts about epistatic interactions between the mutations found, beyond a general tendency towards “diminishing returns” epistasis: the net effect of beneficial combinations of mutations being less than the sum of their individual effects. However, this is not true of all combinations and it may well be that such anomalous combinations are particularly important for driving further evolution [32]. Indeed, there is evidence that a common motif in protein evolution is a beneficial but destabilizing mutation followed by a stabilizing one - a type of positive epistasis [20].

All experimental studies face a spectrum of intrinsic challenges. One is the combinatorial explosion of possible genotypes, which is exponential in the number of sites being studied. This combinatorial explosion is associated with another key feature of epistasis: at large genetic distances, higher order interactions are important. When many genotypes are involved it is not enough to consider only pairwise interactions: correlations involving many sites on the genome become important. Indeed, “expanding” around a particular genome — the ancestor — in additive, pairwise, triplet interactions, and so on, is problematic. This is at best a way to parameterize the fitness landscape in the vicinity of a particular genome which is not essentially special: such an expansion could be already substantially different around another genome not far away.

Another layer of complexity comes from the dynamic nature of environments in which organisms find — and have found — themselves. The landscapes are really “seascapes”, and typically change on timescales relevant for even short term evolutionary dynamics [3, 25, 26]. These changes can be driven by external physical and chemical factors, as well as changes in direct interactions between organisms or feedback on the environment by evolving organisms that populate it. As the net effects of mutations can be a delicate balance between positive and negative effects, even small changes in environment can change which mutations are beneficial and more generally the possible routes to adaptation via multiple successive mutations. Recent experiments have provided hints at how strong the effects of environmental changes caused by evolution are even in nominally “simple” conditions. This is seen for the first mutations that occur in yeast in low glucose [40] as well as from cumulative effects from many mutations in Lenski’s long-term *E. coli* evolutions [23]. On long timescales in natural populations, dynamical changes in the environment, whether produced by abiotic changes or by the evolving populations themselves, are surely major drivers of adaptation.

With large numbers of sites involved in evolution, all one can hope for, generally, is to understand statistical properties of landscapes (and seascapes). New, more general theoretical approaches are vitally needed, both to develop intuition, and to guide future experiments and choose what statistical properties would be most instructive to measure. It is imperative that theorists build and study models that include caricatures of these key features: high-dimensional landscapes with some general statistical forms of epistasis, and, in the same framework, gradual time-dependence of the landscape (about which one can hope more general can be said than about large sudden changes in the environment). Models should have no “special genomes” — such as “wild-type” — a priori, and instead one should carry out evolution from a random point and then try to understand the effects of epistasis and further evolution conditioned on past evolution. Some recent work has endeavored to analyze evolutionary dynamics on some classes of landscapes with simple models of epistasis, [29, 49, 51, 56, 57]; however these analyses are computationally limited from exploring more general, or more high-dimensional landscapes.

In this work, we will develop understanding of the feedback between evolution and epistasis in a particular rich class of toy models. Motivated by the basic observation that effects of genetic changes are sums of many positive and negative contributions, we study (following others [29, 35, 49]) *random* fitness landscapes properties are characterized statistically. We develop a theoretical framework to understand a class of random fitness landscapes with *distance-dependent statistics* where the correlations between the fitnesses of genomes are some function of the genetic distances between them. Our model class is flexible, with desirable biologically inspired features, and encompasses — while illuminating some problems with — many of the most commonly studied models of epistasis. By working in the tree-like limit of genotype space where there are many possible mutations and reversions are rare (as in the infinite-sites approximation) we develop a computational and analytical scheme which can be used to analyze simple evolutionary dynamics on arbitrarily high dimensional landscapes, as well as giving insights into key geometric properties of the landscapes.

Our analysis shows that evolution drives populations to highly atypical places on the fitness landscape — atypical far beyond just having anomalously high fitness. The patterns of epistasis in the neighborhood of the current genome, depends on the whole evolutionary trajectory of the population, which itself depends on long-genomic-distance (arbitrarily high order) patterns of epistasis. This is in part due to the high dimensionality of the fitness landscape, which allows rare events — although ones that evolution finds readily — to dominate the dynamics. Using our framework we compute how far evolution takes populations before getting stuck at local maxima; we show that greedy evolutionary dynamics can be detrimental in the long term, and that the process of getting stuck depends sensitively on the tails of the distribution of mutational effects. We then analyze slowly time-dependent landscapes for which adaptive walks never get fully stuck. We show that when the fitness landscape has a simple time-dependence, the conditioning on the past is limited. Nevertheless, the atypical nature of trajectories and local landscapes remains. We expect the results to hold qualitatively for more general time-dependence.

We conclude with potential directions for future theoretical studies and potentially informative experiments, particularly in light of our findings that intermediate-term evolution even with simple models of effectively-random epistasis are set by *long-distance*, *global* statistical properties of the landscape, rather than *short-distance*, *local* information such as measures of “ruggedness”.

## 2 Fitness landscapes with distance-dependent divergence

### 2.1 Random fitness landscapes

We begin with some general qualitative and quantitative aspects of fitness landscapes and epistasis. The main quantity of interest is the fitness function *F* (*g*), which maps genotypes, *g*, to fitness values. The genome is most simply modeled by strings of 0’s and 1’s of length *L* – points on a hypercube. Recent work suggests the number of potential states per site (four possible DNA base pairs, or 20 possible amino acids in a protein chain) might play a quantitative and qualitative role in the dynamics of adaptation [75]. However such complexity might most affect the local rather than large-scale behavior, especially with large *L* (our regime of interest), and is therefore beyond the scope of our work.

A common convention is that all zeros corresponds to some ancestral/reference genotype, with 1’s then representing mutations; we will not use that convention here as we will not want to assign special characteristics to any ancestor. Distances between genomes, defined as the number of sites at which they differ, will be a fundamental property in our models: of course, for these, the labeling of the reference genome is arbitrary.

The fitness function *F* (*g*) is often called the *fitness landscape*. Evolution tends to drive the population “up” the landscape to higher and higher fitness. A basic idea is that these upward trajectories are impeded by “ruggedness” with local maxima and other features that make some paths easier to traverse than others. In this work, we will study *random fitness landscapes*: ensembles of some classes of random functions over genotype space [29, 35, 49]. The use of a random model is motivated by the fact that “fitness” is a very complex quantity. The net effect of any mutation or combination of mutations in a relatively well-adapted organism will be the sum of beneficial and deleterious effects, all conditioned on the evolutionary history and dependent on the particular environment. The hope is that a caricature of the landscape as a random function will capture some of the essence of the complex interplay between epistasis and evolution. Most previous theoretical work on fitness landscapes falls into two categories, focusing either on some classes of random landscape models [19, 21, 35, 49, 73] as do we, or on models of specific simple situations like mutations in a single protein [71] or aspects of the fitness of particular viruses [41].

We would like to study broad classes of models that have certain key properties motivated by the desire to caricature biologically plausible/relevant features, and, secondarily, by useful analytical properties:

1. **Good scaling properties for large genomes:** We want to have well-characterized behavior in the limit of large genome sizes, but total distances between genomes much less than *L*, in particular avoiding pathologies related to the combinatorics of large genomes that are drawbacks of some oft-studied forms of epistasis.
2. **Evolutionary-conditioned statistics:** We wish to understand evolution conditioned on past evolution and thus regions of the landscape that the population is led to by evolution. We would like such historical conditioning to be derived from the model as opposed to put in by hand. In particular, we want to be careful not to assign any exceptional significance to a “reference” or “ancestral” genotype.
3. **Flexibility:** We want a modeling framework that encompasses models with a large variety of qualitative and quantitative structure in order to broadly understand the effects of epistasis, including the effects of both small numbers of mutations and genomes that are much further apart: i.e., both short-distance and long-distance structure.
4. **Tractable dynamics:** We need to be able to develop theoretical and computational means to explore evolutionary dynamics for long times and large genome sizes, with methods that scale well with genome length. This requires being able to efficiently generate and store pertinent information about the landscape in simulations.

We can satisfy the first three criteria by defining fitness landscapes using *distance-dependent divergence functions* (related to the recently defined *γ* measure [17]). We thus choose *F* from an ensemble of random functions whose statistics depend only on the genetic distances between genotypes. A fundamental property of the ensemble is the statistical relationship between genomes as a function of the distance between them. We define the *divergence function* of fitness differences as

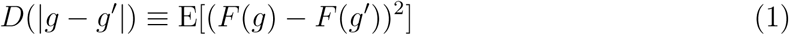

for any pair of genomes *g* and *g′*. This average is taken over the ensemble of random models. Here *D* represents the relationship between the genetic distance *|g −g′|* between two genomes and their distance in fitness space *F* (*g*) *− F* (*g′*). The choice to quantify the model in terms of fitness differences is in fact natural as relative fitness drives evolution. Note that the correlations between any pair of fitness differences can be expressed in terms of *D*(𝓁):

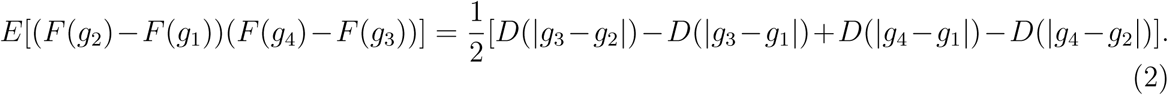

This means that studying models in terms of *D* is equivalent to studying correlations between the effects of single mutations (as is true for the *γ*-measure [17]).

With statistical properties defined in terms of genetic and fitness differences, no genome is special with respect to the ensemble as a whole. This is in contrast to defining models in terms of the distribution of fitness effects (DFE) of potential single mutations around a particular genome. For a general random landscape, the DFE depends on the evolutionary history: in particular, with increasing fitness the supply and magnitude of potential adaptive mutations both decrease. This statistical phenomenon is called diminishing-returns epistasis, and there are many models which are *defined* by this structure [19, 21, 73]. In contrast, we would like to understand the emergence of effects like diminishing-returns epistasis in a more general framework, by studying how well-adapted genomes are special due to the effects of past evolution. Our models allow us to specify global features of the epistasis, then let the entire history of the evolutionary dynamics determine the current DFE and epistasis among combinations of mutations around the current genome. This approach lets us generate and study evolution-conditioned regions of the fitness landscape [29, 49, 51, 55], rather than putting in the structure of the “special” nature of a well-adapted genome in by hand.

While we will primarily be interested in fitness differences, in some landscapes, the absolute fitness is well-defined and is hence an (albeit unmeasurable) characteristic of the initial genome. The absolute fitnesses scale as *D*(*L*)^1^*^/^*^2^. If *D*(*L*) = *O*(1) for large *L* then the absolute fitnesses are well defined in the infinite *L* limit on which we will primarily focus; if instead *D*(*L*) scales as some increasing function of *L*, the absolute fitnesses diverge in the infinite *L* limit. However, fitness differences (which determine evolutionary dynamics) are always well defined.

To get numerically tractable models with large genomes, and even more so to enable general analysis of the evolutionary dynamics and effects of conditioning on past evolution, we must make major simplifications. We will thus (for the most part) assume that *F* (*g*) is a Gaussian random function, and therefore the choice of *D* uniquely defines our models. The choice of Gaussian-random landscapes is certainly not biologically motivated, but it will enable us to begin to address general questions and develop computational and analytical tools. Understanding evolution on non-Gaussian landscapes is a major challenge: in the Discussion, we make some remarks about how some analyses might begin.

To understand the relationship between formulations of *F* (*g*) in terms of epistatic interactions and the divergence function *D*(𝓁) of Gaussian random landscapes, it is instructive to first consider the two simplest models. First, is the *additive model* for which the effects of each mutation are statistically independent so that the fitness can just be represented as the sum of terms for each site with no epistasis. This corresponds to the choice *D*(𝓁) *∝ 𝓁*. At the other extreme is the *independent fitnesses model* for which the fitness *F* (*g*) of any genotype is independent of the fitness of all other genotypes. This corresponds to the choice *D*(𝓁) = 0 for 𝓁 = 0 and *D*(𝓁) equal to some positive constant for 𝓁 > 0. The independent fitnesses model can be considered a “maximally epistatic” model.

It is informative to compute the correlation between the fitness effect of mutations a distance 𝓁 apart along a trajectory through the landscape: ((e.g. the first and 𝓁 + 1th mutations). This correlation is given by the discrete second derivative 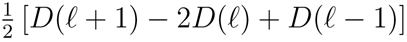^1^. In the additive model, the second derivative vanishes corresponding to no correlation between effects of mutations (no epistasis); by contrast, in the independent fitnesses model, the effects of one mutation and the next are highly anticorrelated, but future mutations are uncorrelated with the first.

Another simple model is called the Rough Mount Fuji (RMF) model; the fitness function is a linear combination of an additive and an independent model [49]. The RMF model thus interpolates between the additive and independent model, but as we will see (Section 4.1), in the large *L* limit for the evolutionary dynamics, it will behave essentially like the additive model.

Another class of models that also interpolates between additive and independent limits has correlations between fitnesses that fall-off exponentially with distance:

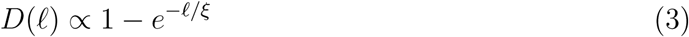

for some correlation length *ξ*. The effects of mutations are then anticorrelated over the length-scale *ξ*, with correlations 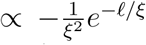. For *ξ →* 0 this becomes the independent-fitnesses model, while for *ξ → ∞* (and rescaling to keep the fitness-differences of order unity), it becomes the additive model. More generally, for evolution on scales 𝓁 ≪ ξ, the landscape is approximately additive while for 𝓁 ≫ ξ fitnesses are approximately independent. We will show that the *NK* model [35], composed of many blocks of interacting sites (defined in the next section), becomes an exponentially correlated Gaussian model in the limit of a large number of blocks [29].

A particularly interesting class of models we will analyze are those with *power law correlations*:

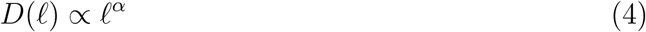

with 0 *< α <* 1. The limit *α* = 0 corresponds to independent fitnesses, while *α* = 1 corresponds to the additive model. But in contrast to exponentially-decaying-correlation models that also interpolate between these simple limits, the power-law correlation models do not have a characteristic genetic distance scale.

Figure 1 shows examples of linear cuts through landscapes for a variety of *α*. For intermediate *α*, *D* is unbounded as in the additive case, but the effects of consecutive (and subsequent) mutations are still anticorrelated. Large fitness differences exist on the landscape and are potentially evolutionarily accessible, but the anti-correlations are felt at long range. We will see that this structure can lead to evolutionary dynamics for which the effects of the diminishing-returns epistasis are weak enough to allow avoidance of local maxima for long periods of time, but they eventually slow down the rate of fitness gain.

**Figure 1:**
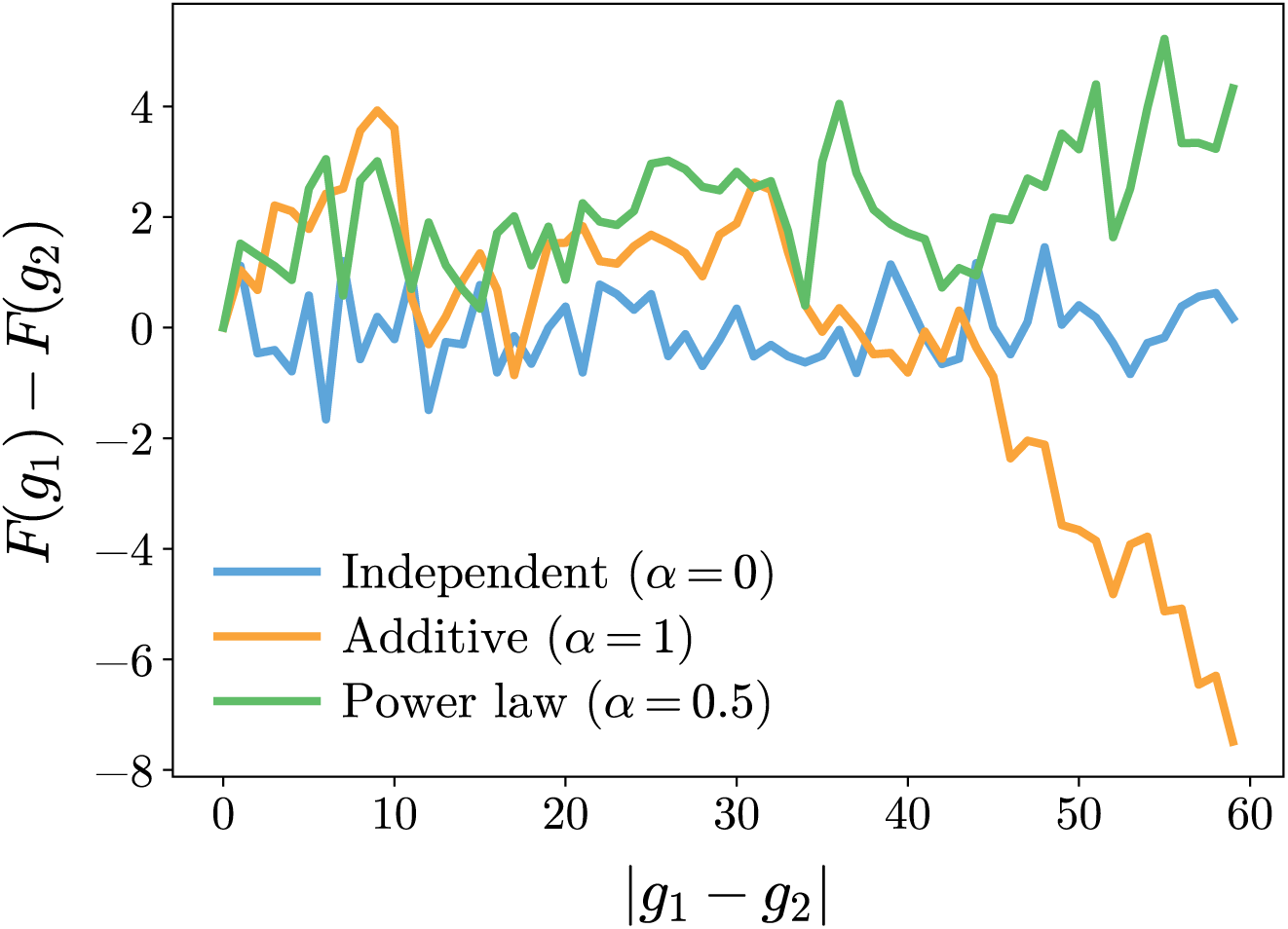
Examples of power-law-correlated fitness landscapes characterized by divergence function *D*(𝓁) = 𝓁^α^ with *α* = 0, 0.5, 1. Shown are typical fitnesses along a random path on the landscape as a function of genomic distance: the number of mutations.

### 2.2 Amplitude spectra and relationship to other models

A common way to characterize epistasis is to break down the fitness into sums of contributions from *k*-wise interactions, with a measure of the overall contribution from the of *k*-wise interactions parameterized by the *amplitude spectrum*, *A_k_* [47, 48, 70]. We now show how to convert from the amplitude spectrum to an equivalent Gaussian model with distance-dependent divergence, and derive the amplitude spectra of some of the models previously defined.

The amplitude spectrum characterizes epistasis by measuring the square magnitude of interactions at each order. It can conveniently be written in terms of the discrete Fourier transform over the hypercube:

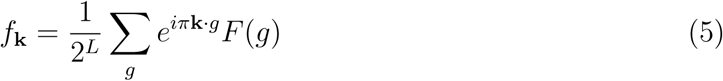

with **K** a vector with *k* ones and *L − k* zeros. Each Fourier component *f***_K_** represents the interactions between a particular set of *k* sites: those with ones in the vector **K**. In the distance-dependent model, the symmetry under permutation of the sites in the genome implies that the statistical properties of *f***_K_** should only depend on the number, *k*, of interacting sites. Their distribution is fully characterized by the amplitude spectrum *A_k_*defined by

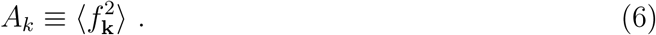

As each Fourier coefficient only depends on *k* sites of the genome, *A_k_* is exactly the mean-square *k*th order global epistasis; *A*_0_ the variance in average fitness (when this is finite), *A*_1_ the variance of the single-site (additive) terms in the fitness, *A*_2_ the mean-square magnitude of pairwise epistatic interactions, and so on up to order *L*. The Fourier coefficients *f***_K_** completely define the covariance matrix, and there is a one-to-one correspondence between the *A_k_*and the distance-dependent divergence *D*(𝓁). For Gaussian landscapes, this amounts to a complete definition of the model.

Note that we define the epistasis without reference to any particular genome, in contrast to expanding in orders of epistasis about some “ancestral” background as 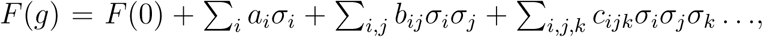 where *σ_i_* = 1 if site *i* is mutated and 0 otherwise. Our definition correctly captures the magnitude of global epistasis. The differences between the two approaches were discussed originally in [70], and more recently in [59].

In the limit of large genomes (large *L*), there are problems with the scaling of *A_k_*. We would like to study behavior that, in this limit, is independent of *L* as long as the distances between genomes are much less than *L*. For example, if the fitness was a sum of single-site terms and pairwise epistatic terms, then one has to decide how the only non-zero coefficients, *A*_1_, and *A*_2_, depend on *L*. If these do not scale with *L*, then the effect of a single mutation would be pathological: the additive piece would contribute *O*(1) while the pairwise terms would contribute 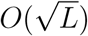.

From this example, we can also see the issues with having only a fixed number of non-zero *A_k_* as *L → ∞*. This remains true even if we rescale terms by factors of *L*. Returning to the pairwise epistasis example, the second order terms could be scaled down such that 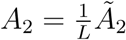 with 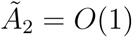, so their total magnitude matches the magnitude of the first order terms — as in the conventional Sherrington-Kirkpatrick spin-glass model [64]. However, for changes of 𝓁 ≪ L sites, the fraction of second order terms where both sites change is 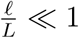. Therefore over distance scales *≪ L*, this model is effectively additive, with 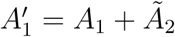.

Another problem occurs if all orders of epistasis are comparably large: i.e., all *A_k_* of the same order. The number of terms (vectors **K**) of order k is 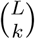. If the A_k_ are all non-zero and don’t scale with *L*, then the middle terms with 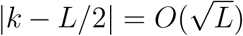 dominate for large *L*. We will see that this gives the independent fitnesses model.

By construction, our distance-dependent ensembles avoid these large *L* pathologies since *D*(𝓁) is independent of *L*. To relate these models to the amplitude spectra and make the large *L* limit well-behaved one needs to define a *rescaled amplitude spectrum*. A convenient choice is to pull out the binomial factor for the number of terms of each order and define *z* = *k/L* as the fraction of sites involved, and the rescaled amplitude spectrum

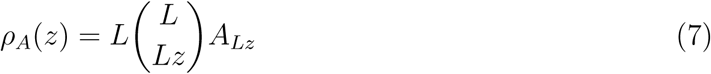

(where the factor of *L* comes from the change of variables). For large *L*, *ρ_A_*(*z*) can be defined continuously over the full range *z ∈* [0, 1]. The *ρ_A_*(*z*) parameterize the magnitude of the *total* epistasis at order *zL*. The *A_k_* (and therefore *ρ_A_*(*z*)) can be computed directly from *D*(𝓁). Similarly, one can use *ρ_A_*(*z*) to compute *D*(𝓁): in Appendix A.2 we show that in the limit of large *L*

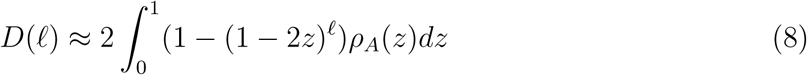

Models with a well defined *ρ_A_* in the limit of large *L* thus correspond to well-behaved distance-dependent divergence functions *D*(𝓁).

Most previously studied models correspond to simple forms of *ρ_A_*(*z*). The additive model has *ρ_A_*(*z*) = *δ*(*z*) (with the limit needing to be taken carefully) while the independent model is *ρ_A_*(*z*) = *δ*(*z −* 1*/*2) since all Fourier modes in it have equal weight. More generally, terms at different orders of epistasis contribute features of different length-scales to *D* with, small *z* — thus small *k* — corresponding to large length-scale features (just as for Fourier transforms). If a single 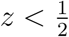 dominates, *C*(𝓁) falls off exponentially with correlation length *ξ* = *−* log(1 *−* 2*z*)*^−^*^1^. Terms with 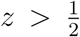 cause oscillations in correlations and make the landscape more “rugged”; in the extreme limit, pure *z* = 1 corresponds to the parity function where every step leads to a change in sign of the fitness. The Random-Mount-Fuji (RMF) model is a sum of additive and independent parts and thus corresponds simply to the sum of delta functions in *ρ_A_* at *z* = 0 and *z* = 1*/*2.

Kauffman’s *NK* model [35] of random landscapes is equivalent, in a certain limit, to a simple distance-dependent model, as first noted in [48]. We rederive the result here to demonstrate the utility of the rescaled amplitude function. The *NK* model splits the contribution to the fitness function into *N* (potentially overlapping) blocks *B_m_*, each consisting of *K* interacting sites. For each block of interactions there is an independent fitness model on the *K* sites in the block, which assigns an i.i.d. random value, *f_B_m__* (*g_B_m__*), to each of the 2*^K^* configurations of the sub-genome g_B_m__, of that block. The total fitness is then 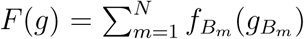. If the *N* blocks are chosen at random, and *L* and *N* are large, a central-limit-theorem-like argument shows that *F* limits to a Gaussian random function. More precisely, one can show that for 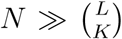, when all blocks are represented many times, all the moments of *F* converge to that of a a jointly Gaussian random function with 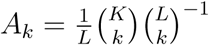. The rescaled amplitude spectrum of the *NK* model is thus simply *ρ_A_*(*z*) *≈ δ*(*z − K/*2*L*). This corresponds to a distance-dependent divergence function with

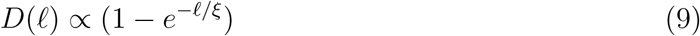

Where 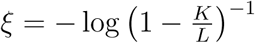.

Typically the *NK* model is studied in a different limit where *N* is *O*(*L*), so only a small subset of all possible blocks are represented [35, 48, 51]. The Gaussian limit corresponds to the mean field *NK* model introduced in [29], with an exponential decay of correlations with the same *ξ* for *K ≪ L*. The analysis of the Gaussian distance-dependent model should still be useful beyond the Gaussian limit. As long as the within-block fitnesses are not broadly distributed, the fitness difference between two genomes at a genetic distance 𝓁 is approximately Gaussian if 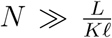, corresponding to many blocks having at least one mutation. For 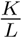 small (long correlation length), this corresponds to roughly 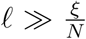. This suggests that the approximation of the *NK* model as a distance-dependent model may hold over a variety of scales, from *x ≪ ξ*, for which each block has at most one mutation and the landscape is approximately additive, to *x ≫ ξ*, for which most blocks have a mutation and the landscape is approximately independent.

The power law model *D*(𝓁) *∝ 𝓁^α^*has weight at all *z* but the crucial large 𝓁 behavior is controlled by small *z*. For large 𝓁, the integrand in Equation 8 is dominated by *z* of *O*(𝓁^−1^); the integrand thus scales as *z𝓁ρ_A_*(*z*) which implies *ρ_A_*(*z*) *∝ z^−^*^1^*^−α^* for small *z* (see detailed calculation in Appendix A.3). The larger *α* is, the larger the magnitude of lower-order terms.

The amplitude spectrum can be used to define distance-dependent models more generally. By defining non-Gaussian but independent Fourier components whose second moments match *ρ_A_*(*z*), we can generate ensembles of functions which have second order statistics defined by *D*(𝓁) but different higher order statistics. We will discuss this approach further in Section 7.3 when we consider time-dependent landscapes. Until then we will restrict consideration to Gaussian-random landscapes and hence need only to work with the divergence function *D*(𝓁); this greatly simplifies computations and understanding.

### 2.3 Local properties of landscapes

Many previous studies focus on “static” properties of landscapes, that is statistics near genomes not conditioned on prior evolution. One of the most commonly studied static properties is the distribution of local maxima [49, 51]. This metric is often taken as a measure of “ruggedness”, with the aim of getting intuition for how the geometry of a landscape might shape evolution. However the notion of ruggedness is rather ambiguous, and, more generally, the dynamical consequences of such local properties of the geometry are very different in high dimensional spaces than intuition built on low dimensional ones might suggest.

In particular, local static properties depend entirely on short-range correlations and are completely insensitive to longer-range ones. Specifically, in Appendix C.1 we show that the probability of a genome being a local maximum in a Gaussian random landscape is given asymptotically by

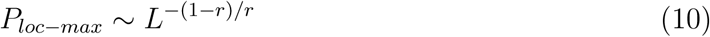

with *r* the correlation coefficient of the fitnesses of the neighbors of a chosen point, which can be computed to be 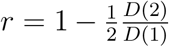 (related to the *γ*-measure from [17] by *r* = (1 *− γ*)*/*2). This probability of being a local maximum is exponentially sensitive to local correlations. For the additive model, *r* = 0 and there is one maximum, for the independent model, 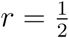 so 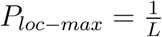, and for the parity function, where every mutation changes the sign of *F* (*g*), *r* = 1 and 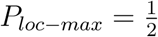.

Note that *P_loc−max_* is independent of correlations in the landscape beyond distance 2: the structure of local maxima has no information about the global statistics. Indeed, the RMF model, *NK* model, and power-law models can each be tuned to have similar numbers of local maxima, but have very different global structure. An RMF model where the additive piece contributes a fraction *p* to the variance has *r* = (1 *− p*)*/*2. An *NK* model with random blocks in the Gaussian limit has 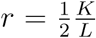, [29]. A power law model with exponent *α* has *r* = 1 *−* 2*^α−^*^1^. Therefore all three models can yield any *r* from 0 to 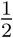. Given any two of these classes of models, one can always find pairs of models with the same local maxima structure, as these only depend on the parameters *p*, 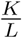 and *α* respectively.

An interesting static property that depends on long-range correlations is the maximum fitness difference, *f*_max(ball)_(𝓁), between a random point and all points in a ball of radius 𝓁 around that point. We are interested in large 𝓁, but 𝓁 ≪ L. For the RMF model, 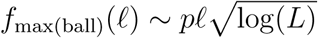 since the maximum is achieved by mutating the 𝓁 sites with the largest additive fitness gains, with an uncertainty of only 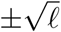 from the independently random piece. For the *NK* model, for *N* = 1 we have approximately *K^𝓁^* independent fitnesses (as there are only *K* sites for which changes lead to fitness differences). This gives us 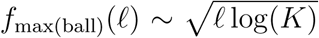 from the maximum of i.i.d. samples from a Gaussian.

For more general correlations, computing *f*_max(ball)_(𝓁) is more complicated; nonetheless, we can proceed in a general way by realizing that most of the genomes in the ball are at the surface (that is at distance 𝓁 from the center). We can then approximate *f*_max(ball)_(𝓁) as the maximum value of these *L^𝓁^* identically distributed Gaussian random variables which have mean 0, variance *D*(𝓁), and covariance 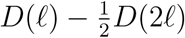 as the points on the surface are typically distance 2𝓁 apart. We thus have, generally,

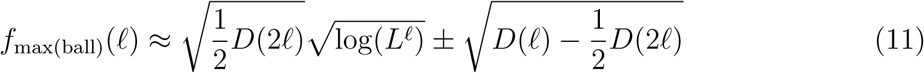

The second, stochastic, term is due to the fluctuations of the average difference at genetic distance 𝓁, while the dominant first term is due to the maximization over all such differences. For the particular case of power-law-correlated landscape, we have, asymptotically

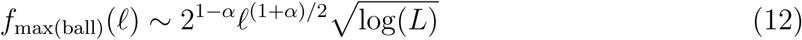

A crude estimate of the maximum fitness on the whole hypercube is given by *f*_max(ball)_(*L*); we expect this to be correct up to polylogs. Therefore the maximum absolute fitness *F ^∗^ ∼ L*^(1+^*^α^*^)^*^/^*^2^ for power law landscapes - see Appendix C.2 for another heuristic derivation.

We see that the behavior of *f*_max(ball)_(𝓁) scales quite differently for the various models — with 𝓁 and sometimes even with *L*. The additive piece of the RMF model has the largest differences between maxima and minima; the *NK* model and other short-range correlated models have the smallest differences as at long-distances they are like independent models. If evolutionary trajectories are able to travel significant distances, then long-range features of the landscape, such as the maximum in a ball, are bound to matter. We will show that it is the long-range structure of *D*(𝓁) that drives evolution, and determines how far uphill evolution can proceed before reaching a local maximum. This discussion should make clear that local structure, such as number of local maxima, gives very incomplete — indeed misleading — information.

## 3 Adaptive walks

“Static” properties of landscapes that focus on behavior near to typical points are misleading: one must take into account conditioning on prior evolution. To do this, we focus on understanding the properties of *random adaptive walks*. An adaptive walk corresponds to evolution in the strong selection weak mutation (SSWM) regime —adaptive mutations are rare, and when they establish they arise to fixation before the next adaptive mutation occurs. By studying the step-by-step dynamics of adaptive walks, particularly over a large number of steps, we will be able to understand directly the feedback between epistasis and evolution. We can then study the properties of genomes and the statistics of available mutations and epistasis among these, conditioned on the evolutionary history. This gives a far better and richer picture of the evolution on rugged landscapes than making qualitative arguments based on local geometric properties.

Specifically, we analyze a clonal population which follows some trajectory *g*(*x*) in genotype space, where *x* indexes the number of mutational steps from the ancestor. At each step, the population randomly samples the space of all possible mutations. For each sampling, the mutation is either rapidly purged, or rapidly fixes. The probability of fixation is some non-decreasing function of the fitness difference *s_g→g′_* = *F* (*g′*) *− F* (*g*) between the mutant *g′* genotype *g* and the current population. Thus the probability of a transition going from *g* to *g′* can be written as

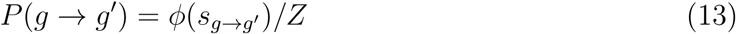

where *ϕ*, the *fixation function*, is the probability of fixation given a mutation, and *Z* is a normalization factor. The dynamics of each step depends only on the local fitness landscape a single mutation away. If *ϕ* allows downhill steps, the dynamics will go on indefinitely. We will focus on walks which must eventually terminate at some local maximum.

There are various choices for *ϕ* that have been studied.

*•* **Random:** the next step is chosen uniformly at random from all uphill directions.
*•* **Greedy:** The best possible step is chosen.
*•* **Reluctant:** The smallest possible uphill step is chosen.
*•* **Natural:** The choice is weighted by some function of the fitness difference to reflect the chances of that mutation fixing. The appropriate choice for moderate sized populations in the strong-selection weak-mutation regime is *ϕ ∝ s*. In large populations, there can be much stronger weighting to large *s* [72].

For the bulk of this paper we will focus, for simplicity, on the *random step* model (i.e with *ϕ*(*s*) the step function): all adaptive mutations have the same probability of fixation, and deleterious mutations are always purged. In Section 5 we will consider both walks which are more greedy and more *abstemious* - tending towards smaller steps.

### 3.1 Adaptive walks on landscapes generated “on the fly”

The simplest way to study adaptive dynamics numerically would be to first draw a particular example of the landscape from the ensemble, and then perform adaptive walks on it. However this would be computationally expensive: for landscapes with non-trivial correlation structure as it would require 2*^L^* random variables. Also, with only numerical work, drawing more general conclusions beyond the particular models studied would be fraught with difficulties.

There have been several recent numerical studies analyzing adaptive walks in the simpler *NK* and RMF models [49, 51, 56, 57]. These works have primarily focused on the lengths of adaptive walks before a local maximum is reached. A phase transition is found in the length of adaptive walks walks as parameters are varied [49, 51, 56]. (We will analytically compute the crossover scale for large, finite *L* in Section 6.) However, the numerical approaches used do not scale well to other parameter regimes, or to other models. The *NK* model, which has complex epistasis, has been simulated only with small blocks (*K* = 8) for which walks of length *∼* 10^3^ have been studied [49, 51]. The RMF models with large *L* are simple enough to run longer (up to walks of length 10^6^) [56]. This is because the additive part of the RMF model only has *L* free parameters, and for the independent fitnesses part, only *XL* random parameters need to be drawn to simulate a walk of length *X*: these can be drawn as the walk progresses.

Distance-dependent landscapes allow us to do something similar for general patterns of epistasis. The Gaussian correlations make computing the local landscape around a single evolutionary trajectory a tractable task, both analytically and computationally. The current distribution of fitness effects can be computed as a conditional distribution based on the parts of the landscape already explored. This enables simulations of dynamics for large *L*, and has the big advantage of making explicit the role of past evolution on the currently accessible fitness landscape. We develop the dynamical approach in the remainder of this section and explore its consequences in Section 4.

The first key observation is that the stochasticity can be divided into two sources. First, the fitnesses of genotypes adjacent to the current genotype *g* need to be generated. This is equivalent to drawing the current distribution of possible single-mutation fitness effects (DFE). The transition probabilities *P* (*g → g′*) can then be computed, and the next step chosen from the resulting distribution. This process is then repeated. Separating the stochasticity in this way ensures that the dynamical step and the DFE step remain statistically independent. The Gaussian distant-dependent statistics will ensure that the conditional DFE only depends on *D*(𝓁) and the already-explored parts of the landscape.

The second simplification is that in the high-dimensional limit, a walk on a hypercube looks like a path on a tree (Figure 2a), with single mutational steps correspond to edges. The tree-like structure comes from the fact that the same site is not likely to be mutated twice, which is generally true if *L* is much larger than the total number, *X*, of mutations accumulated during the evolution. (More precisely we need *X*^2^ *≪ L*, but as the number of sites mutated twice is small as long as *X ≪ L*, we do not expect substantial errors from the tree approximation.) With the statistical symmetry of the landscape in the tree approximation, we can choose to label the genomes, *g*, by the number of mutational steps taken, *x*, with the distance *|g*(*x*) *− g*(*x′*)*|* = *|x − x′|*. We will henceforth label the genomes and other quantities by the step number *x*.

**Figure 2:**
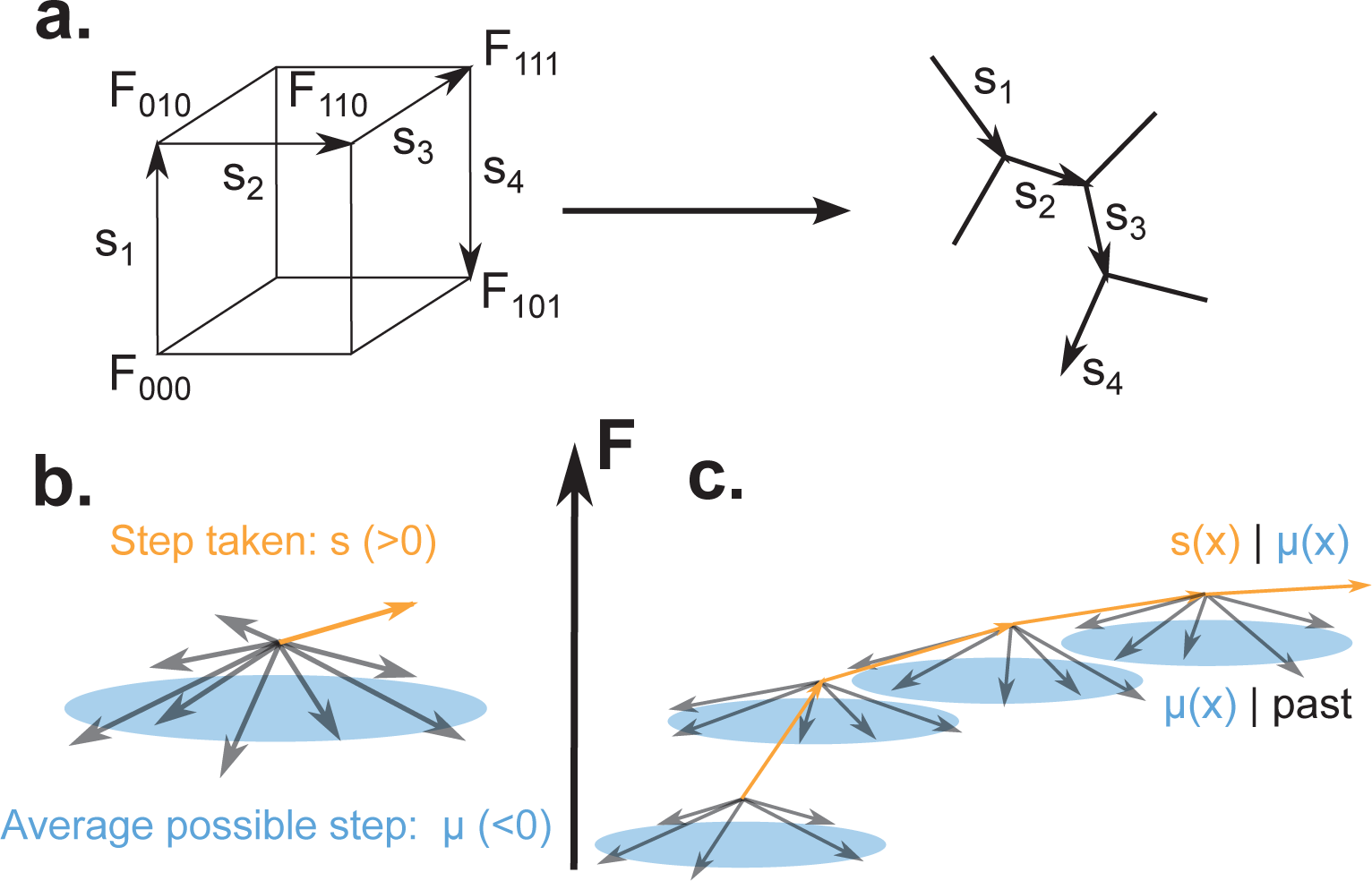
Schematic of adaptive walks on the hypercube. (a) In the high dimensional limit, a single site is mutated at most once, which causes the fitness landscape to look like a tree, with genotypes as nodes, and edges corresponding to fitness differences. (b) At any point, the distribution of single-step fitness gains, *s*, is parameterized by its mean *µ* (blue plane), which is typically negative, while individuals gains have some variance around *µ* (grey arrows). A particular uphill step (orange) is chosen randomly from the possible uphill steps, and taken. (c) Adaptive walks with the right statistics can be found by computing *µ* and then taking a random uphill step *s* from the distribution with mean *µ*; *µ* depends linearly on previously observed *µ* and the previous uphill steps *s*, plus a Gaussian random part. Evolution tends to cause uphill steps to become rarer and rarer by driving typical *µ* to be more and more negative, although the stochastic variations can sometimes make it less negative after a step.

A key property for the adaptive walk is the set of potential *single-step fitness gains* away from the genome at step *x*: {*s_i_*(*x*)}, with *i* labeling the mutated site. The probability distribution of these, the DFE for the possible next steps, has empirical mean

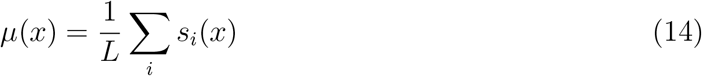

the *average available fitness gain*. Both the potential fitness gains and their empirical mean, *µ*(*x*), are covariate Gaussian. The value of *µ*(*x*) is typically negative during evolution, indicating that beneficial mutations are less common than deleterious ones. Note that we will refer to empirical means within a landscape, such as *µ*(*x*) around a particular genome, as “averages”, while means over the ensemble of landscapes we will refer to as “expectations”. (In statistical mechanics terminology, expectations over the random landscape would be called “quenched averages”.)

The set of single-step fitness increments, *s_i_*(*x*), around any point *x* have variance *D*(1) and identical pairwise correlations 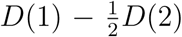. We can thus decompose them into a sum of independent Gaussians *Z* + *z_i_*, where *Z* is the “shared” randomness with variance 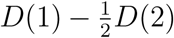 and *z_i_* is the “private” randomness with variance 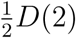 *D*(2). In the limit of large *L*, the average available fitness gain is *µ*(*x*) *≈ Z* (with corrections of order 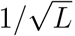), and the conditional gains {*s_i_*(*x*)*|µ*(*x*)} are essentially independent with conditional variance

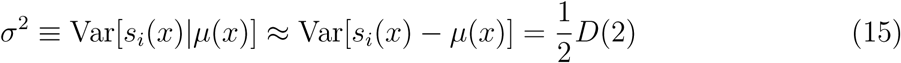

independent of *x*. Therefore, knowing *µ*(*x*) completely determines the DFE around the current genome — the DFE is a Gaussian with mean *µ*(*x*) and variance *σ*^2^ (Figure 2b).

The value of *µ*(*x*), conditioned on previously observed fitnesses, is itself a Gaussian random variable whose statistics can be computed using standard methods which we will review here. Suppose we have a jointly Gaussian collection of vector valued random variables **Z** and **W**, with covariance matrix

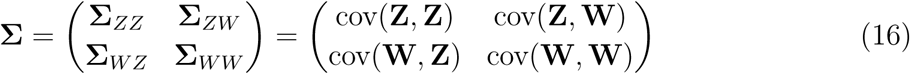

The conditional expectation E[**Z***|***W**] is a linear function of the values of **W**. It can be computed as **K***_Z_***W**, where the linear *response kernel* **K***_Z_* is defined by

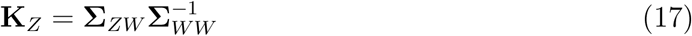

The conditional covariance can be written as

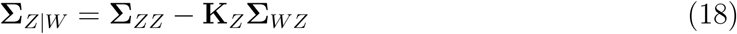

In our case, the **Z** is merely *µ*(*x*), and the **W** is the set of all *µ*(*y*) and *s*(*y*) for *y < x*, where *s*(*y*) are the fitness gains of the single steps actually taken during the evolution (dropping the site subscript *i* as we are in the tree-like limit). Therefore, we can write the conditional expectation as

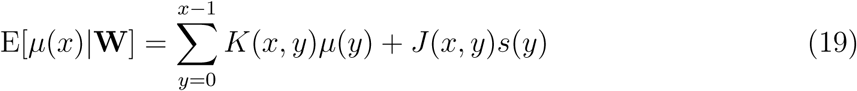

where for ease of later computation we write the response kernels *K*(*x, y*) and *J* (*x, y*) separately. The variance can be computed as:

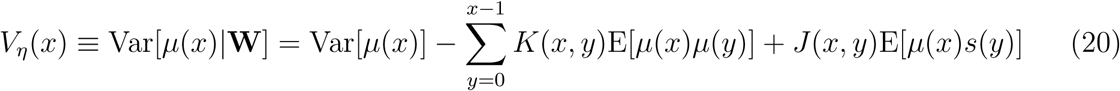

In other words, we can write:

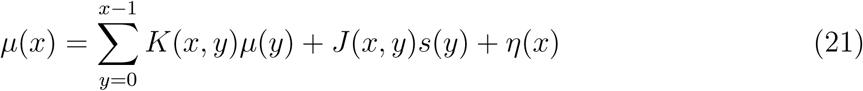

where *η*(*x*), the random part of the average available gain, is Gaussian distributed with variance *V_η_*(*x*).

We again emphasize that this randomness is over the ensemble, conditioned on the structure **W**; *η*(*x*) represents the uncertainty over the distribution of landscapes consistent with the observations **W**, not the stochasticity of the dynamics. Also note that the statistics of *µ*(*x*) depend only on fitness differences and not the absolute values *F* (*x*); therefore they are determined by *D*(𝓁) and are well-defined independent of *L*, as desired.

The structure outlined above implies that, in the large *L* limit, we can simulate the dynamics by computing the statistics of the DFE, choosing a beneficial mutation, and repeating (Figure 2c). More precisely, we use the following dynamical procedure:

1. Generate the conditional expectation of *µ*(*x*) given the previously observed {*µ*(*y*)} and {*s*(*y*)} and the kernels, *J* (*x, y*) and *K*(*x, y*).
2. Choose a value of *µ*(*x*) drawn from Gaussian distribution with its conditional expectation and known variance, *V_η_*(*x*).
3. Generate a step *s*(*x*) from the fixation function *ϕ* applied to a Gaussian with mean *µ*(*x*) and known variance, *σ*^2^.
4. Repeat.

Code which implements this procedure (used for all simulations in this paper) is freely available at: https://github.com/distance-dependent-landscapes/hypercube-walks.

The joint Gaussianity makes computation highly efficient, as the distribution of *µ*(*x*), and hence the distribution of the *s*(*x*), can be computed using simple linear algebra. Computing the response kernel at step *x*, requires a 2*x ×* 2*x* matrix inversion. This can be performed iteratively from the covariances computed at the previous step with cost *O*(*x*^2^). Computing *µ*(*x*) using the response kernels costs only *O*(*x*). The total computation time for *x* ranging from 0 to some *X* thus grows as *X*^3^.

As an explicit example, consider a walk of 2 steps started at a random point (labelled 0). Because of the need to condition the distributions of later steps on the neighborhood of 0, we need to know the mean of the possible first steps, *µ*(0), which has variance *V*_η_(0) = 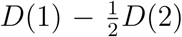. The first step, *s*(0), then takes its value from a Gaussian with this mean, *µ*(0), and variance 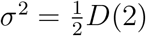 (as per Equation 18). After a step has been taken, the mean of the next available steps, *µ*(1), is the first quantity with non-trivial correlations. It is drawn from a Gaussian with variance *V_η_*(1) and mean *J* (1, 0)*s*(0) + *K*(1, 0)*µ*(0). Direct calculation using Equations 17 and 18 shows that *J* (1, 0) = *−*(*b* + *c*)*/*(*D*(1) *− c*), *K*(1, 0) = (*b* + *c*^2^)*/c*(*D*(1) *− c*) and *V_η_*(1) = [*c*^2^(1 *−* 2*b −* 2*c*) *− b*^2^]*/c*(*D*(1) *− c*), where *b* = *−D*(3)*/*2 + *D*(2) *− D*(1)*/*2 and *c* = *D*(1) = *D*(2)*/*2 (both non-negative). Thus *J* (1, 0) *<* 0 and *K*(1, 0) *>* 0 which means that *µ*(1) is correlated with, but on average more negative than, *µ*(0). The process of computing further *µ*(*x*) and *s*(*x*) then continues as described above.

For all the simulated dynamics in this work, we chose, for simplicity, to approximate the response kernel at all steps by the kernels *µ*(*X, y*) and *s*(*X, y*) at the final step *X*. Likewise, we used *V_η_*(*X*) to approximate *V_η_*(*x*) throughout the walk. This gives us the same computational complexity as the iterative inversion method. Numerical results show that *V_η_*(*x*) saturates quickly to the long time value *V_η_*(*∞*) (as expected) and the analytical results in Section 3.2 suggest that the key parts of the response kernels converge as well. This approximation gives largest errors at the start of the walk, but, as we will see in Section 3.3, the long-term dynamics depends only weakly on early times.

The response kernels depend only on the geometry of the path. If the walk geometry is known in advance (as it is for a population in the strong selection weak mutation regime on which we focus), the response kernels can be computed once and then used for multiple simulations on independent but statistically identical landscapes. Therefore *T* simulations of length *X* can be achieved in *O*(*X*^3^ + *TX*^2^) time in the large *L* limit. The generation of the landscape “on the fly” has removed any dependence of the computational complexity on the dimension of the genotype space.

The form of Equation 21 assumes a walk along a single path, with the geometry of a straight line. However, the framework extends to trajectories with more complicated topologies as well. In particular, there is a computationally straightforward extension to the case of multiply branching paths on a single landscape. The response kernels are computed via Equation 17 to obtain *µ*(*x*) at the end of any branch, and the potential single-step fitness gains *s_i_*(*x*) are once again independent when conditioned on *µ*(*x*).

### 3.2 Single-path response kernels

In addition to enabling efficient computation, the response kernels can be used to gain a qualitative and quantitative understanding of how the dynamics of adaptive walks depend on the past. We will take advantage of this structure to analytically approximate the statistics of fitness trajectories for a variety of landscapes, and obtain exact results for some simple cases.

The response kernels encode exactly the effect that the past evolution has on evolution in the near future. Because this depends on the *trajectories* of *µ*(*x*) and *s*(*x*), and not simply their most recent or average values, the detailed dynamics of evolution determines the current DFE. The Gaussian nature of the landscape means that the present depends linearly on the past; however the evolutionary dynamics causes non-linear feedback via the selection of the uphill step from the DFE. This interplay between past and present is what gives rise to the complexities of evolutionary dynamics even in these Gaussian-distributed landscapes.

As we shall show explicitly, the response of *µ*(*x*) to *s*(*y*) for *y < x* (via the kernel *J* (*x, y*)), tends to be negative; the current possible steps are anticorrelated with previous ones taken. This negative feedback makes it harder and harder to find uphill steps as the evolution continues. The response of *µ*(*x*) to *µ*(*y*) (via the kernel *K*(*x, y*)) is somewhat subtle, but its net effect is to make *µ*(*x*) correlated with earlier *µ*(*y*).

#### 3.2.1 Exponentially-correlated landscapes

We begin by analyzing the case of exponential correlations:

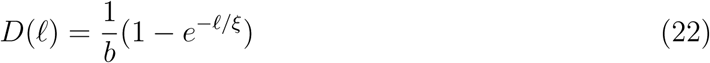

where *b* = (1 *− e^−^*^1^*^/ξ^*) normalizes the family so that random single mutants have identical statistics (*D*(1) = 1) for any *ξ*.

For this model, the response structure is easy to compute in terms of the absolute fitness values *F* (*x*). A direct calculation (detailed in Appendix B.3) shows that we have

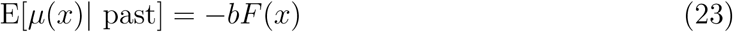

The dynamics is “memoryless” (Markovian) with regards to the fitnesses — only the last observed fitness matters. This relationship is in fact exact; that is, we can write

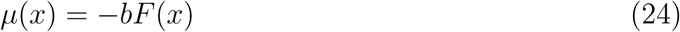

without the conditional expectations. This is a very special property of the exponential landscape (and indeed, completely characterizes it for *b ∈* [0, 1]). The relationship is most clear in the independent limit where *b* = 1. Here in the large *L* limit we have

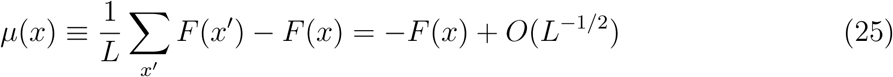

where the *x′* are genomes one mutation away from *x*, whose average is zero due to the mutual independence of all fitnesses

Equation 24, along with the definition of the step taken, *s*(*x*) = *F* (*x* +1) *− F* (*x*), implies that the joint covariance matrix of the *µ*(*x*) and *s*(*x*) is singular. The response kernel is degenerate to degree *x*. We can take advantage of the freedom of choice to make the response kernels take simple forms; two possible choices are

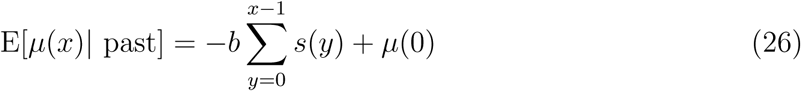

or

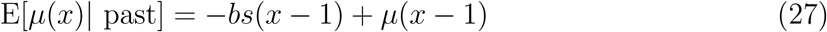

The negative correlation with the previous fitness will drive the typical *µ*(*x*) to be more and more negative during evolution. Since the variance of *µ*(*x*) is constant, this leads to the typical DFE having fewer and fewer beneficial mutations, slowing down the rate of fitness gain, as expected. In Section 3.3, we will compute the dynamics for exponential correlations explicitly.

From Equation 15 the variance of the available *s*(*x*) given *µ*(*x*) is

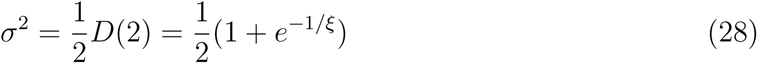

which ranges from 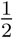 (independent model) to 1 (additive model) for increasing *ξ*.

#### 3.2.2 Power-law-correlated landscapes and integral approximation

To understand the behavior of long walks — in which we are primarily interested — it is instructive to approximate the discrete response equation by the integral equation:

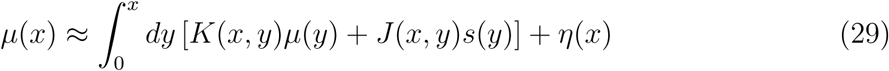

where we abuse notation and use *J* and *K* for the continuous analogues of the discrete response kernels. Now *η*(*x*) is the random part of *µ*(*x*), a Gaussian random variable with autocorrelation

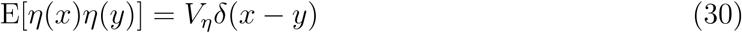

As we are interested in the long-walk behavior, we use the asymptotic value, 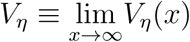, at which the conditional variance saturates.

In general, the kernels will depend on both *x* and *y*. But for *x − y ≪ x*, we anticipate that they will approximately be functions of just the distance, *x − y*. In this limit, the integrals become convolutions and Fourier transforms can be used. For the case of power law divergence functions, Wiener-Hopf analysis (Appendix B) can be used in this *x − y ≪ x* regime, to obtain *J* (*x, y*) *≈ −c_J_* (*x − y*)*^−^*^(1^*^−ν^*^)^ and *K*(*x, y*) *≈ δ*(*x − y*)+ *c_K_P*(*x − y*)*^−^*^(1+^*^ν^*^)^ with the exponent *ν* = (1 *− α*)*/*2 and 𝒫 denoting the principal part (so that integrals over *K* have no contribution from the divergence at *x − y →* 0). Here *c_J_*and *c_K_* are positive coefficients that can be written in terms of Beta functions and combinations of *V_η_* and *σ*^2^.

Armed with their scaling forms, and the structure of the singular integral equations that have to be solved, one can guess and check the exact kernels (valid when *x*, *y* and *x − y* are all large):

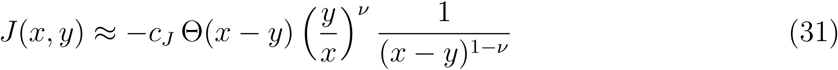

and

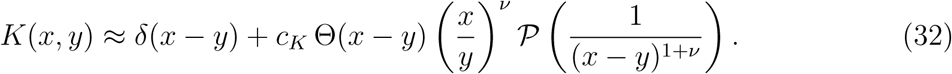

where Θ(*z* > 0) = 1 and Θ(*z* < 0) = 0.

The responses are of opposite signs, and the *J* drops off less sharply than the *K*. However, the sign of *K* is misleading due to the principal part. If the convolution with the past is done by parts, then we get

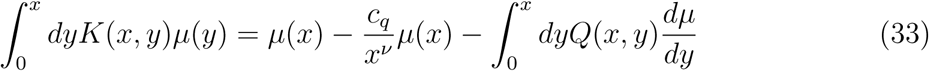

with positive coefficient *c_q_* and the kernel of 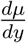 defined by

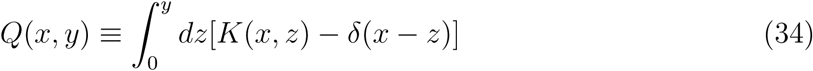

This form of the response shows that *µ*(*x*) is anticorrelated with both *s*(*y*) and 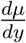 of the past.

The analytically predicted asymptotic scaling forms of the response kernels, agree well with the numerically computed kernels. In Figure 3, we plot the numerically computed *−J* and *K* for a variety of walk lengths *x*. The predicted scaling forms are plotted as dashed lines. We choose the coefficients *c_J_* and *c_K_* to match the forms at *x − y* = 1 in lieu of computing these numerically. We see that for large *x*, the response kernels drop off with the predicted power law form until *x − y* is large (*y* small). The predicted asymptotic forms also capture the behavior at small *y*, where the response kernel is non-monotonic due to the dependence on *x/y*. The *J* response kernel is closest to its asymptotic form for *α* = 1 and the *K* is closest for *α* = 0 (detailed in Appendix B.2); these are expected, as the corrections to the asymptotic forms go as 1*/x^α^*and 1*/x*^1^*^−α^* respectively.

**Figure 3:**
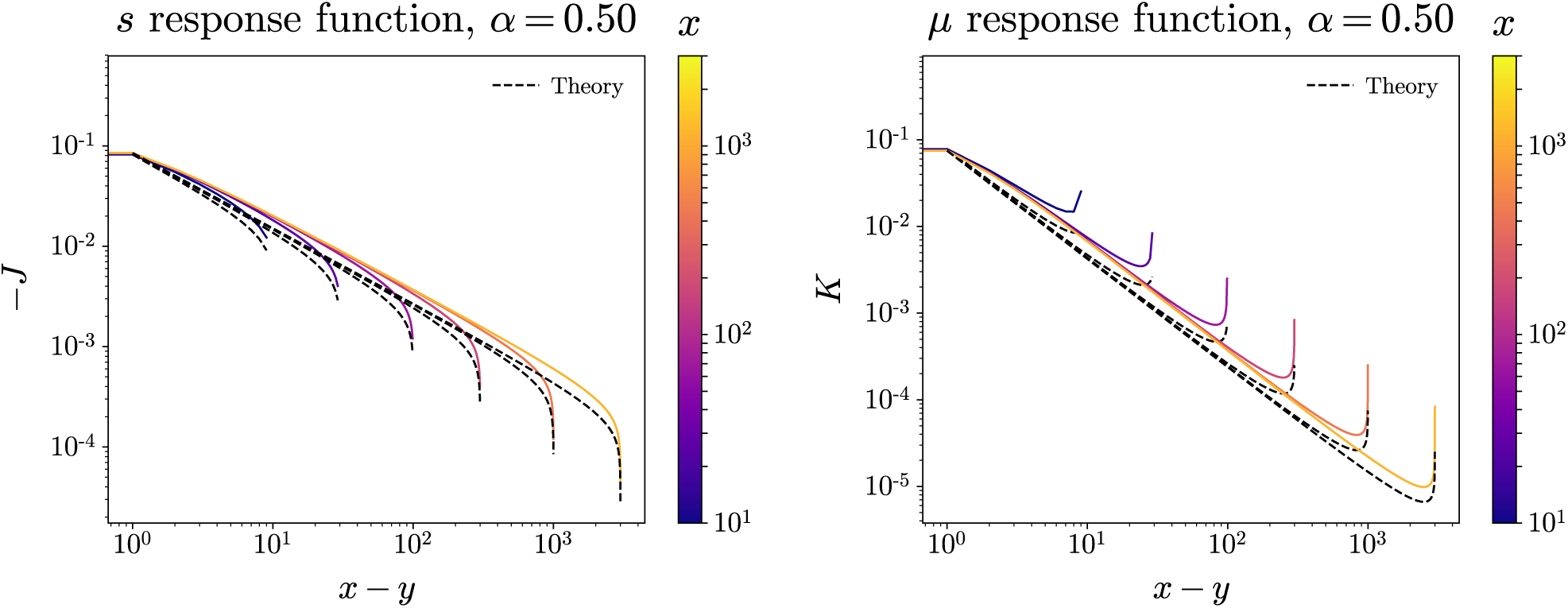
Log-log plots of *−J* and *K* against *x − y* for various *x*, *α* = 0.5. response kernels explicitly encode dependence of evolution on the past. Intermediate power law scaling regime makes long range of evolutionary history important. Theoretical form (dashed lines) rescaled to match numerical form at *x − y* = 1. Theory captures intermediate power law scaling regime, as well as non-monotonic behavior for small *y*.

From the form of the response kernel, we can already see a qualitative difference between the power-law and exponential cases; the conditional probabilities in general depend on the whole fitness-trajectory of the past evolution, not just the most recent fitness. This dependence on the long-ago past is crucial for shaping the long-term dynamics of random adaptive walks.

### 3.3 Typical fitness trajectories of adaptive walks

Given an analytical form for the long-distance response kernel, we can obtain analytical understanding of the typical fitness trajectories of random adaptive walks. The response kernels give an integral equation for the conditional expectation (across adaptive walks over the whole ensemble of landscapes) of *µ*(*x*) in terms of past *µ*(*y*) and *s*(*y*), and then the value of *µ*(*x*) determines the distribution of *s*(*x*). Since the evolution-conditioned E[*µ*(*x*)*|*past] is linear in past values, we can use the response kernels to get a self-consistent set of equations for the typical trajectories of *µ*(*x*) and *s*(*x*).

We define the *fitness gain* of a trajectory as a function of the number of steps, *x*, along it as

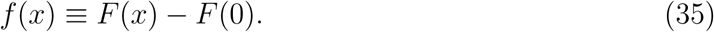

Due to the distance-dependent statistics, *f* (*x*) should, for walks with *x ≪ L*, be independent of *L*. We start with simple examples where the answers are known from simple methods, and work up to the more complicated cases.

#### 3.3.1 Simple landscapes

The simplest case is the additive model: the fitness steps are uncorrelated, so both *K* and *J* are 0. Therefore *µ*(*x*) is always 0 on average, and uphill steps are taken from the same distribution. The total fitness gain, *f* (*x*), typically grows linearly in *x*, with variance around the average fitness trajectory, 〈*δf* ^2^〉, also growing linearly. The 〈*·*〉 notation indicates randomness due to stochasticity of the dynamics, rather than (possibly conditional) averages E[*·*] over the ensemble.

The other simple case is the independent model. Since the fitnesses are all independent, there is a simple relationship between the average available fitness gain and the (absolute) fitness: *µ*(*x*) = *−F* (*x*) for any *x* in the large *L* limit, with no additional variation (i.e. *V_η_* = 0). The difficulty of finding uphill steps is directly related to the current fitness. (Note that the absolute fitnesses are well defined in the independent model since *D*(𝓁) is bounded, as per the discussion in Section 2.1).

To proceed, we need to understand the typical uphill steps actually taken, given *µ*(*x*). The distribution of possible steps is just determined by *µ*(*x*) and *D*(2) for any *D*(𝓁) (with *D*(1) = 1). For large negative *µ*(*x*) (*−µ*(*x*) *≫ σ*), the distribution of uphill steps is distributed approximately exponentially with average value

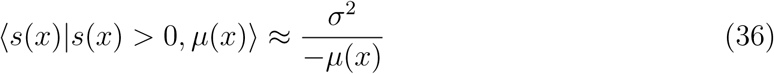

where, as before, 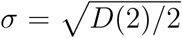 is the — history independent — standard deviation of the potential single-step fitness gains {*s_i_*(*x*)} given *µ*(*x*). Using the above expression, we have, for the independent fitnesses model, an approximate differential equation for *F*:

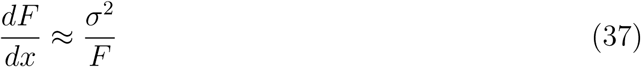

which integrates to *F* (*x*) *∼ x*^1^*^/^*^2^ for large *x*. The fitness in the independent model thus increases sublinearly, but still as a power law in the number of steps. For *x≫*1, the effects of the initial fitness, *F* (0), are negligible (yielding corrections to *F* (*x*) of order 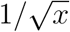). Thus whether or not we condition over the initial fitness, the long-distance behavior is essentially the same.

For landscapes with exponentially decaying correlations, we can proceed similarly, since far-apart regions of the landscape are approximately independent. Normalizing *D*(1) = 1 as before, Equation 24 gives

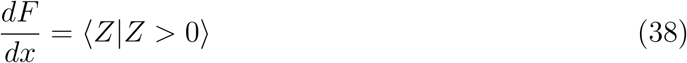

where *Z* is normally distributed with expectation *−F* (*x*)*/b* and variance 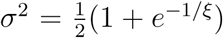 as calculated previously.

As long as *F* (*x*)*/b ≪σ* (equivalent to *F ≪ ξσ* for large *ξ*), the dynamics is like the additive model with 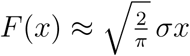. The correlations are too weak to inuence the dynamics and F increases roughly linearly until *F ∼ ξσ*, where the correlations start to matter. For *F ≫ ξσ*, the large *µ* approximation holds and we have

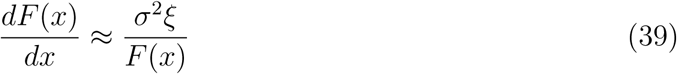

which gives us the approximate trajectory

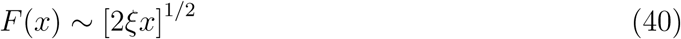

at long times for large *ξ*. The long-distance behavior is thus like the independent-fitness model as could have been anticipated since fitnesses that are much more than *ξ* apart look roughly independent.

#### 3.3.2 Power-law correlated landscapes

Analyzing adaptive paths on power law landscapes is more complicated, because past evolution matters at all stages due to the power laws in the response kernels. Figure 4 shows a typical example of the evolutionary dynamics. The uphill steps get smaller and smaller as the average, *µ*(*x*), of the DFE decreases.

**Figure 4:**
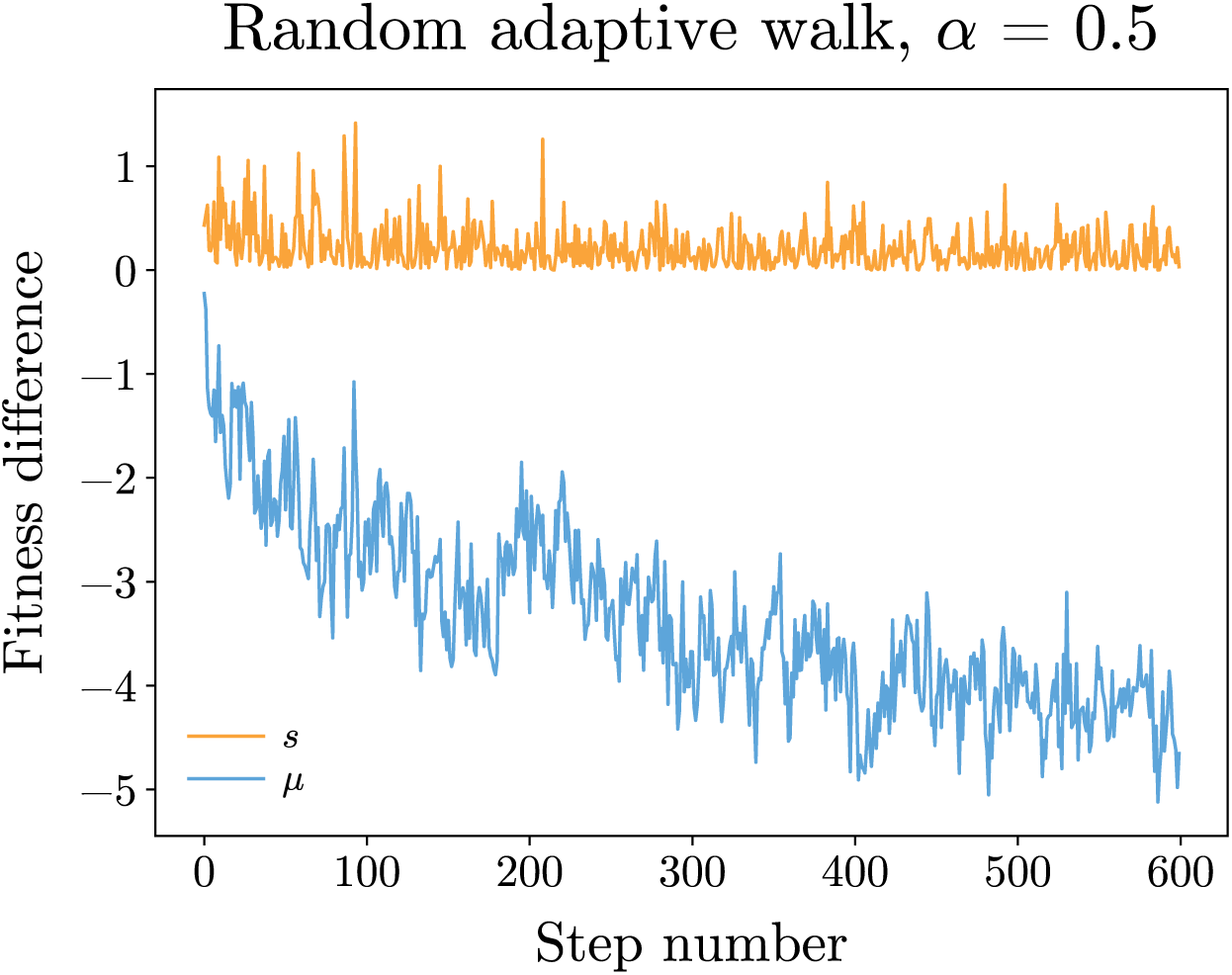
Properties of typical random adaptive walk on a power-law-correlated landscape, with *α* = 0.5. Fitness gains *s* decrease, on average, as a power of the number of steps, while the average available fitness gain, *µ*, is negative with *|µ|* increasing as a power of the number of steps.

Regardless, we can use the power law forms of the response kernels to self-consistently solve for the dynamics. For power-law correlated landscapes, we must deal with fitness differences only since *F* (0) depends on *L* (*D*(𝓁) is unbounded). We will solve for the dynamics of the fitness gain *f* (*x*) by making a power-law ansatz for the typical behavior

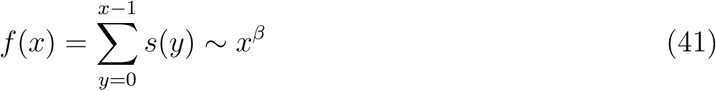

and by solving for the scaling forms of *µ*(*x*) and *s*(*x*). We conjecture that

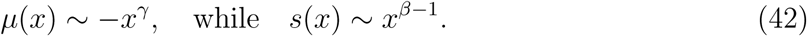

Evaluating the convolutions in Equation 29 (with short-hand notation *∗* for the integrals over the past) using Equation 33 and neglecting the noise term (justified *a posteriori*), we have 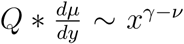 and *J ∗ s ∼ −x^ν^*^+^*^β−^*^1^, where *ν* = (1 *− α*)*/*2 *<* 1. The neglected terms in Equation 33 are much smaller than than 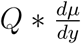 for large *x*; therefore the *Q* term must nearly cancel the *J ∗ s* term. Thus both those pieces must scale the same way with *x*, so we conclude that 1 *− β* + *γ* = 2*ν* = 1 *− α*.

To get a second equation relating the exponents, we consider the dynamics. Once *−µ* becomes large — which taking random beneficial mutations it will be after only a few steps — then *s* must be selected from the tail of the Gaussian distribution. Equation 36 then typically holds and *s*(*x*) *∼ σ*^2^*/*[*−µ*(*x*)]. Combining with the Ansatzes in Equation 42, we have 1 *− β* = *γ*.

Combining with the first equation for the exponents, we can now solve for *β* and *γ* and conclude that

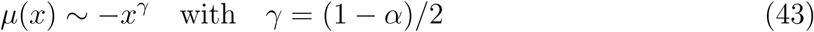

and

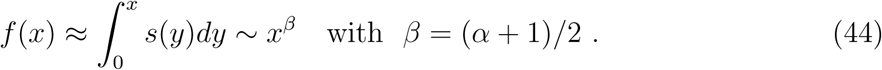

The exponents interpolate linearly between the extreme cases of the additive (*α* = 1) and independent fitnesses (*α* = 0) models.

These analytic predictions for power-law correlated landscapes are supported by the numerics. Figure 5 shows the mean and standard deviation of the trajectories of 300 simulations of adaptive walks on different instantiations of the fitness landscape for *α* = 0.5 (left column). As expected, *−µ* is constantly increasing, while *s* decreases accordingly. In the right column, we show log-log plots of *µ*(*x*), *s*(*x*), and *f* (*x*) for various *α*. Since *−µ*(*x*) increases more slowly for larger *α*, *f* (*x*) grows more quickly as *α* increases. Figure 6 shows the scalings of the log derivatives of the mean and variance of each set of trajectories (computed numerically after 100 steps, with Gaussian smoothing of width 5). The estimated power law exponent, *γ*, of *µ* deviates from theory as *α* goes to 1; however, this is also where the response kernel *K* has corrections of relative order 1*/x*^1^*^−α^* (as well as 1*/x^α^*), and *µ* is only slowly increasing. In all, the scalings from the numerics appear to agree well with the analytic predictions.

**Figure 5:**
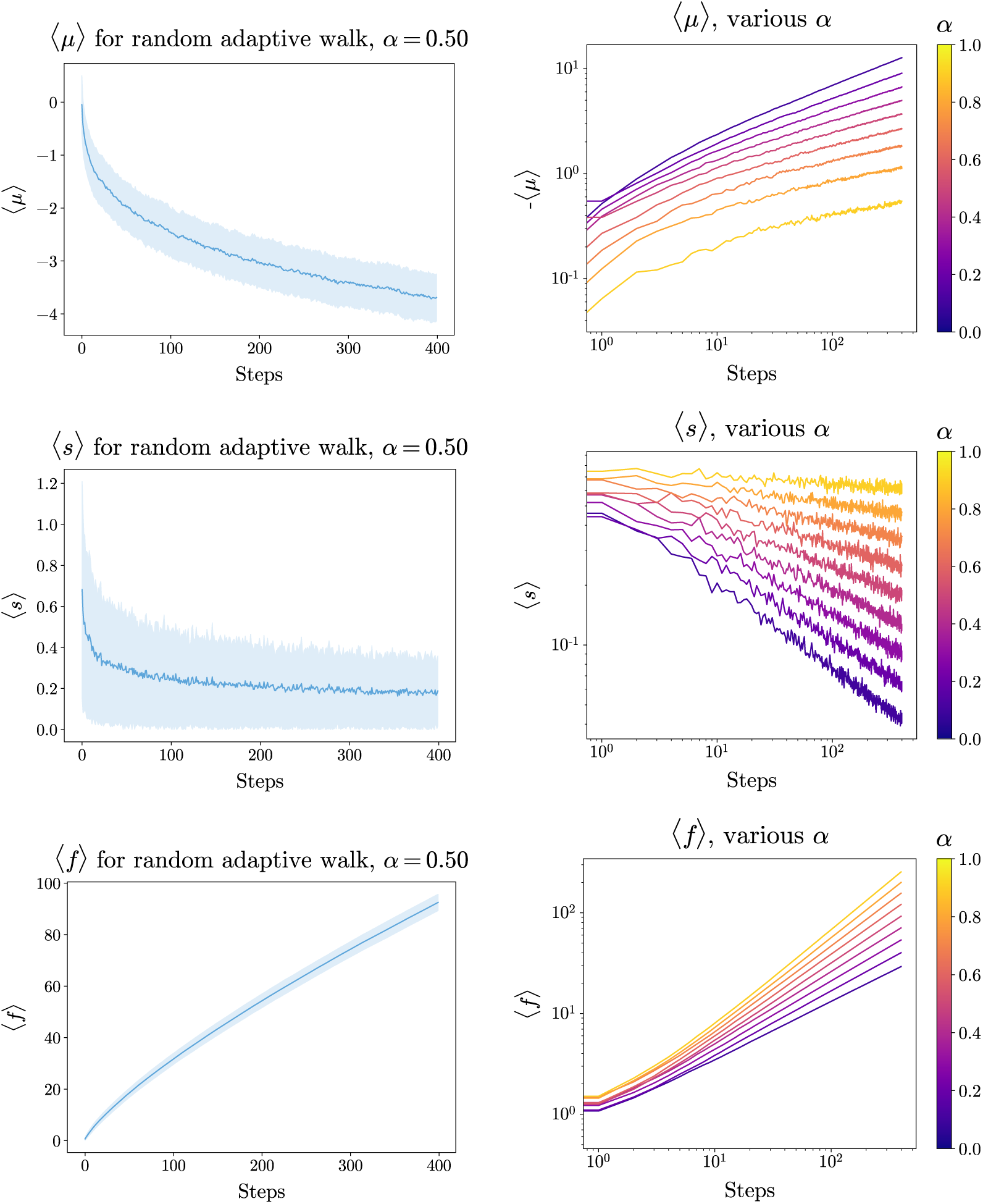
Mean and standard deviation of *µ*(*x*), *s*(*x*), and *f* (*x*) for adaptive walks with *α* = 0.5 averaged over 300 trajectories (left column). Log-log of averaged trajectories for various *α* (right column). *µ*(*x*) is negative and decreases more slowly for larger *α*. This slower decrease leads to larger *s*(*x*), and therefore to larger *f* (*x*). Relative variability of *µ*(*x*) and *f* (*x*) decrease with *x*, while that of *s*(*x*) does not.

**Figure 6:**
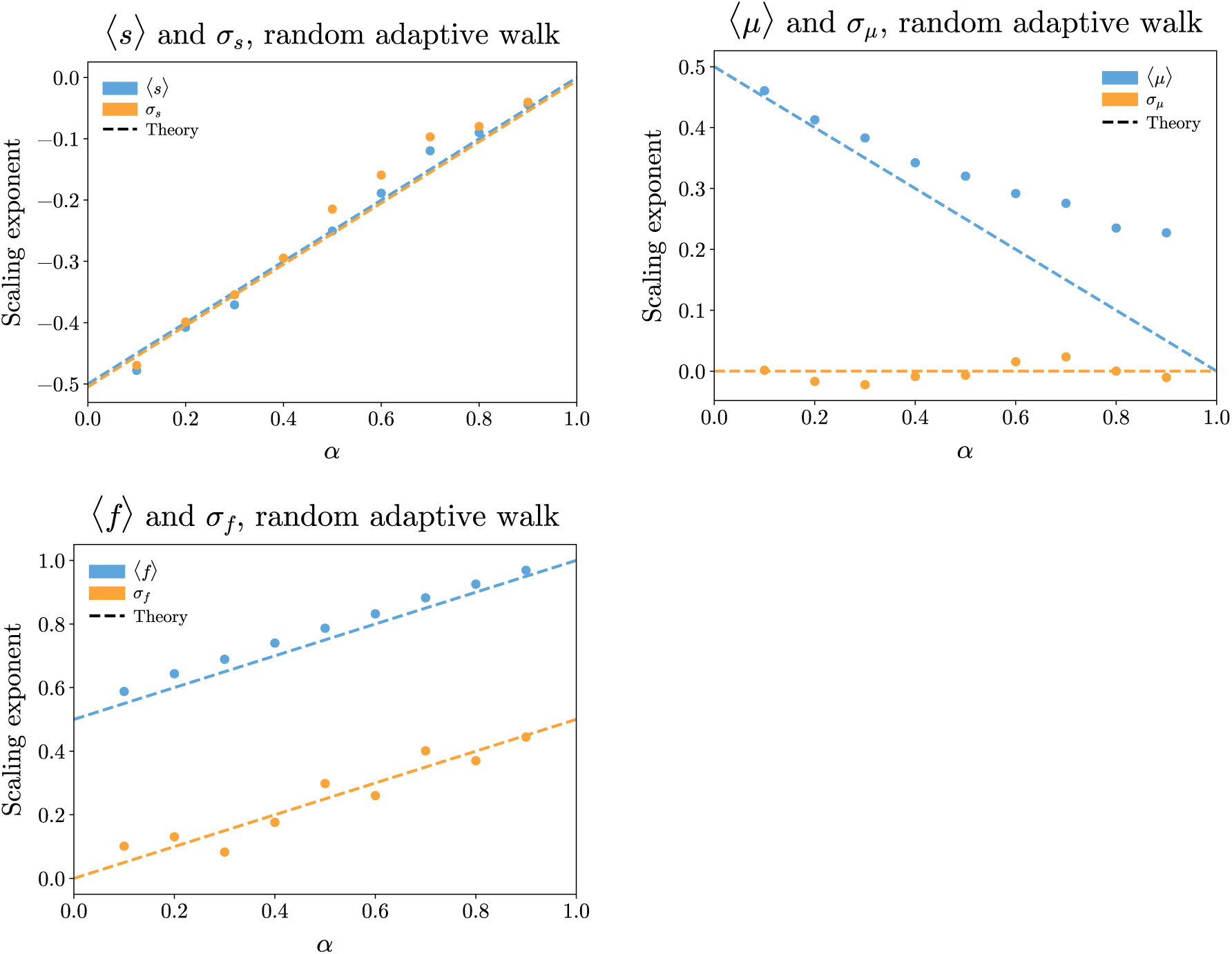
Logarithmic derivatives *d* log(*quantity*)*/d* log(*x*) of trajectories for *s*, *µ*, and *f*. Derivatives computed numerically from trajectories in Figure 5 around *x* = 100. Derivatives of average values in blue, derivatives of standard deviations in orange. All quantities roughly follow power law trajectories. Exponents of *s* and *f* increase linearly with *α*, while *µ* decreases. Simulations (dots) match theory (dashed lines) well for *s* for all *α* and for *µ* for low *α*. Deviations are due in part to finite walk-length effects.

The simulations also illustrate the scalings of the variations about the mean of the various quantities. The variance of *µ* goes to a constant while *|µ*(*x*)*|* increases. The variance of *s* is proportional to 〈*s*〉^2^, as one should expect for an exponentially distributed random variable; this is roughly correct even though the variations in *µ* make the distribution of *s* across realizations is not simply exponential, the standard deviation is still proportional to the expectation. The variance of the fitness scales as 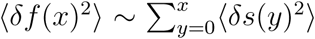, as would occur if the uphill steps were uncorrelated. (These statistics are for the dynamics across different landscapes; in Section 4.3, we discuss variability due to dynamics on a single landscape.) The relative variances of both *f* (*x*) and *µ*(*x*) go to 0 for large *x*, and therefore the “typical” trajectory is well described by the average trajectory.

The key feature of the dynamics is that the “global” epistatic information — the long-distance correlations in fitness – determine the rate and predictability of evolution, as opposed to local, short-range statistics (which are the ones responsible for things like the number of local maxima). The evolutionary trajectories of different landscapes look similar if their statistics are similar. We will discuss the relationship of fitness trajectories within the same landscape in more detail in Section 4.3.

## 4 Properties of adaptive walks

The analytical framework introduced in the previous section allows us to understand many features of the evolutionary dynamics quantitatively. The explicit dependence on past evolution means that the long-range statistical structure of the landscape is important for understanding evolutionary trajectories. In particular, evolution drives populations to highly non-generic places on the fitness landscape. We highlight some of the important properties of the dynamics and the evolution-conditioned fitness landscape here.

### 4.1 Finite ***L*** and reaching a local maximum

The results discussed thus far are for the limit of infinite *L*: we have assumed that there are enough beneficial mutations at any step that the distribution of fitness effects is essentially deterministic given *µ*(*x*), and many uphill steps are always available, even when these constitute a very small fraction of the total of *L* potential mutations. A crucial consequence of finite *L* is the tendency to eventually run out of beneficial mutations: this will occur when the fraction of mutations that are beneficial decreases to *O*(1*/L*). We can predict the magnitude of the total fitness gained, *f*_max_, before a local maximum is reached. This can be calculated directly for the independent fitnesses and independent steps (additive model) cases. For power-law correlated landscapes, we can use our large-distance-scale approximations to find how *f*_max_ scales with genome size, *L*.

For the additive model, each site contributes independently to the fitness. Thus the global maximum can be reached and *f*_max_ *∝ L*. For the independent fitnesses model, at every step, we are presented with *L* independent choices of new fitnesses, *F*. A large fitness is unlikely to be found when the probability of any individual neighboring fitness being greater than the current fitness is *< O*(1*/L*). This implies that the absolute fitness *F_max_* will scale as 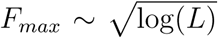, and since a typical starting point has *F* (0) *∼ O*(1), 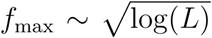 as well. Note that this *f*_max_ is a consequence of the Gaussian tail of the fitness distribution: with longer-tailed distributions, as we discuss in Section 6, the fitness can increase further.

The maximum fitness reached can be estimated more generally, similarly to the independent fitnesses model: simply follow the deterministic dynamics until a value is reached such that there are only *O*(1) available uphill steps. This occurs after a number of mutations *X* where *P* (*s*(*X*) *>* 0*|µ*(*X*)) *∼ O*(*L^−^*^1^). With our Gaussian random landscapes, this condition is 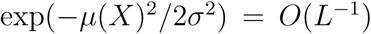, i.e. when 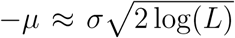. Until then, with high probability there are available uphill steps. Once the form of *µ*(*x*) is known, we can solve for *X*, and estimate *f*_max_ *∼ f* (*X*).

In Section 3.3, we found the scaling forms of *µ*(*x*) and *f* (*x*) for exponentially correlated landscapes and power-law correlated landscapes, allowing us to solve for *f*_max_ for each. For exponential landscapes the cumulative fitness gain saturates at *f*_max_ *∼ σ*(1 *− e^−^*^1^*^/ξ^*)*^−^*^1^*^/^*^2^[log(*L*)]^1^*^/^*^2^. For *ξ ≫* 1, this is approximately *σξ*^1^*^/^*^2^[log(*L*)]^1^*^/^*^2^, the same scaling with *L* as the independent case. (Of course, if *ξ* = *O*(*L*), then *f*_max_ *∝ L*.)

In contrast, for power-law correlated landscapes we have

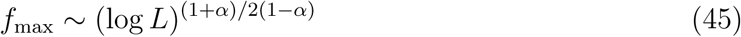

This is still logarithmic in *L*, but can be many times higher than for the independent case (which corresponds to *α* = 0).

How good are the maxima which adaptive walks reach? Specifically, one can ask: how large is *f*_max_ compared to the maximum fitness *f*_max(ball)_ of any genome within the same distance from the starting point as the length of the walk, *x*? Using Equation 11, we can compute *f*_max(ball)_ for a ball of radius *x*. For power-law landscapes, 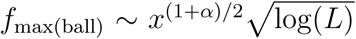. As the adaptive walk reaches *x* = (log *L*)^1^*^/^*^1^*^−α^* and *f* (*x*) *∼ x*^(1+^*^α/^*^2)^, both *f*_max_ and *f*_max(ball)_ are powers of log(*L*), but the latter has an extra factor of 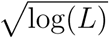.

Another way to quantify how effective adaptive walks are is to compare their fitness increase to *f*_max,up_, the maximal fitness gain attainable via *any* adaptive walk. The question of whether or not *f*_max,up_ reaches the maximal possible fitness is called *accessibility percolation* [29, 50]. For the independent fitnesses case, explicit computations (Appendix C.2) yield *f*_max,up_ *∼ L*^1^*^/^*^4^. This is a factor of *L*^1^*^/^*^4^ smaller than the maximum fitness *F ^∗^ ∼ L*^1^*^/^*^2^ of any genotype on the hypercube. Therefore the largest possible fitnesses are inaccessible via purely uphill evolution. This is consistent with previous proofs that the probability that there exists an uphill path to the maximal fitness goes to zero in the independent fitnesses model (unless the initial fitness is anomalously small) [28].

In contrast, the RMF model behaves like the additive model, with probability one of having an uphill path to the maximal possible fitness. For landscapes without an additive piece, results are likely similar to the independent fitnesses case. As we will show in Section 5, even alternate weighting schemes to choose better adaptive walks do not enable them to reach anywhere near *F ^∗^* in power law models.

All the above results underscore that understanding typical behaviors of adaptive walks is very different than understanding either “typical” local or large-scale geometric properties of the fitness landscape.

### 4.2 Comparisons with typical paths to uphill points

One way to quantify the “non-generic” nature of the evolutionary dynamics is to compare adaptive trajectories to ‘typical” paths that reach the same fitness gain in the same number of steps. We thus must ask: on a power-law correlated landscape, what does a typical random walk of length *x* and fitness gain *x*^(^*^α^*^+1)^*^/^*^2^ look like?

We can use the covariances calculated in Appendix A.1 to find the average trajectory conditioned on the endpoints using formulae for conditional expectations of Gaussian random variables: For power-law correlated landscapes, we find

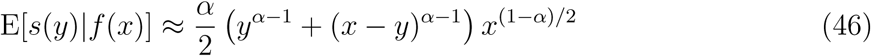

(Note that here the conditioning is done simply over the ensemble of landscapes.) In contrast to adaptive trajectories, the typical trajectories are symmetric, steepest at both ends. The average step size is at most *x*^(1^*^−α^*^)^*^/^*^2^, but decreases greatly in the middle to *x^−^*^(1^*^−α^*^)^*^/^*^2^.

Sample trajectories for *α* = 0.5 are shown in Figure 7. The typical conditioned trajectories tend to be very lucky at the beginning and end (with high slopes there), and increase only very slowly in the middle. This is in contrast to the adaptive walks, which are typical at the beginning, and then gradually slow down but continue to gain fitness. In the middle portion of the typical conditioned random walk the average step size is *x^−^*^(1^*^−α^*^)^*^/^*^2^, but the variance of the step sizes is *O*(1), which means that the typical conditioned path does not go systematically uphill. Indeed, the probability of a step in the middle being deleterious is only slightly less than 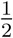. Conversely, the late steps are much larger than those of adaptive walks.

**Figure 7:**
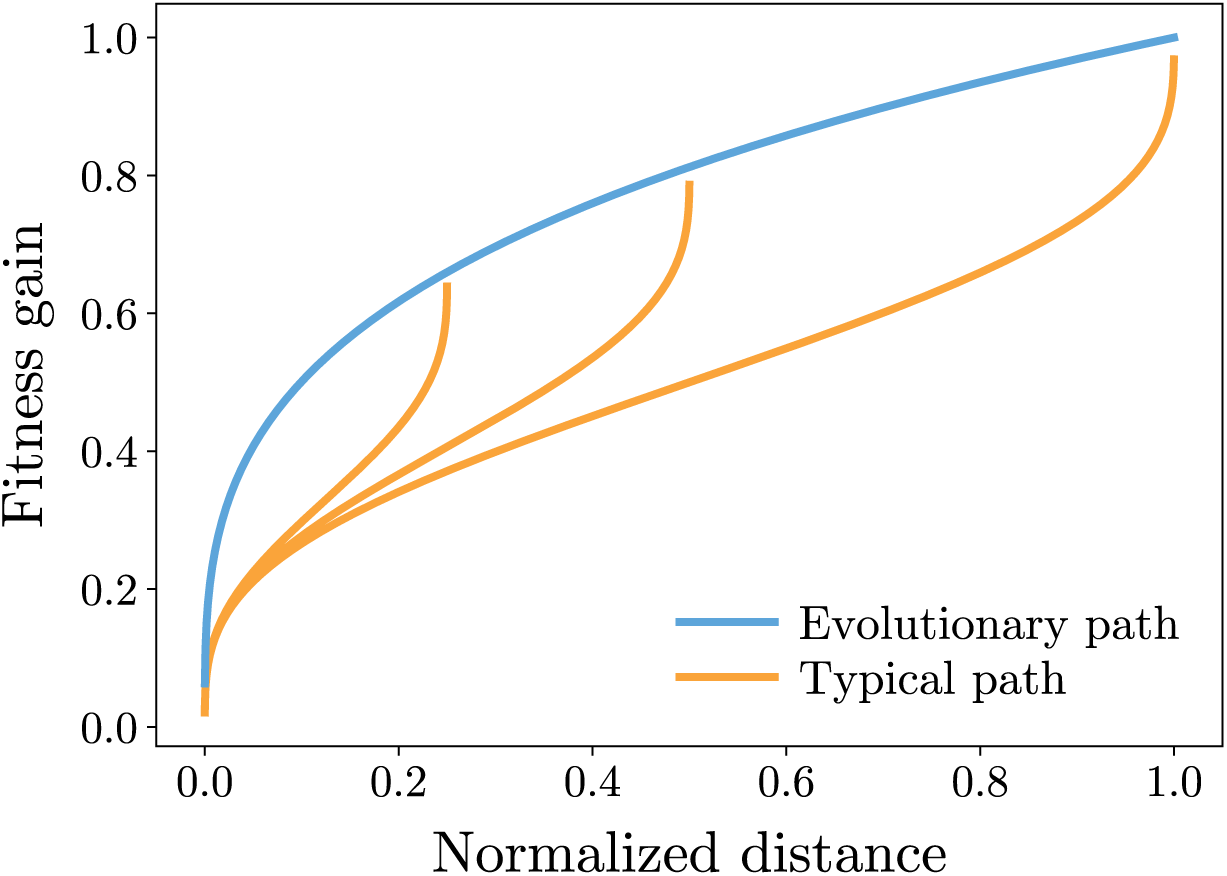
Average evolutionary trajectory (for *α* = 0.5) compared to typical trajectories of random walks on the landscape constrained to end at a distance and fitness on the evolutionary trajectory. Evolution finds a steady path uphill, while typical trajectories with the same fitness gain are lucky at the start and end of the path.

Despite these differences, the neighborhood of the endpoint of the typical conditioned walk has *µ*(*x*) *∝ −x*^(1^*^−α^*^)^*^/^*^2^ just like the adaptive walk. However, even though the scalings of the *µ*(*x*) at the end of the typical walk and the adaptive walk are the same, the coefficients are not; the typical walk has a larger negative *µ*(*x*) by a multiplicative factor. Since the fraction of mutations that are adaptive falls off as a Gaussian in *µ*(*x*), this difference can be very important for future dynamics — adaptive walks are more likely to continue further before getting stuck.

More generally, the unexplored local neighborhood around the endpoint of the two types of walk looks different. For the typical conditioned walk, the mean available step a distance 𝓁 away from the endpoint, *µ*(*x*+𝓁) increases towards zero smoothly as *−𝓁^α−^*^1^*µ*(*x*). However, for an adaptive walk, the magnitude of *µ*(*x* + 𝓁), decreases sharply from *µ*(*x*) before a smoother decrease towards zero. Characterizing in detail the landscape as a function of 𝓁 for the adaptive walk is more complicated than for the typical walk; although our methods could be generalized to do so. Regardless, the behavior in the immediate vicinity of the end of a past evolutionary suggests that it is easier to go further uphill than it would be from a typical conditioned path with the same overall fitness gain.

### 4.3 Variability of evolutionary outcomes

The variances computed in Section 3.3 were computed across the entire ensemble of fitness landscapes, by comparing all uphill walks on all landscapes with particular statistics. For power law walks, we noted that the variability across landscapes 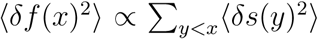 - suggesting that the correlation between individual steps is weak enough to not contribute (at least in a scaling sense) to the overall variability of evolutionary trajectories. However, for natural populations we are usually interested in the variability between two evolutionary trajectories starting from a single genotype on the *same* landscape. This is important to understand populations separated by spatial structure, the variability of evolutionary outcomes, and in experimental settings to understand the statistics of parallel evolutions from a single common ancestor.

The analysis of adaptive walks suggest that the conditioning on past evolution is important to understand this variability; therefore we should consider two adaptive walks starting from a genotype which is itself the result of evolution. More concretely, we consider the following scenario: a population evolves via an adaptive walk for some genetic distance *x_b_*. Then the population is separated into two non-interacting subpopulations, each of which explores a separate direction in the same landscape. We will refer to this trajectory as a *branching path*. The walk starts at a *root x* = 0, continues along a shared *trunk*, and then splits into two *branches* at the *branch point x_b_*, each branch continuing until the total number of steps in each reaches a total of *x*. We will be interested in tracking the fitness *f_i_*(*x*) relative to the root for the two paths *i* = 1, 2, and the difference Δ*f* (*x, x_b_*) *≡ f*_1_(*x*) *− f*_2_(*x*) between the fitness of the endpoints of the two paths which branched at *x_b_*. Figure 8 shows the trajectory of *f* for such a walk on a power-law correlated landscape.

**Figure 8:**
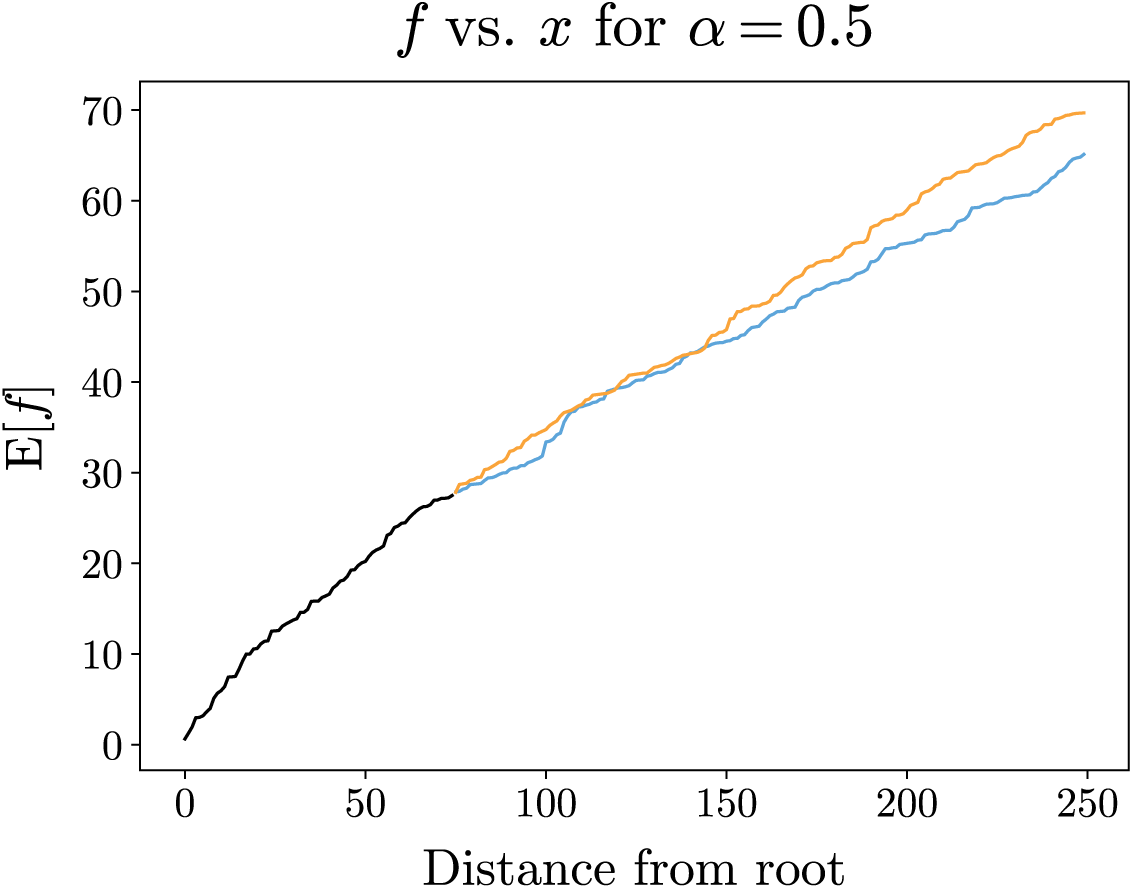
Typical example of cumulative fitness gain *f* for the two branches of a branched trajectory in a power-law correlated landscape. Paths branch at end of black curve (a distance of *x_b_* = 75 steps from root of walk). Correlations between branches quickly decay, but global structure of epistasis keeps relative variability of *f* small.

On power-law correlated landscapes, simulations show that long after the branching, the variance between the two branches on the same landscape behaves the same as the variance of a single path across different instantiations of the random landscape (trivially true for the independent fitness model). Specifically, Figure 9 shows the ratio of Var[Δ*f* (*x, x_b_*)] to twice the between-landscapes variance of the post-branch parts of the walk: 2Var[*f* (*x*) *− f* (*x_b_*)] (with the factor of two accounting for the variance of the difference between two — putatively — independent fitness increases.) As expected, Var[Δ*f* (*x, x_b_*)] is smaller than 2Var[*f* (*x*) *− f* (*x_b_*)] due to correlations induced by the shared trunk. However, the ratio approaches unity for *x ≫ x_b_*, suggesting that, indeed, variance between landscapes behaves the same as variance between branches in the same landscape at large genetic distances. We expect that the ratio approaches unity as a power of 1*/*(*x − x_b_*).

**Figure 9:**
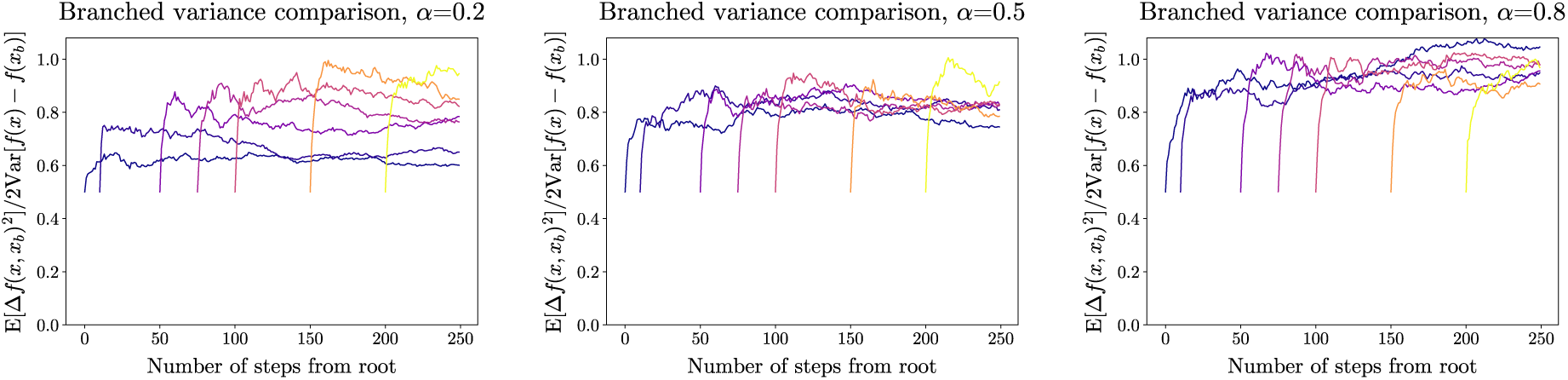
For branched random adaptive walks, ratio of the between-branch variance on a single landscape, to the single branch variance across landscapes: Var[Δ*f* (*x, x_b_*)]*/*2Var[*f* (*x*)*− f* (*x_b_*)]. The different colors correspond to different distances, *x_b_*, of the branch point from the starting point. Results for power-law correlated landscapes with several values of *α* are shown. The variance ratios saturate to unity, suggesting that variability between different paths on the same landscape behaves similarly to variability between paths on different landscapes with the same statistics.

The numerical results suggest that for longer evolutions, there is not much difference between taking multiple paths on the same fitness landscape, or taking paths on different fitness landscapes. For a branching path, the shared part provides some extra correlation, but this only persists for short distances from the branch. The long-term variability between branches only depends on the long-range statistics of the fitness landscape.

## 5 Greedy and abstemious walks

Thus far, we have considered walks that take uphill steps with probability independent of the fitness increment, *s*. However, in actual evolutionary processes the probability of fixation can depend on *s*. A natural question, then, is whether it pays to be more *greedy* (weighting towards larger *s*) or more *abstemious* (weighting towards smaller *s*) in the long run.

In general, whether being greedier increases or decreases the maximum fitness reached depends strongly on the statistics of the landscape and the resulting conditioning on past evolution. It has been previously found that in the independent fitnesses model, greedy walks do better than random (or abstemious) adaptive walks. For the *NK* model, the most effective walk type depends on the choice of parameters [51]; for small *ξ* (close to independent) or large *ξ* (close to additive), greedy walks are better, but for intermediate *ξ*, *reluctant* walks which take the smallest possible steps (the most abstemious dynamics) are better. Reluctant and greedy walks have also been studied in the context of the Sherrington-Kirkpatrick model in physics [54].

We will analyze a family of strategies with varying amounts of greediness on both exponentially correlated and power law correlated landscapes. We will also be interested in rare (but attainable) adaptive walks whose fitness trajectories are highly anomalous.

### 5.1 Adaptive walks with weighted choices of steps

With birth-death fluctuations in the strong-selection weak-mutation regime, the probability of fixation of a beneficial mutation is proportional to the fitness benefit, *s*. We only need a slight change in the analysis to understand the dynamics under such a weighting, or, indeed, weighting by any power *s^κ^* (although we know of no natural biological process that would correspond to negative *κ*.) Because the distribution of non-negative *s* is exponential in the limit *−µ ≫ σ*, for a large class of weighting functions we have:

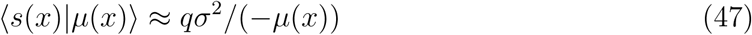

where the “greediness factor” *q* is the ratio of the average step taken to the average positive step available: it parameterizes the greediness of the walk. For weighting *s^κ^*, *q* = Γ(*κ* + 2). For unweighted steps *q* = 1; with fixation probability proportional to *s*, *q* = 2. Abstemious walks can be analyzed by taking non-negative *q <* 1. As we will see, the change in average step size will not change the scaling of *f* with respect to *x*, or the critical *µ* before getting stuck; however, it can change the maximal fitness reached, *f*_max_, by a multiplicative factor.

On exponentially correlated landscapes, as long as the evolutionary rule is local, the statistics of the current DFE only depends on the current fitness *F* (*x*). Therefore, the current *µ*(*x*) does not depend on the shape of the fitness trajectory; asymptotically in the limit of large *L*, the value of *F* (*x*) reached when the probability of an uphill step is *O*(*L^−^*^1^) is independent of *q*. The walk will proceed until *F* (*x*) is in a quantile *a*(*q*)*/L* of its cumulative distribution function. The *L*-independent coefficient *a*(*q*) will depend on the correlation length, *ξ*, of the landscape, and the results about the NK model from [51] suggest *a*(*q*) will be non-monotonic in *q*.

For power law correlated landscapes, *q* has a non-trivial effect which can be analyzed simply. The analysis in Section 3.3 relied on self-consistently solving Equations 29 and 36 given the response kernels. For weighted walks, we replace Equation 36 with Equation 47. A simple substitution shows that if *µ*(*x*) and *s*(*x*) satisfy the original equations, then 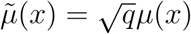 and 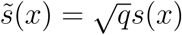 satisfy the weighted walk equations.

If we compute *f*_max_ as in Section 4.1 using *µ̄*(*x*) and *s̄*(*x*), we find that the walk reaches a local maximum *f*_max_(*q*) after *X* steps when

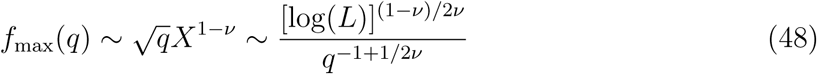

with *ν* = (1 *− α*)*/*2. Note that 1*/*2*ν* = 1*/*(1 *− α*) *>* 1 for all positive *α*, and *q >* 1 corresponds to greedy walks. We thus conclude that more abstemious walks go further than greedy walks. Being greedy is good in the short term but bad in the long term.

Though the individual fitness gains *s̄*(*x*) scale as 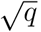 (larger for greedier walks), the total number of steps before getting stuck goes as *X ∼* [log(*L*)*/q*]^1^*^/^*^2^*^ν^* — larger for more abstemious walks, and enough to overcome their smaller step sizes. Concretely, going from randomly chosen mutations to fixation probability proportional to *s* decreases the maximum fitness reached by a factor of 2*^−α/^*^(1^*^−α^*^)^. The suppression is strongest for *α* near 1. (But note that for *α* = 1 this result does not apply as the maximum fitness is of order *L* independent of *q*: *q* only determines in which order the steps are likely to be taken).

Contrasting the results for power-law correlated and NK model landscapes, we see that whether being greedy or abstemious is better in the long run depends on the the statistics of the landscape and the past evolution. A surely interesting question is what happens for real fitness landscapes.

In larger populations, the fixation of mutations tends to be greedier than in the strong-selection, weak-mutation regime on which we have focused. Multiple mutations will arise in each generation and compete with each other: this yields fixation probabilities that grow much more rapidly than linearly in *s*. However large populations are polymorphic with the population spread out over many genomes — effectively taking many paths in parallel — and the behavior is much more complex; we discuss this briefly in Section 8.

### 5.2 Slow but steady adaptive walks

The above results suggest that to reach as high as possible fitness in an adaptive walk on a landscape with long-distance correlations, one should carefully — and artificially, although using only local information and memory — chose which uphill steps to take, being as abstemious as possible. Although unlikely relevant biologically, this is of potential interest for evolutionary algorithms, thus we summarize some results here, leaving the analysis to Appendix C.5, and raise some interesting questions.

The analysis carried out above is correct for any fixed *q* for large *L*. What happens if we choose much smaller *q*, e.g. *q ∼ L^−ζ^* for some positive *ζ <* 1? In particular, is it possible to take of order *L* steps before getting stuck? We show that, in power-law correlated landscapes, it is indeed possible. By considering only paths that do not change any site more than once, we find a lower-bound on the fitness reached:

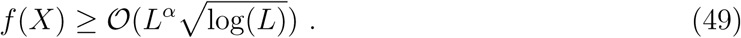

As explained in Section 2.3 the absolute maximum fitness on the hypercube is of order *L*^(1+^*^α^*^)^*^/^*^2^. For *α* = 1, the additive model, the above bound — up to the log factor which is unreliable in this case — is the same as this; but of course, any strategy works perfectly for additive models. In the opposite limit, with independent fitnesses, *α* = 0, the analysis breaks down but naive extrapolation gives 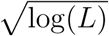 with no power of *L*, the same form as natural walks which have *q* = 2.

For power-law correlated landscapes how well can one do if even longer walks are allowed, so that many mutations are reversed with some — presumably many — reversing multiple times? This is a complex question which we will not endeavor to answer here. But the method of conditioning on the past should enable some exploration even in this regime. We leave as an intriguing open question whether, using only *local* information, the lower bound of the maximum reachable fitness in Equation 156 can be beaten.

Note that there are already subtleties in actually *finding* a path by choosing from the lowest *q* fraction of the available positive steps. This could be done by trying many steps and picking the lowest uphill one: in the regime where *−µ* is still of order *σ*, only roughly 2*/q* steps need to be tried. But once this is done, then in principle the future possibilities are conditioned on the *s*’s tested but not chosen. Due to this, the permutation symmetry in the yet-untaken mutational directions can no longer be used. We suspect that this would not actually affect the scaling of the fitness achievable, but that would require a much trickier estimate of the effects of the conditioning-by-testing. What we have shown, is the *existence* of an uphill path that can reach a fitness gain of order 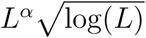, but it is not quite clear how to find it with only local information.

What if some global information is available? In the independent fitnesses model, we can come up with a good strategy given only knowledge of the statistics of *F*. If from each point the smallest available step is taken, then the walk can continue for of order *L* steps of size *∼* 1*/L* before *−µ*(*x*) becomes of order *σ* and the number of available steps starts being limited. Along such a walk, *L*^2^ neighbors will have been “seen”. Therefore a good strategy is to take the smallest step until a genotype with fitness of quantile *O*(*L^−^*^2^) is observed, take that step, and – most likely – get stuck there. With appropriate tuning of the decision point, with high probability such a fitness will be observed.

If information about the landscape itself is known, we can ask: how high a fitness increase will the best possible path reach from a random initial genome? In the independent case, a direct computation (Appendix C.2) suggests that this goes as *L*^1^*^/^*^4^. More generally, answering this question is likely to be very difficult analytically because of the strong correlations among different directions, and is numerically intractable without generating the whole landscape and exploring it — exponentially hard in *L*.

## 6 Beyond Gaussian landscapes

A key question about all our results is the generalizability of the quantitative and qualitative features to other models of epistasis. In particular, how sensitive are the adaptive walks to the assumptions of Gaussianity of the random landscape? Addressing this generally is beyond the scope of this paper, but we can begin to explore by considering *mixed landscapes* whose fitness function is a linear combination of an epistatic Gaussian function and a non-Gaussian additive part. More precisely, the fitness *F_tot_* is given by

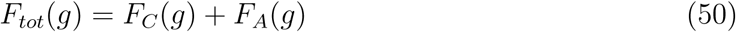

where *F_C_* is a distance-dependent correlated landscape, and *F_A_* is an additive landscape. Both parts of the landscape depend on the same parts of the genome, but they are statistically independent. Note that this is different than having a fraction of the sites on the genome not participating in the epistasis with mutations at those sites simply adding to the fitness: in that case adaptive walks can always continue until all the non-epistatic mutations are exhausted. In contrast, the models we consider have many local maxima and adaptive walks can not necessarily take *O*(*L*) steps. The RMF model is the simplest example, which assumes *F_C_*is the independent fitnesses model.

One way to understand models of this mixed-type is to think of the additive piece as being a modification of the dynamical rules. Recall the general transition rule for steps, to be taken with probability *P* (*g → g^t^*) = *ϕ*(*s_g→g′_*)*/Z*. With a mixed landscape, we can decompose the fitness benefit, *s*, of each potential mutation into an additive part, *s_A_*, and the part from the correlated landscape, which we will call *s_C_*. Then, the probability of taking a step becomes

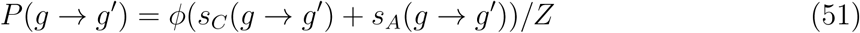

After convolving with the distribution of *s_A_*, we can consider this to be a modified transition rule for steps, *s_C_*, to be taken in the correlated part of the landscape. With *ϕ*(*s*) still a step function, the additive part of the landscape can compensate for the non-additive part; the path can take negative steps in the correlated landscape so long as a sufficiently positive step in the additive part is available to make the total fitness step positive.

The effects of the loosening of the requirements for steps in the correlated part of the landscape can keep the average available fitness gain, *µ_C_*, of the correlated part of the steps from getting too negative and allow for long evolutionary trajectories. In the limit of infinite *L*, *µ_C_*(*x*) will saturate for large *x* and from then on the statistics of the available steps will no longer change. In Appendix C.3, we use the modified transition rule together with the response kernels to directly study combinations of an additive model with a power-law-correlated model, first analyzing the saturation of *µ_C_*.

A key question is the behavior large finite *L*: when can the additive part of the fitness landscape stop walks getting stuck at local maxima? We first consider the simplest case: the RMF model (*α* = 0) [49]. In this case, it is simple to show that a Gaussian additive part with relative variance 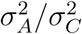 larger than *O*([log(*L*)]*^−^*^1^), will ensure that with high probability there are always uphill steps available. This result also holds for all power-law correlated landscapes (Appendix C.3). This quantifies the crossover from walks that get stuck after a power of log(*L*) steps, (Section 4.1) to walks of length *O*(*L*) (which in previous work were analyzed in the infinite *L* limit only [49, 56]). For any fixed non-zero 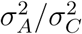, in the limit of large *L* adaptive walks can take *O*(*L*) steps. We note that the behavior depends on the tails of the independently random part of the RMF landscape; for example, with exponential tails there is a phase transition at finite 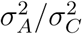 in the limit *L → ∞* [56].

It is instructive to compare how much of an additive part is needed to avoid random adaptive paths getting stuck, with the probability that there *exists* an uphill path of length *O*(*L*) (a “traversable” path [29]). An upper bound for the relative variance needed is only 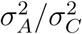 ∼*O*(*L*^−2^). Here, the strategy is to pick a step with positive *s_A_*and random *s_C_* with *|s_C_ | < s_A_*. We can do this with high probability so long as *σ_A_ ≫ L^−^*^1^. With this strategy, the induced *µ_C_* (*x*) is random and *O*(1), so the walk can continue until all positive additive steps are taken. Such a walk would only have fitness gain of *O*(1); it remains an open question if a more refined strategy can lead to large fitness gains.

The effects of additive parts of the fitness function are more dramatic if the additive parts of the step-sizes are broadly distributed, especially when there are rare mutations that have very large fitness effects. As an explicit example, we consider a distribution of *s_A_*with a power law tail so that the cumulative distribution function is

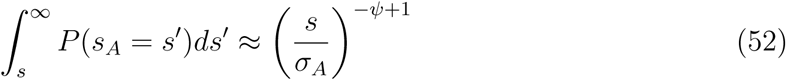

for large *s*. For *ψ >* 3, the scale *σ_A_* sets the standard deviation; for smaller *ψ*, the variance is infinite. When the *µ_C_* of the correlated landscape is large and negative, adaptive mutations must involve rare values in the tail of *s_A_*. If there is a large negative step *s_C_* in the correlated landscape, the probability that the mutation is net adaptive is given by

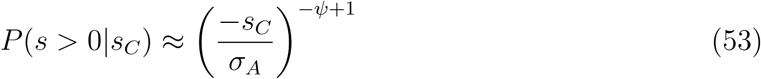

So long as this probability is greater than *O*(*L^−^*^1^), one of the *L* possible mutants, is likely to have sufficiently large *s_A_* and the rare, high effect mutation will prevent getting stuck at a local maximum. The correlations in the landscape reflected in the response kernel structure, implies that taking deleterious steps in the correlated part of the landscape makes the DFE average *µ* tend to increase becoming less negative. Therefore if the large additive steps are rare but not too rare, there will be a dynamical balance: evolution proceeds by usually taking small uphill steps in the correlated landscape, and occasionally uses the broadly distributed piece to keep going and “release the pressure” on the correlated landscape. This dynamics is studied in detail in Appendix C.4. We find that the effects of the additive piece are strong enough to allow this process to continue if

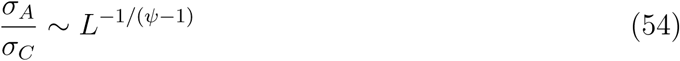

(with 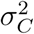 the variance of the correlated parts of the steps). Thus only a very small additive piece is needed to un-block long adaptive walks. (The limit *ψ → ∞* is like the Gaussian case which requires 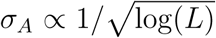. For *ψ =* 2, we only need an additive piece of O(L^-1^) for long walks — just enough to offset the smallest possible *s_C_* when *µ* = *O*(1).

Note that with a long-tailed distribution of available mutations, the behavior can depend strongly on the probability, *ϕ*(*s*), of a mutation fixing. If *ϕ*(*s*) *∝ s*, then for *ψ <* 2 the largest effect (“jackpot”) mutations would be chosen which modifies the nature of the walks and their dependence on *L*.

The analysis of the additive-plus-correlated landscapes shows explicitly how long-term evolutionary dynamics can depend sensitively on the tails of the distribution of fitness effects of mutations. Rare large effect mutations can drive evolution in parts of the landscape where the “typical” modest effect mutations — the bulk of the DFE — are deleterious. And such rare large mutations can unlock new sections of the fitness landscape and enable the typical DFE to again drive the evolution. As we will discuss below, these phenomena do not require an additive part of the landscape: they can occur whenever the distribution of effects of available mutations around some genomes has a long tail.

## 7 Time-dependence of walks and landscapes

### 7.1 Adaptive walks in time

The properties discussed thus far all depended only on the *geometry* of the random adaptive walks. The pseudo-“dynamics” was in terms of number of mutations. However we are also interested in how the evolution progresses in *time*. The simplest way to convert the mutation steps to actual dynamics, which is essentially correct for modest size populations, is to take the time between mutations to be exponentially distributed with characteristic rate 1*/τ_M_* proportional to the total population mutation rate (the product of the mutation rate per site, *L*, and the population size). If a mutation is deleterious, it is purged with probability one in a time much shorter than *τ_M_*. If the mutation is adaptive, it fixes in time much shorter than *τ_M_*. We will henceforth set *τ_M_* = 1.

With this dynamics, evolution on any Gaussian landscape slows down dramatically with time. Figure 10 shows an example of the fitness gain as a function of time, plotted together with the DFE average *µ*(*x*(*t*)). When *µ* is a large and negative (which it will systematically go to as the number of mutations increases), the waiting time for the next mutation is very long. More precisely, we let *τ_U_* (*x*) be the time taken for the *x*th uphill step. The average time 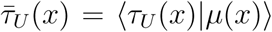 goes as 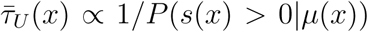, which with our Gaussian fitness DFEs gives

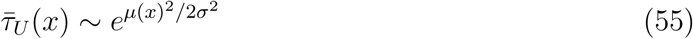

when *−µ ≫ σ* (Appendix D.1). There is a super-exponential slowdown in the beneficial mutation rate due to the rarity of uphill directions; 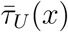 will reach *O*(*L*) as the number of beneficial mutations becomes *O*(1) and at that point the walk is likely to stop at a local fitness maximum.

**Figure 10:**
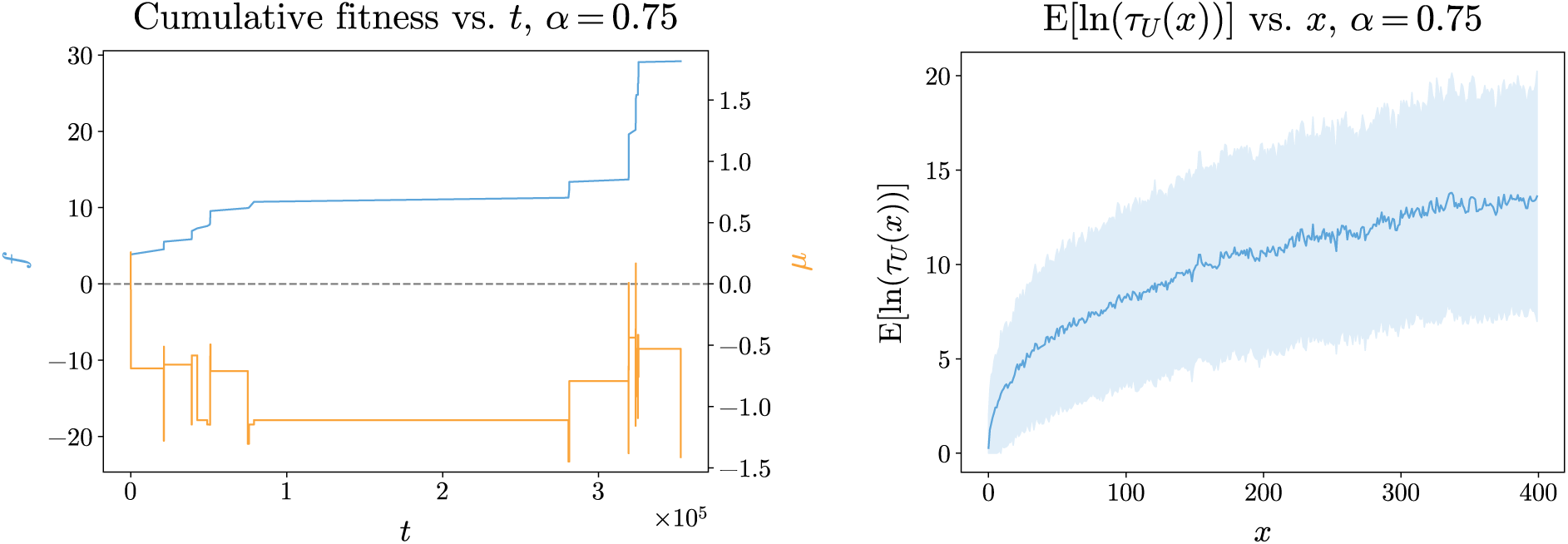
Time-dependence of adaptive walks. Left panel shows an example of the fitness gain versus time, *F* (*x*(*t*)), of a typical adaptive walk (blue) on a power-law correlated landscape with *α* = 0.75, along with the dynamics of the DFE average, *µ*(*x*(*t*)) (orange). The typical time between steps, *τ_U_* (*x*), goes as exp[*µ*[*x*(*t*)]^2^*/*2*σ*^2^], so that the logarithm of the waiting time for a beneficial mutation to arise and fix has mean and variance that both increase rapidly — although not monotonically — as the evolution progresses (right panel). Time is in units of total population mutation rate.

While there are still many available beneficial mutations, the logarithm log(*τ_U_* (*x*)) is roughly normal, with mean and variance that increase as *x*^1^*^−α^* (Figure 10, right panel). The broad distribution of the waiting time between mutations means that there is not a steady slowing down: initially, there will be a small number of quick, large effect mutations, but after a while there will be a mixture of long and short intervals as seen in Figure 10, before the evolution stops. Of course, if fluctuation effects are included, deleterious mutations can fix, or, in larger populations, “tunneling” through deleterious intermediaries can occur: the dynamics then becomes more complicated.

### 7.2 Time-dependent landscapes

In nature, and even in the lab, environments are never completely static and whether or not mutations are beneficial can be very sensitive to even small changes in the environment [40]. As this can result from changing the subtle balance between the deleterious and beneficial consequences of a mutation, random fitness changes are a natural caricature of these effects. More generally, we would like to understand the effects of conditioning on evolutionary history in environments that have been somewhat different than the current one. Thus we are led to consider evolution in slowly changing fitness landscapes — sometimes dubbed “seascapes”.

To make progress, we again focus on Gaussian correlated fitness functions, but now with temporal correlations as well. The response kernel framework can be used to analyze fitness landscapes that change simply in time — we comment later on more complex time-dependence.

Given a time-dependent fitness function *F* (*g, t*), we can define a *time-dependent divergence function D*(𝓁, *t, t′*) as

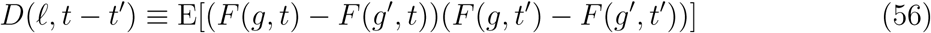

where *|g – g′|* = 𝓁. We have restricted ourselves to the case where *D* is time-translation invariant. The time-dependent divergence therefore is the correlation of the fitness difference between genomes a distance 𝓁 apart, with the fitness difference between the same genomes a time *t* in the future (or past).

Consider the case of exponential decay of time correlations, given by

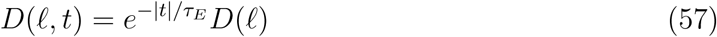

At any given instant in time the statistics are the same as in the time-independent case; however, fitnesses at different times become decorrelated on a timescale of *τ_E_*. The exponential time decay corresponds to a memoryless process, which simplifies both analytical and computational understanding (see Appendix D.2 for details).

Time-dependence qualitatively changes the nature of random adaptive walks — especially their dynamics. The system approaches a statistical steady state for which the distribution of the time to take an uphill step, *τ_U_* (*x*), loses explicit dependence on the mutational distance from where it started. The average time between steps in the statistical steady state, 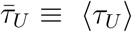 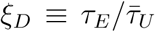 plays an important role in the overall dynamics — but there are a whole spectrum of time scales. The behavior in the random seascape is controlled by the balance between finding uphill steps, in *τ_U_* (*x*), and significant decorrelation of the landscape tending to decrease *−µ*(*x*) back towards zero on time scale *τ_E_*. Calculations in Appendix D.3 show that for *τ_E_* large, our focus, the typical value of the DFE mean in steady state, which we denote as *µ_c_*, scales as

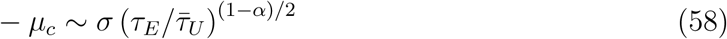

This is because 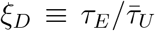 serves as an effective length scale: it is the number of steps taken in a time *τ_E_*, and gives the number of mutations in the past that still have significant effect on the current DFE, via contribution to the current *µ*(*x*).

From Equation 55, we know that *µ_c_* only depends logarithmically on *τ_U_*. Equation 58 then implies that in the dynamical steady state, to logarithmic accuracy, 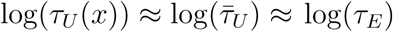 so that, for self-consistency we must have

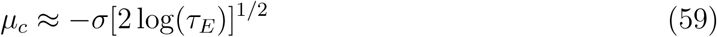

independent of *α* due to the sharp Gaussian tails of the distributions.

Numerical simulations confirm this predicted scaling: Figure 11 shows *µ_c_* versus log(*τ_E_*)^1^*^/^*^2^. Each simulation was run for a fixed number of uphill steps *X* = 200. For low *α*, the relationship is linear over a large range. For larger *α*, the behavior is linear for modestly large *τ_E_*, but for longer *τ_E_*, namely those where *|µ_c_| ≪ X^−^*^(1^*^−α^*^)^*^/^*^2^, *µ* did not saturate in the number of steps allotted for the simulations. The data in Figure 11 is thus plotted only for *τ_E_* less than this characteristic crossover value.

**Figure 11:**
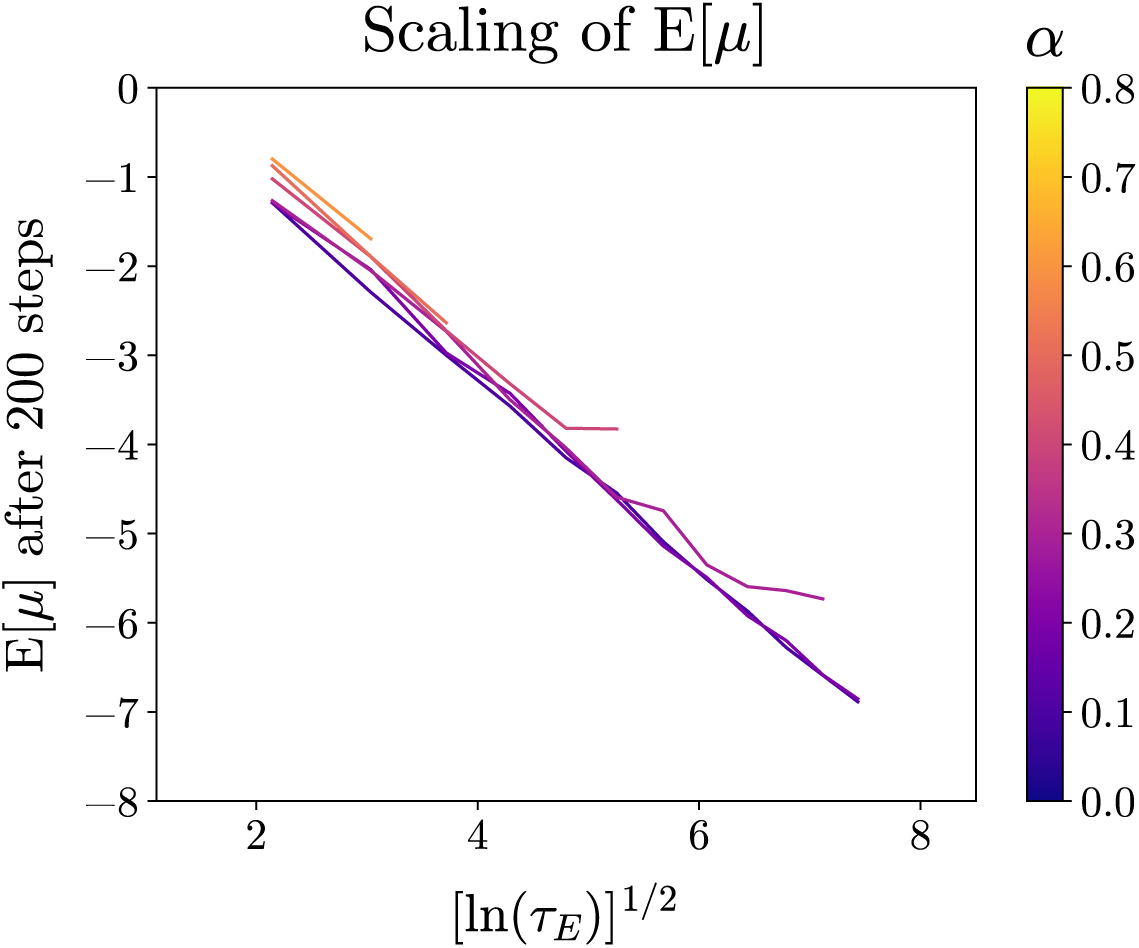
Scaling of the steady state average available fitness step, *µ_c_*, with log(*τ_E_*)^1^*^/^*^2^ for walk of length *X* = 200. Relationship is linear until ln(*τ_E_*) *∼ X*^(1^*^−α^*^)^*^/^*^2^, where simulations have not saturated (not shown). For large *α* and *τ_E_*, *µ* did not saturate for any simulations with this *X* due to slower growth of *−µ*(*x*).

The typical fitness gain of the *x*th mutation taken at the time *t* where it fixed, *s*(*x, t*), is of order *σ*^2^*/*(*−µ_c_*). This, times the average rate of fixing of beneficial mutations, 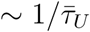, is the “fitness flux” — the apparent rate of increase of fitness [45]. If *−µ_c_* is small — when the landscape changes quickly relative to mutations — then the fitness flux is high. Conversely, the fitness gain in any one environment — that is, the maximal fitness of any genotype reached during evolution as measured in the fitness landscape at a fixed time — scales, when the changes are slow, as *ξ_D_σ*^2^*/*(*−µ_c_*). The faster the environment changes, the less adaptation occurs measured in any fixed environment.

The details of the dynamical behavior are much more interesting and subtle. Due to the broad distribution of *τ_U_*(*x*), to understand the qualitative behavior of the dynamics we must analyze the distribution of mutation times and not just the average 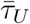. This can be done asymptotically for large *τ_E_*, as carried out in Appendix D.4. A key quantity is *µ*(*x*), here the average-available fitness step *just after the previous step*. For very slowly changing landscapes, 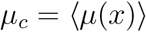, is much larger than *σ*, and their ratio, 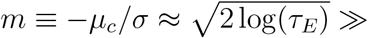 1 is the basic large parameter that enables the analysis.

There are two classes of mutations: those that occur in time much smaller than *τ_E_*, so that the landscape does not change significantly between mutations, and those with long enough waiting times that some significant changes in the landscape have occurred. This can be seen by analyzing *r*(*t*), the probability per-time of a beneficial mutation occurring a time *t* after the previous mutation. This rate is increased by the exponential decay of the average available fitness step; since 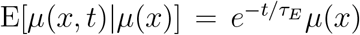, for *t ≪ τ_E_* we have 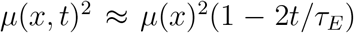 with the stochastic part of the change (of order 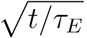) only giving negligible corrections (Appendix D.4). Thus *r*(*t*) increases roughly exponentially with time after the previous step.

The first beneficial mutation will typically occur at a time *τ_U_* at which 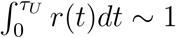 For typical *−µ*(*x*), the time-dependence of the landscape turns out not to matter: the next mutation is very likely to occur before the environment changes by enough to make a difference (i.e., *r*(*τ_U_*) *≈ r*(0)). The *typical τ_U_*(*x*) are broadly distributed but they are much smaller than *τ_E_*by a factor exponentially small in the large parameter *m* (Equation 176).

In contrast to the typical relatively fast mutations, there are rare situations in which *−µ*(*x*) is anomalously large so that a beneficial mutation is unlikely to occur until after the environment has started to change substantially. The waiting time for these is proportional to how much *−µ*(*x, t*) has to decrease for the mutation to have a high chance of occurring. Although such waiting only occurs for a very small fraction of cases, the average waiting time 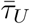 is dominated by these rare situations, which have waiting times of order *τ_E_/m*. As this waiting time is still much smaller than the landscape-decorrelation time, *τ_E_*, these can be thought of as situations that are only “slightly-stuck”. Note that although mutations in such situations are very unlikely to occur before the landscape has changed, there will typically be some beneficial mutations that *could* have occurred without waiting, but there are few enough of these that this is unlikely to occur.

For power-law correlated landscapes, the effects of past history from Equation 58 gives 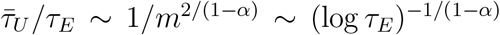. As many slightly-stuck situations will occur in a decorrelation time, the total number of steps per *τ_E_* is well approximated by 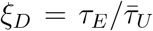 with smaller variations around this. The dynamics is thus quite smooth on time scales of *τ_E_* and length scales of *ξ_D_*.

Our analysis implies very heterogeneous dynamics. For all but a small fraction, (*∼* 1*/m*^(1+^*^α^*^)^*^/^*^(1^*^−α^*^)^) of the steps, the effects of the time-dependence of the landscape are negligible. Many relatively rapid steps are taken as if in a static landscape, until a rare slightly-stuck genome is reached: the population then “waits” for small changes in the environment — even though for large *L* it might still have had many potential beneficial mutations available before the landscape changed. Of course, as for most quantities in these Gaussian seascapes, the parameter that controls the intermittent dynamics is only logarithmically large — in this case, 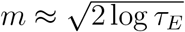.

The numerics confirm this picture. Figure 12 shows a typical example of the dynamics of fitness gain with exponential temporal correlations. There is a period of rapid adaptation before the time-dependence of the environment comes into play: the fitness flux is initially large and *µ* decreasing (left panel). As the transient dynamics slows down the time-dependence of the fitness seascape matters; *µ* and the fitness flux (whose integral is plotted in the figure) saturate and then fluctuate stochastically with steady-state distributions (middle panel). The middle panel shows the heterogeneity in the fitness flux; the fitness looks as if it is taking very discrete steps. Looking more closely (right panel), one can see that there are periods during which mutations occur rapidly, interspersed with occasional anomalously long gaps between successive mutations, as predicted.

**Figure 12:**
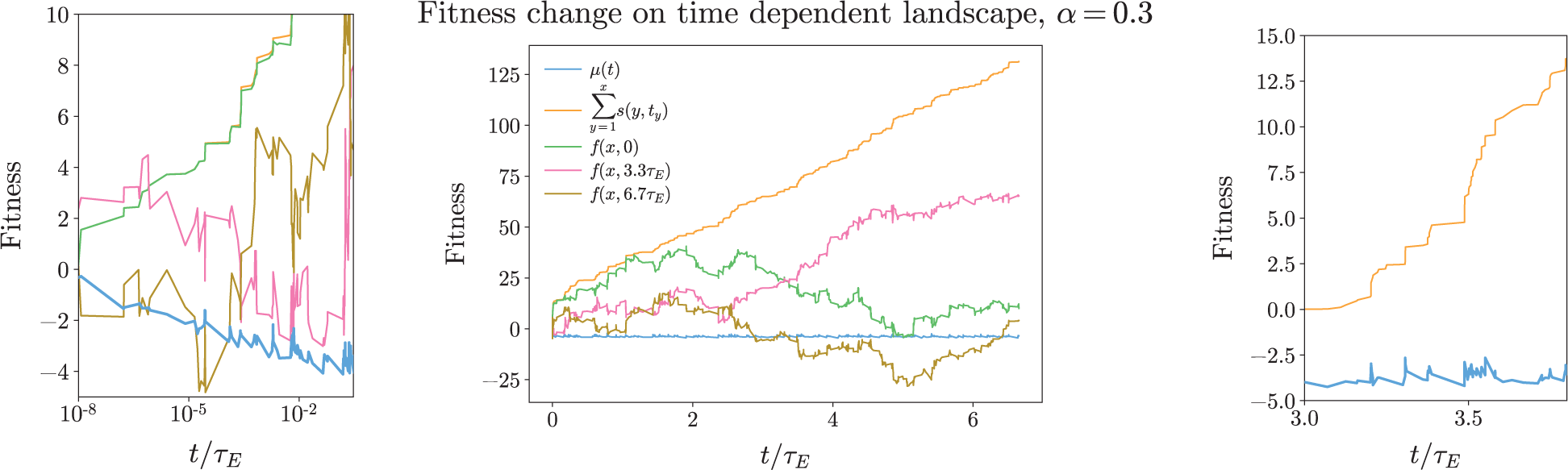
Dynamics of *µ* and fitness gains, *f*, for uphill random walks in exponentially time-correlated power-law random landscapes, with time in units of the landscape correlation time *τ_E_* = 10^8^. Left panel: rapid adaptation occurs at start of evolution as *µ* (blue) decreases. Middle panel: cumulative fitness flux 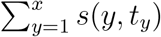 (sum of fitness gains evaluated at times of mutations, orange) increases linearly in time. For any fixed time, *t^∗^*, fitness gains in the landscape at *t^∗^*, *f* (*x*(*t*)*, t^∗^*), are guaranteed to be positive only for times near *t^∗^*. Gain in initial landscape, *t^∗^* = 0, (green) is correlated with fitness flux at start but then becomes a random walk with diffusion coefficient 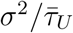. Fitness gains in later landscapes, at *t^∗^* = 3.3*τ_E_* (red) and *t^∗^* = 6.7*τ_E_* (brown) are random, then correlated with fitness flux, then random again. Right panel: In steady state, on a finer temporal scale, *µ* is seen to fluctuate (blue) leading to heterogeneous fitness flux. Cumulative fitness flux (orange) plotted from 0, for convenience.

Although the integral of the fitness flux is well defined, the increase in fitness, *f* (*x*), no longer is. However, the fitness gain measured in the environment at a chosen time *t^∗^*, *f* (*x*(*t*)*, t^∗^*), is well defined (Figure 12). With respect to the initial landscape (*t^∗^* = 0), the fitness first increases and then does a random walk with diffusion constant 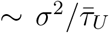 as the dynamics become uncorrelated from the initial landscape. The transition from uphill behavior in the initial landscape to the random walk behavior depends on *D*(𝓁). If anticorrelations with previously observed genotypes are strong, the dynamics drives the fitness in the original-landscape fitness back to zero before commencing the random walk. If correlations with other genotypes are weak, then the random walk commences almost immediately and is centered around the highest fitness value reached in the initial landscape during the early part of the trajectory. This can be seen most easily by examining the independent and the additive models, respectively. In the independent model, once the steps are no longer biased towards being uphill, any single step will tend to take the fitness to a random genotype (with fitness distribution centered around zero). Conversely, for the additive model, future steps are independent of the previous fitness gains; therefore, the average value of the fitness in the present landscape at future times will be the average value of the current fitness — the evolution “remembers” that it gained fitness when the reference landscape at *t^∗^* was the one driving the dynamics. Models that lie between the two extremes — such as the exponentially and power-law correlated landscapes — will either have smaller memory of the fitness gain, or memory that decays more slowly with time than for the independent fitnesses model.

The above analyses of the intermittent dynamics hold when there are significant anticorrelations in the landscape, more quantitatively, when the anticorrelations extend over the dynamically induced length scale *ξ_D_*. For landscapes that are additive up to a length-scale *ξ* (like the exponentially-correlated landscapes), the behavior of the dynamics is qualitatively different for *ξ_D_ < ξ* than for *ξ_D_ > ξ*. If *ξ_D_ > ξ* then the previous analyses hold, and there is intermittent dynamics where the population occasionally waits a long time for the landscape to change before taking a step. However, if *ξ_D_ < ξ*, then the landscape is effectively additive and *µ_c_* is always small (zero for the perfectly additive model). In this case evolution always proceeds steadily as beneficial mutations are always available. Such seascapes forget past fitnesses quickly enough that their correlations remain small and do not drive *µ*(*x*) to large negative values.

An important question is: In what sense are the fitness landscape neighborhoods in steady-state atypical? They do not look like neighborhoods of “typical” points with the same DFE mean *µ*(*x*) *≈ µ_c_*. To see this, consider what happens when the dynamics starts at some random point on the fitness landscape with *µ*(0) *< µ_c_*. Figure 13 shows that the dynamics very quickly “resets” to what it would be if originally had *µ*(0) = 0. The initially large negative *µ* decays away due to a combination of the exponential time decorrelation at rate *τ_E_*, but also due to the history of the trajectory. As at early times, there have not been a large number of genotypes visited in the near past that had large negative *µ*, thus the response kernel does not have enough “memory” to drive down future *µ*. (This is analogous to the future evolution of a “typical” random trajectory conditioned only on the end point fitness, discussed in Section 4.2.) This lack of a past history causes the dynamics to “forget” its initialization.

**Figure 13:**
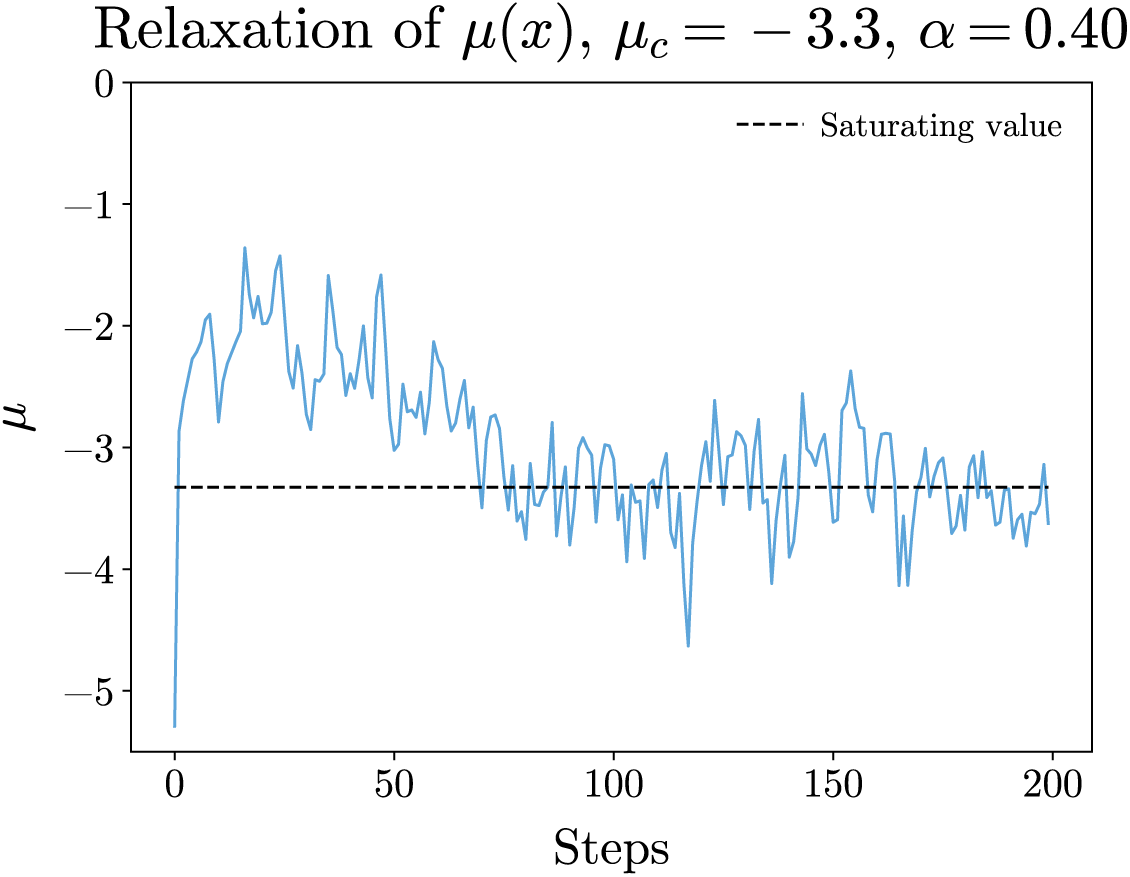
Dynamics of walk started from a randomly chosen point with anomalously negative *µ*(0) *< µ_c_* in exponentially time-correlated power-law landscape. Large negative value of the mean-available fitness step, *µ*, is “forgotten” as it quickly becomes greater than the steady state average value, *µ_c_*, then approaches this from above (as occurs with random initialization, *µ*(0) = 0, not shown), and then fluctuates around *µ_c_*. This illustrates that history of a trajectory, rather than just the local neighborhood parameterized by *µ*, matters, and in steady state, the genomes are not at all like typical points with *µ ≈ µ_c_*.

Further evolution then drives *µ* down towards *µ_c_* slowly; stabilization around *µ_c_* requires the development of a history of large negative *µ* in the near past. After some time of order *τ_E_*, *µ* will reach, and then fluctuate around, *µ_c_*. At this point, the last few steps (more precisely, the last *ξ_D_* steps) will all have had *µ ≈ µ_c_*. The population, now in dynamical steady state, has arrived at a location on the fitness landscape on a “ridge”. In the neighborhood of the current genome, most directions tend to decrease the fitness *F* sharply while relaxing *µ* to less negative values. But there are some special directions — back along the evolutionary trajectory — in which both *F* and *µ* do not change very much. The history of evolution has led the population to a region that has different geometry than a typical region with the same DFE average *µ*.

Though many of our results depend on the Gaussian nature of the landscape, the nature of local landscapes in dynamical steady state is less sensitive to it — and therefore may be a general phenomenon. The “ridge-like” character only requires that the current average size of potential steps is correlated with the past evolutionary trajectory as a whole. This type of dependence on the past means that to reach any sort of dynamical steady state in a stochastically varying high-dimensional seascape, the population must be in a region of the current landscape for which its whole past trajectory is consistent with the steady state, so that the memory of past history affects it in a steady-state manner. This criterion is very different than just matching the statistics of the single-mutation DFE.

The analysis of simple exponentially-decaying temporal correlations of the landscape suggests that even very slow time-dependence qualitatively changes the nature of evolutionary dynamics and the character of local landscapes conditioned on the past evolutionary history. Exponential time correlations “blur out” the effects of the static-landscape structure. We have seen that at long timescales in randomly changing environments, evolution reaches a dynamical steady state where it does not improve too much in any one landscape; however, the population is always improving in the current environment. This means that the rate of fixation of new mutations, the fitness flux, does not decrease, on average. But in direct head to head competition with an ancestor in the original environment, the fitness difference of the evolving population would increase only slowly and then vary randomly. In spite of this seemingly simple behavior, the complex dependence on the past remains, as evidenced by the fact that neighborhoods at steady state have a special “flat” direction in contrast to random points in the landscape with the same DFE (i.e. *µ ≈ µ_c_*); the latter tend to have uniformly large negative curvature.

### 7.3 General time-dependent Gaussian landscapes

Many of the concrete features discussed above depend on the simple exponential time-correlations of the landscapes. But within the Gaussian framework, one can consider more general time correlation, parameterized by a *time-dependent amplitude spectrum*. In short, each Fourier mode — roughly corresponding to each length-scale — has its own time-dependence. The most general form of the divergence function consistent with the distance-dependent statistical structure is given by

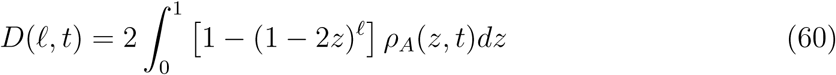

with *ρ_A_*(*z, t*) any function of time with non-negative temporal Fourier coefficients, 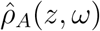 which could be used to parameterize it.

Though we can formally write down the general time correlation, our current methods are not sufficient to analyze cases beyond simple exponential time correlations — each *ρ_A_*(*z, t*) having the same pure exponential time-dependence — as in the previous section. Because, in general, every order of epistasis has a different time-dependence, generating the landscape on the fly becomes problematic since the conditional probabilities of the *µ* no longer have simple response kernel forms; they depend on the entire history of the known landscape. In principle, one could compute “space-time” response kernels which were a function of distance as well as time. However this would quickly become problematic numerically as in steady state the landscape could need to change significantly between each mutational step so that the number of variables would grow at least as *x*^2^ for a walk of length *x*, leading to matrix inversion on the order of *x*^6^ per step for a total complexity for a walk of length *X* of *X*^8^ which, while still polynomial, rapidly becomes intractable. This is likely still true if the space-time kernels were computed iteratively, although the complexity might then be held at *X*^6^.

Further development of, heuristic, asymptotic, and approximate numerical methods are needed to study general-time-dependent Gaussian random landscapes. Because of the complex environmental and evolutionary histories, and strong effects of conditioning on the history, the behavior is potentially especially interesting. For example, relatively rapid changes of the small-scale properties of the landscapes could open up possibilities to continue evolving in the much more slowly changing large-scale structure of the landscapes.

## 8 Discussion

### 8.1 Lessons from the theory

Our study of evolution on random landscapes with distance-dependent correlations has yielded some key insights into the interplay between evolution and epistasis. The distance-dependent divergence function enables model definition based on features about which one might have intuition, such as the relationship between genetic and phenotypic relatedness, with more flexibility and more explicitly than previous models (like the *NK* or RMF models). The simple relationship between the locally-defined (distance-dependent divergences) and globally-defined (rescaled amplitude spectrum) descriptions of epistasis facilitate classification and understanding of models; for example, we recapitulated the finding that *NK* models, in the appropriate scaling limit, correspond to correlations that decay exponentially in genetic distance (as in the mean field *NK* model [29]).

Our analysis of random adaptive walks on high-dimensional landscapes gives direct insight into how epistasis can shape evolution and *vice versa*. One key point is that the evolutionary dynamics depends more on *long-range* properties than *short-range*, local, properties. Intuition that the “ruggedness” of landscapes — in particular distributions of local maxima — determine how epistasis affects evolution is, in high dimensions, misleading at best. The high dimensionality of the landscape, and consequent large number of possible mutations, means that rare beneficial mutations can occur even when most mutations are deleterious. Subsequently such mutations can drive large gains in fitness. This is especially true when the distribution of fitness of potential mutations (DFE) is broadly distributed, as we showed in the analysis of mixed models.

A primary result of our analysis is that — and how — *the evolutionary history crucially matters*. The current distribution of available fitness effects of mutations (DFE) is determined by the past evolutionary trajectory as a whole, not just the recent past. Except in special cases — in particular with exponentially decaying correlations — it is not enough to condition over summary statistics like the current fitness, or the fitness gain over the last few mutational steps. For landscapes in which the fitness differences between genetically distant genomes can be large — in particular the illustrative class we have studied for which fitness differences typically grow as a power of genetic distance — mutations that occurred far in the past collectively contribute in important ways to the present DFE. This reflects the general conclusion that evolution is far more sensitive to large-scale properties of fitness landscapes than to local structure.

The dependence on the past leads to highly *non-generic* points on the landscape near which the statistics of interactions between mutations in the neighborhood of the current genome are very different than the neighborhoods of more generic, but superficially similar, genomes, such as those with similar DFEs. For example, at a random point, the double-mutant DFE (distribution of fitness difference to all genotypes distance 2 away) is determined solely by the current (single mutant) DFE, but when past evolution has occurred, the double-mutant DFE depends on an integral over the past evolution (a generalization of the dependence of *µ*(*x*) on the past that we have analyzed in detail). The non-generic nature of genomes conditioned on evolution is supported by other studies; in particular, on a variety of theoretical landscapes, epistasis is found to be stronger further along evolutionary trajectories [16, 24].

Differences in the longer-range statistics determine how the future evolution is conditioned on past evolution. We have shown that adaptive walks tend to take populations to places on the fitness landscape where they can continue uphill. Adaptive walks, while taking individual steps that are not special — other than being uphill — are overall rare and special paths through the landscape. In particular we have shown that they are very different than typical uphill paths that take the same number of steps and have the same total fitness increase.

Uphill evolution can continue as long as there are some beneficial mutations available. The depletion of beneficial mutations depends on the entire past trajectory. Our results suggest that long-range correlation leads to long-term benefits to “abstemious” walks where smaller fitness gains are accrued at each step; if the current DFE depends on the shape of the fitness trajectory, and not just the current fitness, “greedy” walks tend to drive down the average available fitness *µ*(*x*) and lead populations to dead ends much faster. An interesting open question is whether or not this is the case in real evolutionary contexts.

In some families of landscapes, in particular with an additive part of the fitness function, an adaptive walk can “unstick” itself from genomes near which there are very few beneficial mutations and move to where there are more. This is easier if the additive parts of the step distribution are long-tailed: evolution can proceed in a mixed additive-plus-correlated landscape by mostly steps that are uphill in the correlated landscape, only occasionally using a rare, large additive mutation to get unstuck.

Similar behavior occurs when the landscape changes slowly in time. In the statistical steady state that is reached after a long time, a large fraction of mutational steps — almost all in the limit of very slow rate of environmental change — are uphill in the current landscape and occur fast enough that the landscape is effectively static. Only a small fraction of the steps “wait” for more beneficial mutations to become available because of the slightly changed environment — but these few mutations occur slowly enough that they dominate the average time between mutations fixing and thus the average fitness flux.

Collectively, the various features of evolution on epistatic landscapes show that the statistical properties of fitness landscapes (and more generally seascapes) are insufficient for understanding evolutionary dynamics. What really matters is the statistical properties of the landscape (seascape) conditioned on past evolution — in general very different than unconditioned neighborhoods of the landscape. This conclusion is much broader than the particular class of landscapes we have analyzed.

### 8.2 Possible experimental connections

The genomic fitness landscapes we have analyzed are, of course, far from those that occur in nature. However some insights can be gleaned for experimental microbial evolution and some other real systems.

Richard Lenski has carried out *long-term evolution experiments* in *E. Coli* in a nominally simple environment with selective pressure designed to be due solely to glucose limitation [37]. This experiment has run continuously for over 65,000 generations, with 12 replicate lines. Quantitative analysis of the populations has shown that the fitness relative to the ancestor has been steadily increasing throughout the evolution, but the rate of fitness increase has slowed down, initially strongly and then more gradually [73]. The fitness trajectories have been fit by a rough model of diminishing returns epistasis that predicts logarithmic time-dependence of the fitness trajectories [21, 73]. The shapes of fitness trajectories will certainly depend on properties of the landscape, but our conclusion that they will depend on large genetic-distance scale properties, rather than just simple features such as overall diminishing-returns epistasis, suggests that better models are needed.

The time-dependence of the fitness, and the similarity among Lenski’s 12 separate lines, can not be compared to our analysis, even roughly. The experimental populations are large enough that many mutations arise, interfere, and add before any can fix. This tends to make the evolution of large populations much steadier than the highly sporadic nature of the evolution in small populations for which, as we have seen, the epistatic interactions can make the number of available beneficial mutations change stochastically as the evolution proceeds. Large populations are likely to average over some of these variations because of the multiple directions that they explore in parallel. In addition, the likely non-Gaussian nature of the evolution-conditioned DFEs likely makes the dynamics in both time and number of mutations quantitatively different.

A surprising feature of Lenski’s data, revealed by following trajectories of mutations by deep sequencing of populations every 500 generations, suggests that the rate of fixation of new mutations does not change much after the initial decrease (with the exception of the emergence of hypermutator strains) [23]. This could occur if the environment were gradually changing so that the increases in fitness relative to the ancestor were not indicative of changes relative to the current population — i.e. the fitness flux could decrease much more slowly than the rate of increase of fitness in the original environment. Evidence from the mutation frequency trajectories for the development of ecological interactions (including coexistence of evolved strains for much longer times than could be attributed to them by chance) [23], support this candidate explanation. This would mean that the systems are evolving in a landscape that is more like an externally time-dependent “seascape” than a static landscape. It is worth noting that even if the time-dependence of the environment had been steady and systematic instead of determined by the evolution itself, one would nevertheless have expected a period of rapid adaptation in an effectively static landscape followed by reduced adaptation in any one environment but nevertheless a relatively steady accumulation of further mutations. Lenski’s experiments do not seem to have reached a state in which the fitness increases in the original environment — measured by competing with labeled ancestors that dominate the population [73] — have become random. But that is not what would be expected if, in addition to mutations with epistatic interactions that depend on the environment, there were also a small but still available supply of unconditionally beneficial mutations (most simply with additive effects) that have not yet been depleted: these would cause steady increase in fitness even measured in the original environment.

An opposite extreme of genome-wide adaptive evolution is to artificially construct a large number of closely related genomes, and directly measure *empirical fitness landscapes*. Great advances in genomics have made it possible to map out portions of a fitness landscape by creating all possible variants at a small number of sites in the genome of an organism, and measure the relative fitness of each variant (see the recent review article [10]). Examples include a set of mutations in a single protein, *β*-lactamase, which increase antibiotic resistance [71], biosynthetic loci in yeast [27], and 24 loci on an RNA whose “fitness” is defined by its binding affinity to a GTP agarose resin [33]. Researchers can fit the landscapes to models or, more generally, decompose them, via their Fourier spectrum, as sums of all possible orders of epistatic interactions. A recent meta-analysis showed that the Fourier amplitude spectra of some low-dimensional (*L* = 4 *−* 7) complete landscapes range from nearly additive to highly epistatic (70% of variance from non-linear terms) [67]. Previous analyses had mainly focused on fitting Rough Mount Fuji or *NK* models to the empirical landscapes [48, 67], with limited success. More recently, correlation based measures have been used to characterize the statistics of empirical landscapes [1].

In addition to the question of what more general can be learned from empirical landscapes in particular contexts, there is the much broader question of whether evolution on high dimensional landscapes is fundamentally different than on low-dimensional ones. Most empirical landscapes are very low dimensional with the number of sites varied usually ranging from 10-20. Larger studies have been carried out — see [39] for a recent example where variants at 69 sites of an tRNA gene were considered – but they are far less comprehensive, only probing modest pairwise-distances — in the tRNA gene, only 9 or so. With a small number of sites, 𝓁, changed, high order epistatic effects — in the more general representation, larger larger length-scale components — are not probed. The combinatorial explosion of genomes and hence interactions with 𝓁 means that even rare, strong, interactions can contribute to the overall landscape for even modest 𝓁. The higher order interactions can dominate the long-distance statistics of fitness landscapes, which our modeling and analysis suggests can be a crucial driver of evolution via multiple successive mutations. Recent computational work on fitness landscapes from data suggests that epistatic interactions determine *available* evolutionary paths even in low-dimensional landscapes [63].

There are several problems with drawing lessons from empirical landscapes involving only very small parts of the genome. Such restricted studies leave out the potential for rare mutations on other parts of the genome that might have weak epistatic interactions with the sub-system studied, but sufficient to “unblock” the sub-system’s fitness landscape and enable evolution to avoid getting stuck at local maxima. Further issues are that the studies are conditioned on both past evolution, and the sub-space studied often has special properties. For example, for the *β*-lactamase protein, the collection of mutations analyzed were known to be jointly beneficial, while for the yeast study analyzed in [67], mutations were individually deleterious. We have shown that the properties of the fitness landscape in the neighborhood of a particular genome, depend heavily on conditioning: this includes on whatever is already known about relationships among mutations being considered.

Studying restricted regions of the genome involved in important processes is valuable for evolution of specific traits. Additionally, the interplay between evolution and epistasis in individual (and pairs of interacting) proteins has been extensively used to predict aspects of protein structure using evolutionary co-variation to infer functional relationships between residues [42, 43, 52]. But we feel that focusing on evolution of small subsets of genomes gives only limited insights about how even intermediate-term evolution is affected by, and determines, epistasis among the great number of other possible beneficial genetic changes.

To glean information about epistatic interactions between many mutations, and the effects of conditioning on past evolution, can one overcome the biases caused by choice of “wild-type” genome and sets of mutations around it? One system where this should be possible is with yeast for which a wide range of genomic technology has been developed and the ability to bring random combinations of mutations together is enabled by controlled mating. Empirical studies of epistasis have been made with *yeast crosses*, where two genetically distinct strains of yeast are mated repeatedly to generate a large number of offspring which are genetically “between” — in the sense of genetic distance — their two parents [4]. In this particular experiment, the two parental strains differ on 0.5% of their genome (roughly 3 *·* 10^4^ possible SNPs). The fitness of a large number of the created recombinant variants — upwards of 4 *·* 10^3^ in [5] — were assayed in a variety of environments. The resulting fitness measurements in each environment were combined with genotyping information to estimate the fraction of fitness variation between types that is explained by additive fitness effects: the unexplained variance is then a statistical estimate of the total epistatic contribution to the fitness landscape in that environment. The findings vary across environments; some have *>* 90% of the variance explained by an additive model, while others have closer to 50%.

Unfortunately, even from the very large number of yeast-crosses studied, and the very high-dimensional sub-space spanned by the set of mutations by which the parents differ, extracting information about the important statistical properties of the fitness landscape is problematic. One issue is that the distribution of pairwise genetic distances between variants tends to be narrow. In [4] the distribution of distances was peaked at 50% of the genomic distance between the ancestors with a standard deviation of *∼* 5%. Thus the data provide information primarily about the overall scale of the fitness landscape at one, particular, long distance. The crucial distance dependence is thus not explored. Another issue is parameterizing the distribution of the fitness landscape by a few simple summary statistics — here decomposing into additive and “unexplained” variance. While the models we have analyzed were, for tractability, assumed Gaussian, that is certainly not expected in nature and more information on *distributions* of fitness differences, rather than just covariances, is needed. (Indeed, we found this already in our analysis of mixed models with Gaussian epistatic parts and non-Gaussian additive parts for which the tail of the distribution was particularly important.)

Regardless of the difficulties of extraction of the most useful information, experiments that combine substantial numbers of mutations to make a great many combinations have the potential to tell us much about the statistics of genomic-scale — rather than protein scale —fitness landscapes. The ensemble of landscapes with Gaussian distance-dependent correlations that we have studied are, of course, far from realistic. But the goal of advancing general understanding — as well as the paucity of relevant data — mandates the exploration of caricature models as we have done. Our findings as to how the long-genetic-distance statistics can crucially affect intermediate term evolution, should provide additional motivation to develop high throughput means to probe the statistical effects of combinations of large numbers of mutations.

An advantage of using crosses of well-adapted strains to produce the subset of mutations to explore, is that effort is not wasted on large numbers of unconditionally deleterious mutations. Another approach would be to take a set of mutations that arose individually in evolution experiments and make many recombinants of these, either engineered or by crosses of evolved mutants that already differ by a substantial — but far less than the strain-cross experiments —number of mutations. Such biased sampling approaches have recently been experimentally realized [15, 60]. Explorations of the distributions of fitness differences as a function of genetic distance could be explored with either approach, as well as, crucially, how these vary with small changes of environment. As datasets become more sophisticated, one can start to develop better caricatures of fitness landscapes and seascapes using the data to guide development of models and insights from theory to guide experimental design. The time is ripe for more detailed back and forth between theory and experiment.

### 8.3 Theory beyond Gaussian landscapes

Studying adaptive walks on landscapes with distance-dependent statistics has given us considerable insight into the interplay between evolution and epistasis; however we are limited by the simplifying assumptions of the family of models, especially by the Gaussian assumption, which enabled us to generate landscapes on the fly and analyze the effects of past history via the response kernel framework. Real fitness landscapes are surely not Gaussian [38]; in the future one needs to study caricatures that include non-Gaussian statistics, especially the potential for anomalously large effect mutations that can arise even after much evolution has already occurred in the same environment, as seen during experimental microbial evolution [6]. In addition, to gain better understanding of the conditioning on past history, one needs to study different forms of time-dependent seascapes than the very simple exponential decay of temporal-correlations that we analyzed.

One way to go beyond the Gaussian simplification while keeping the desirable distance-dependent statistical structure would be to define the elements of the amplitude spectrum using other distributions. This is equivalent to transforming to the Fourier basis, then generating independent distributions for each Fourier mode. A natural extension of the Gaussian model would be Levy distributions; these would give a family of self similar random variables which are broadly distributed with power-law tails.

Seascapes can similarly be generated in the Fourier basis with each mode a stochastic process in time instead of a static random variable. Even within the Gaussian-correlated models, one can have a broad range of time scales with longer distance correlations of the landscape changing more slowly. The response kernel framework we have developed should enable progress to made on such models. More generally, one could consider dynamics that have more complex temporal structure than simple random walks, for example, having periods of relative tranquility punctuated by large, abrupt changes — loosely, a temporal analog of Levy distributions. This sort of model can be implemented within the framework described in Section 7.3.

Unfortunately, as soon as one goes beyond Gaussian randomness, one loses the ability to efficiently sample landscapes (or seascapes) on the fly as the evolution proceeds. In general, one needs to define 2*^L^* Fourier coefficients — intractable for numerical explorations. We propose that further progress can be made by defining *sparsified* models of fitness landscapes which keep only a small number of interactions while matching some of the believed-to-be important statistical structure of the landscapes. By analogy with sparse random matrices, where *O*(*N* log(*N*)) non-zero elements in an *N × N* matrix are sufficient to give the same eigenvalue spectrum as with all elements random, preliminary calculations suggest that at least the pair correlation structure of a Gaussian landscape, in particular a power-law correlated one, can be preserved with a number of Fourier coefficients polynomial in *L*. With this, our analytical framework could not be used, but numerical simulation would be computationally inexpensive to perform as with such a sparsified representation one could cheaply evaluate the landscape at any point in (genotype) space and time. An important question is what features of the sparsified landscapes determine the long-term evolution: even with Gaussian random coefficients, the rare regions into which evolution takes the population may be very different in a sparsified than a fully Gaussian landscape with the same amplitude spectrum.

With non-Gaussian statistics, many of the statistical features of adaptive walks will change, some quite dramatically. We have already seen that a broadly distributed additive piece greatly aids evolution. Walks in non-Gaussian landscapes will have different relationships between the past evolution and the distribution of available steps, as well as different conditions for how they get stuck for large but finite *L*.

### 8.4 Beyond weak-mutation strong-selection dynamics

We have in this paper considered only the simplest evolutionary process: a population that is small enough that mutations arise only infrequently, but large enough that drift is unimportant, especially that deleterious mutations cannot fix. Such a population takes a simple uphill adaptive walk. We end by discussing several important effects that are ignored in this simplified caricature.

Small populations do not need to evolve solely uphill because weakly deleterious mutations can drift to fixation with probability of order *e^−Nδ^* for a deleterious mutation of selective disadvantage *δ* in a population of size *N*. This means that the population never gets completely stuck. Its dynamics is essentially identical to stochastic motion on the landscape with a “temperature” of 1*/N*. There is a rich literature analyzing stochastic thermal dynamics on random landscapes [8, 9, 36] which would be instructive to extend to power-law correlated landscapes such as those we have studied. In spite of never getting fully stuck, the dynamics will continue to slow down, at least at low temperatures, but investigating how it does so, and how this depends on additional features such as long-tailed distributions of potential fitness steps, is a possible direction for future research.

Large populations are much more interesting. They can also escape local maxima but instead by “tunneling” through intermediate lower fitness genomes to reach a higher fitness one. This process involves only part of the population and its rate is a complicated function of the population size and fitness differences especially when the move to a higher fitness involves more than one intermediate lower-fitness step [72]. But the overall effect can be crudely approximated by allowing double (or triple, etc.) steps to find points of higher fitness. As the number of double net-uphill steps is much larger than single uphill steps when the mean available fitness, *µ*, is strongly negative, the effects of tunneling processes on the evolution can be large even before a local fitness maximum is reached.

But this is only one of the complexities of large population caused by the diversity engendered by multiple independent mutations occurring each generation. Complications include competition between mutants (clonal interference), as well as the fact that several mutations can occur on the same lineage before any fix. Large populations effectively explore many directions of the landscape in parallel. The rate of evolution is then limited primarily by selection rather than mutation. The competition that will make one of the directions eventually win is played out over a longer time scale — extensively studied for additive models [13, 18, 46] — and involves luck in both when and which several steps of further beneficial mutations occur.

Naively, being able to explore multiple directions in the fitness landscape before committing to any would suggest that, even if only beneficial mutations are allowed, large populations will increase their fitness more before getting stuck than small populations will. But there is another effect that goes in the opposite direction. Large populations are much greedier: the mutations that are likely to fix are much more heavily weighted toward larger effect — exponentially so – than in small populations [18, 22]. We have shown that greedier adaptive walks can get stuck at lower fitness local-maxima. Furthermore, experiments [62] and theory [31] have shown that there are at least some rapid adaptation scenarios in which larger populations fare worse at later times. More generally, which of the two competing effects — exploring multiple paths or being too greedy — dominates in determining when large populations get stuck is certainly a key question. Understanding the evolution on epistatic landscapes of large populations that are continually diversifying and pruning, is an important avenue for further research which should be enabled by the progress we have made thus far.

Beyond their interest for natural evolution, questions about how fast and how far evolution proceeds and how these depend on the nature of the evolutionary process, including strength of competition, sizes of population or multiple partially separated sub-populations, etc. are also of practical interest as evolution is being used in both bioengineering [44, 65] as well as machine learning [61] in order to optimize a wide spectrum of systems. Developing more general understanding of the interplay between the evolutionary dynamics of populations, and the statistical properties of fitness landscapes and seascapes is surely merited.

## 9 Acknowledgements

We thank Michael Pearce, Shamit Kachru, and Surya Ganguli for useful discussions and Michael Tikhonov, Oskar Hallatschek, and Joachim Krug for comments on earlier versions of the manuscript. This work was supported in part by the National Science Foundation via PHY-1305433 and PHY-1607606, and by a Stanford Bio-X Fellowship and a Pre-Doctoral Fellowship from Stanford’s Center for Computational, Evolutionary and Human Genomics both to AA.

### A Distance-dependent divergence functions

#### A.1 Divergence function derivatives

It is often useful to compute the correlations between the fitness differences induced by two sets of mutations in terms of the divergence function *D*. Recall that we have

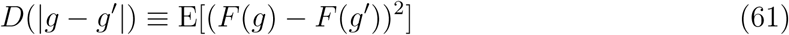

where *|g – g′|* is the number of sites where *g* and *g′* differ. Consider the pair of fitness differences *F* (*a*) *− F* (*b*) and *F* (*c*) *− F* (*d*), oriented so that the shortest path *a → b* runs in the same direction as *c → d*. Then we have

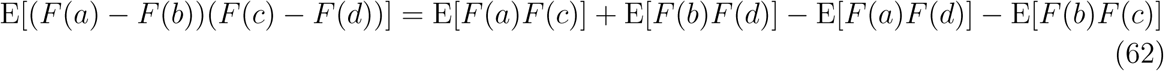

We can rewrite this correlation as

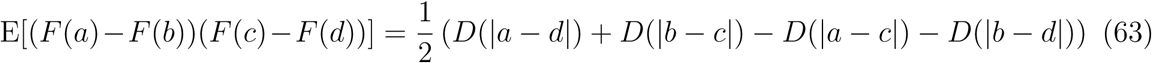

where we use the fact that E[*F* (*g*)^2^] is independent of *g*.

One case of interest is when *c* and *d* are adjacent, and the edge *c → d* is contained inside the line between *a* and *b*. Let 𝓁*_tot_*= *|a − b|* and 𝓁*_ac_*= *|c − a|*. Then we have

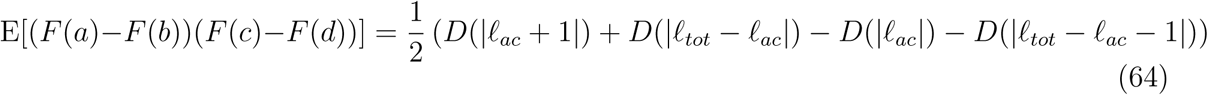

which in the continuous limit gives us

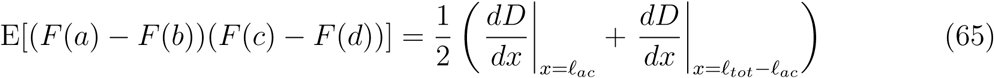

This tells us that the total fitness gain on a path is correlated with the fitness effect of a single mutation by 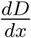 for a random path. For power law walks, we have

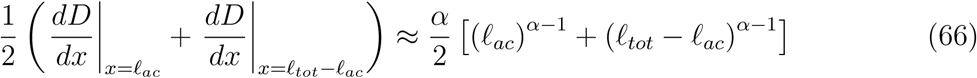

That is, for a random path the total fitness gain is most correlated with the fitness gains at the end, and least correlated with the fitness gains in the middle.

Another useful case is the correlation between the effects of random single mutations between two genomes a distance 𝓁 apart. In particular, we care about the correlation between two mutations on a single trajectory. Labeling genomes by their distance along a single path, we have

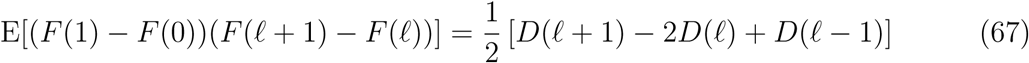

where *F* is indexed by steps along a path. We can approximate as

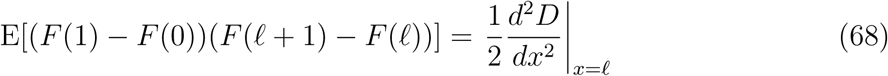

This approximation is good when *D* changes slowly with genetic distance. For sublinear *D*, 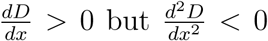 - giving negative correlations between mutational steps on the same path.

#### A.2 Computing divergence functions from amplitude spectra

One undesirable feature of the amplitude spectrum is that it depends on the total genome size *L*. We would like to be able to write the spectrum in a scale-invariant way so that we can understand what models look like in the large *L* limit. Accordingly, we define the *rescaled amplitude spectrum ρ_A_* by

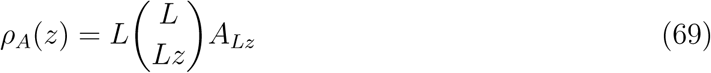

where *Lz* is rounded to the nearest integer. This normalization is chosen so that weighted sums over the amplitude spectrum can be converted to an integral. For example, consider a function *ψ*: [0, 1] *→* ℝ. The average of *ψ* weighted over the amplitudes *A_k_*of the Fourier modes (equal amplitude for all **K** with *|***K***|* = *k*) can be computed as:

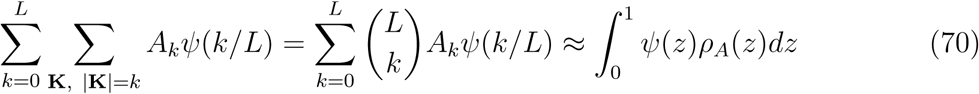

The amplitude spectrum has been described previously, but the rescaled limit — crucial for giving landscapes whose structure is well behaved in the large *L* limit — has, to our knowledge, not been studied previously.

For simple models we can compute the *A_k_* easily and therefore the *ρ_A_* as well. For the additive model, only terms at order 1 contribute; we have

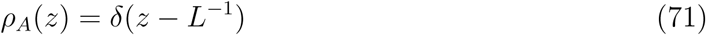

For independent fitnesses, each mode has equal weight (since it is a graded sum of independent random variables). The combinatorics of terms at each order sets the distribution; we have

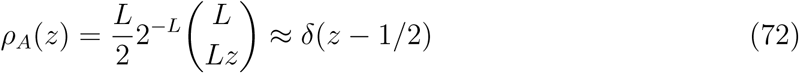

We can readily compute the divergence function *D*(𝓁) given *ρ_A_*(*z*). We begin by calculating the correlation E[*F* (*g*)*F* (*g′*)] between two genomes *g* and *g′* such that *|g – g′|* = 𝓁. We have

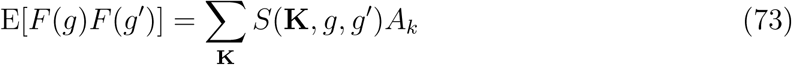

where

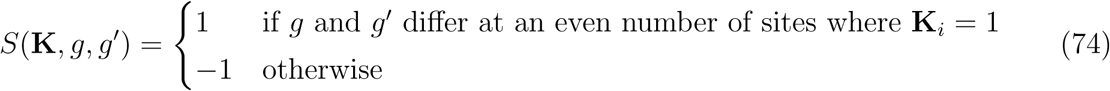

The sum over the **K** can be rewritten as

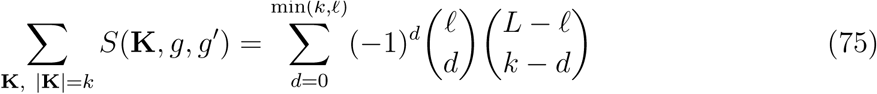

(Note that the terms on the right hand side are Krawtchouk polynomials - as first developed in [66]).

This sum cannot be evaluated exactly, but can be evaluated approximately in two cases. First, for *L, l ≫ k* — low order epistasis but macroscopic distances — we have

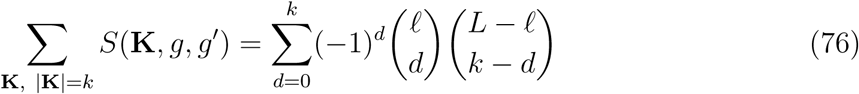

Use of Stirling’s approximation gives

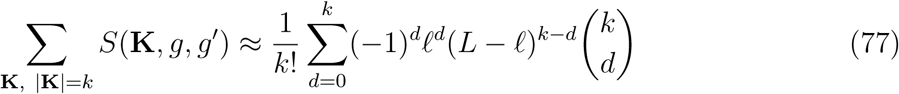

which can then be evaluated via generating functions as

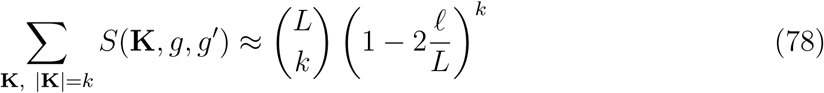

To evaluate the full sum with this approximation, we use the rescaled amplitude spectrum:

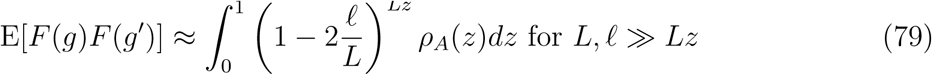

This approximation is useful if most of the weight of *ρ_A_* is at low orders. If terms of some particular lower order dominate, the correlations drop off polynomially, with fitnesses becoming uncorrelated at 𝓁 = *L/*2 and anticorrelated at 𝓁 = *L*.

The second and more useful case is when *L, k* ≫ 𝓁 — short-distances (compared to *L*, but arbitrarily high order epistasis. Then we have

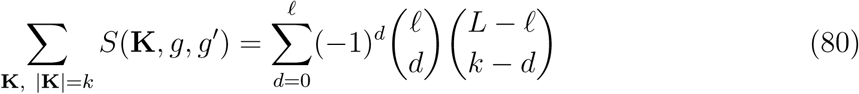

Stirling’s approximation then gives

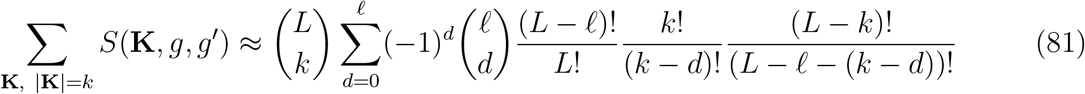

which simplifies to

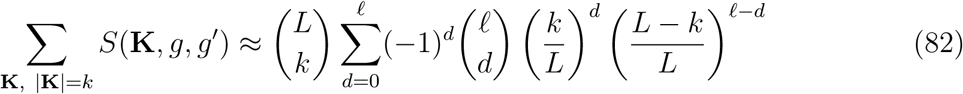

and a similar generating function computation gives

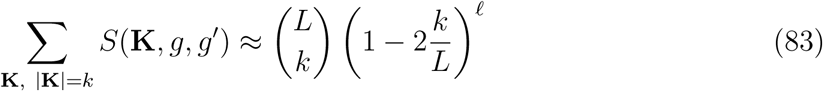

In terms of the rescaled amplitude spectrum we have

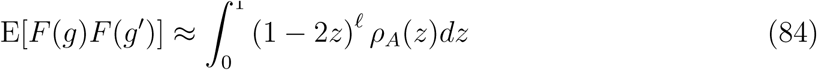

and therefore

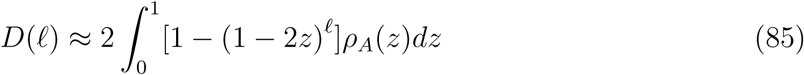

where we have rescaled so that *D*(1) = 1. Here we arrive at a nice intuition for how terms of different order affect the statistics of the landscape. Epistasis of order *Lz* contributes a term with exponential decay of correlations with lengthscale log(*|*1 *−* 2*z|*)*^−^*^1^. Terms of order less than *L/*2 contribute positively to correlations, while higher order terms contribute alternating signs on average.

Arbitrary combinations of different orders of epistasis can be represented in the large *L* limit, with 𝓁 ≪ *L* – the regime of interest for almost all we study in this paper — by the general scaled linear combinations of Equation 84.

#### A.3 Amplitude spectrum of power law landscapes

In order to have correlations with super-exponential long-range structure, *ρ_A_*(*z*) needs to diverge as *z* → 0. Specifically, consider power law divergence functions. In general, going from divergence functions to amplitude spectra is difficult; it requires inversion of the matrix *M* whose elements are given by

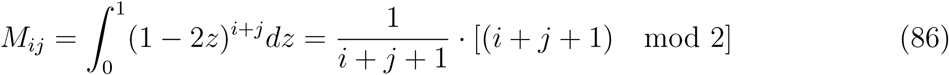

However, we can make an ansatz for the form of *ρ_A_*that corresponds to power law scaling of *D*(𝓁). We want a *ρ_A_*(*z*) such that

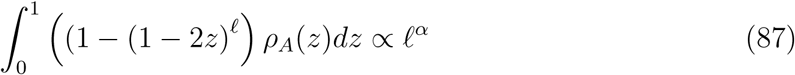

Differentiating with respect to 𝓁 gives us the condition

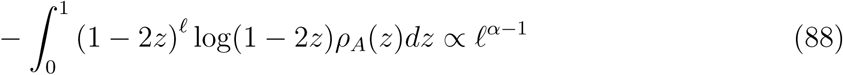

For large 𝓁, we expect the integral to be dominated by small *z*. The integrand, sans *ρ_A_*, peaks around *z* = *O*(𝓁^−1^), and has significant value over a range that is *O*(𝓁^−1^) as well. The value it takes is also *O*(𝓁^−1^). This suggests that *O*(𝓁^−2^)*ρ_A_*(*z* = 𝓁^−1^) = *O*(𝓁^α−1^) or

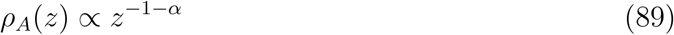

for small *z*. This is peaked at *z* = 0, and is consistent with the approximation that the integrand in Equation 88 is peaked around 𝓁^−1^. Note that the exponent of *ρ_A_*(*z*) is less than - 1. Therefore, power law distance-dependent divergence functions correspond to power law distributed power spectra, with most of the weight at small *z*.

Note that though *ρ_A_*(*z*) is weighted towards *z* = 0, for large *L* we can still have a significant weight at higher order (*k >* 1) Fourier modes. This suggests quantitatively that higher order epistasis matters increasingly for increasing genetic distances, at least in models that have a consistent large *L* limit (as all models defined by the divergence function do). We also again come to the lesson that there is some flexibility in the exact form of *ρ_A_*(*z*) with regards to its effect on *D*(𝓁); the form of the divergence at 0 sets the long tail behavior.

### B Response kernel approach

#### B.1 Dependence of DFE on past evolutionary history

A key property needed to understand evolutionary processes is the distribution of fitness effects (DFE) of all single mutants away from a given genotype. Consider a genome *g* on a distant-dependent correlated landscape, and let *s_i_* be the fitness gain from a mutation at site *i*. The *s_i_* are distributed as a covariate Gaussian, with zero mean, variance *D*(1) and correlation 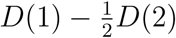.

We can write each difference *s*(*y*) as a sum of two Gaussian random variables:

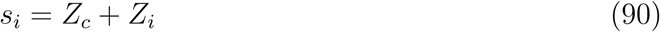

Here *Z_c_* is the “shared” part of the variability - a random variable with variance 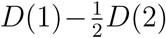 identical across all the sites. The *Z_i_* give the “private” part of the variability – Gaussian random variables with variance 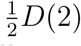, independent for all sites. This type of decomposition is always possible with identically correlated Gaussian random variables.

In the limit of large *L*, we can approximate *Z_c_* with the *average available fitness step*, 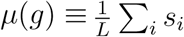. We have:

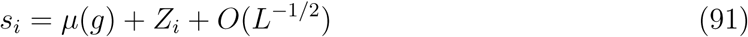

In this limit, the (empirical) DFE is approximately Gaussian with mean *µ*(*g*) and variance 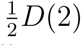. Since *D*(2) is a constant set by the landscape, once *µ*(*g*) is known, the entire DFE is known as well.

This gives us the following strategy for constructing adaptive walks:

- Draw the value of *µ*(*x*) from its distribution.
- Use the resulting DFE to take an evolutionary step.
- Define *x* + 1 as shorthand for the genome after the step is taken.
- Draw the next *µ*(*x* + 1) conditioned on past evolution and repeat
 where *x* labels the number of steps taken. Note that the DFE conditioned on *µ*(*x*) (and the previous evolution) is still Gaussian, even though the actual step taken *s*(*x*) is not. This simplifies the dynamics considerably, as each step in the evolution involves first evaluating a conditional Gaussian random variable, and then drawing from a simple distribution — the positive *s* part of the Gaussian DFE.

This suggests that in order to understand the dynamics along a single random adaptive walk, in the limit of large *L* all we need to understand the DFE at any point is its conditional mean *µ*(*x*) at that point. This approximation starts to break down when we ask about events which have probability *O*(*L^−^*^1^) in the Gaussian distribution; in particular, extremal values of the empirical distribution are no longer in the large number regime and need to be treated with care. But the continuous Gaussian approximation for the DFE holds so long as the number of possible adaptive mutations is large.

#### B.2 Deriving response kernels on power-law correlated landscapes

We now analyze how the DFE around the current genome is conditioned on the past history of the adaptive walk to that genome. Assume the walk has taken a sequence of steps, *y* = 1, 2*, …, x −* 1 leading to the current point labeled *x* with these steps having fitness changes *s*(*y*), each taken from the local DFE of average available fitness steps, with average *µ*(*y*). In this section we will describe how to compute the response kernels *J* and *K* such that

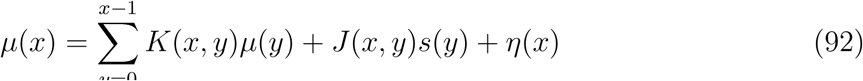

where *η*(*x*) is the additional random part of the DFE average: *η*(*x*) has mean zero and variance, *V_η_*(*x*). We make the continuous approximation for the sums:

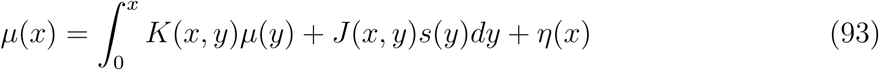

Note that the *J* and *K* are non-zero only for *x > y*.

For most cases it will suffice to use this continuum approximation; but there are a few special cases of landscapes with particular divergence function *D*(𝓁) which are more easily understood with the discrete equations.

It is useful to define and use the following correlation functions in order to obtain self-consistent equations for *J* and *K*:

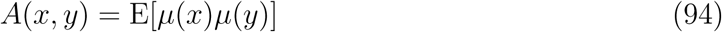

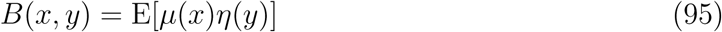

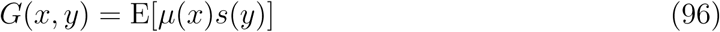

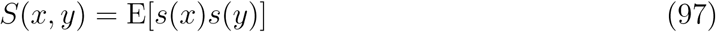

In addition, we define the convolution operator *∗* by

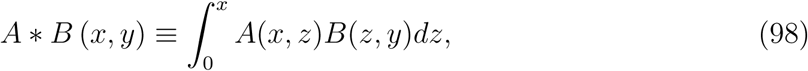

and the transpose, T, by

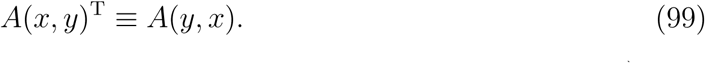

(In terms of the discrete sums, these are simply matrix multiplication and transpose.)

By explicitly computing *A*, *B*, and *G* using Equation 93 multiplied by on of *µ, s*, or *η* at a different point on the path, and averaging, we get:

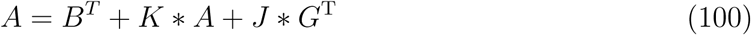

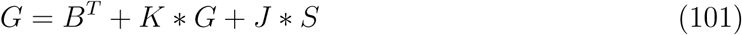

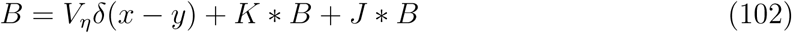

Note that *K* shows up once in each equation, convolved with the quantity on the left hand side. With some foresight, we rewrite *K* = *δ* + *P* to cancel the left hand sides. In terms of *P* we are left with:

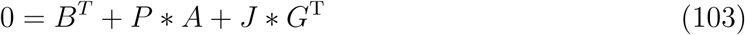

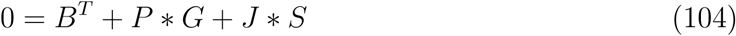

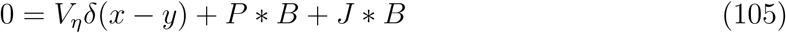

Up until now, the equations are exact (and also hold for the discrete sums). We now take the large *x* limit keeping terms to leading order in powers of 1*/x* and similarly for *y* and *x − y*. We now define a useful quantity related to the second derivative of *D*(𝓁) in the large 𝓁 regime:

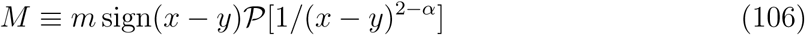

with *m* a positive coefficient given by the normalization of *D*, and the principal part, 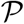, ensuring that 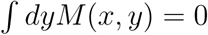. Then in the large *x* limit, from the definitions we have:

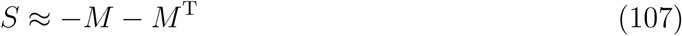

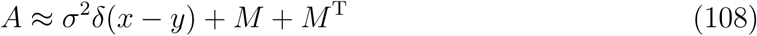

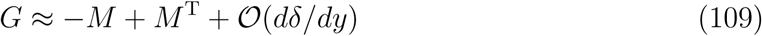

with *σ*^2^ proportional to *V_η_*. Corrections to all of these correlations are smaller by one power of 1*/x* except for *A*: the symmetry under the transpose means that there is only a second derivative of *δ*(*x − y*) correction to the *δ* function part of *A* (like a correction of *O*(1*/x*^2^)).

We now make the ansatz (readily shown to be correct) that *J* scales as *x^ν−^*^1^, with 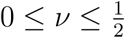, and that *P* is much smaller for large *x*. Then from Equation 105 we have

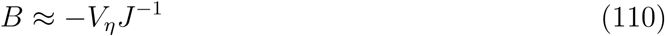

with inversion taken over *∗* (like matrix inversion). In Equation 104, the *P ∗ G* term is negligible and in Equation 103, *P ∗ A ≈ σ*^2^*P* plus smaller corrections. Writing an expression for *B*^T^ and substituting, we get

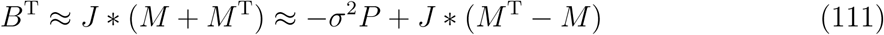

which can thus determine *J* and *P* = *−*2*J ∗ M/σ*^2^, subject to conditions that they are only non-zero for *x > y*, as is *B* – the latter not automatic. Counting powers of *x* we see that *B ∼ P ∼ x^−^*^1^*^−ν^* and, from the form of *M*, that must have

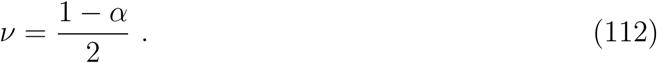

One can directly get the scaling behavior and this result for *ν* by considering the behavior for (*x − y*) *≪ x*. This can be obtained straightforwardly by approximating the kernels as functions of only *x − y*, taking *x* → ∞ and then Fourier transforming and decomposing functions into sums of parts that are analytic in upper and lower half-planes corresponding to being zero for either *x < y* or *x > y*. Armed with these forms (and some knowledge of such singular integral equations), exact solutions can be guessed:

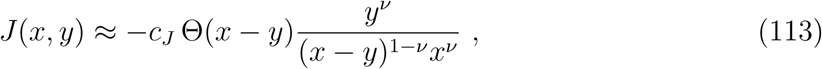

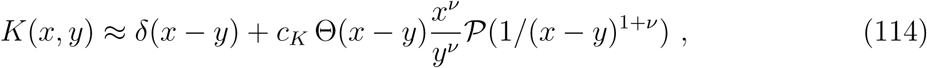

and

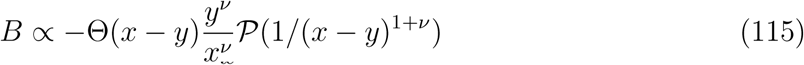

with *c_J_* and *c_K_*positive coefficients that can be written in terms of Beta functions and combinations of *V_η_*and *σ*^2^ to set the scales, and Θ(*z >* 0) = 1 while Θ(*z <* 0) = 0. Note that with the correct value of *ν* — indeed only with that value of *ν* — the *x > y* part of *J ∗* (*M* + *M^T^*) = *B^T^* vanishes as it must.

The sign of *P* is rather misleading due to the principal part. If the convolution integral is done by parts, we have that

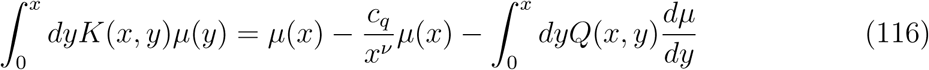

with the kernel for 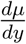,

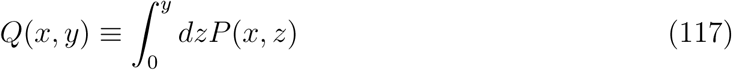

and *Q* and the coefficient *c_q_* both positive. Thus the effects of *K* are: *δ*(*x − y*), a negative — but needed — correction to this, and a negative convolution with *dµ/dy*, the latter two being the same order for large *x*.

In order to show that the solutions to the integral equations are of the form given, two integrals need to be done. To show that *B* is proportional to the inverse of *J* — i.e. that 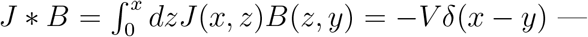 one only needs to note that the *z* dependent parts of the *z^ν^/x^ν^*and *y^ν^/z^ν^* factors from *J* and *B* cancel. To do the integrals *J ∗ M* and *J ∗ M^T^*, the substitution *z* = *xy/*(*x* + *y − ζ*) changes the integral over *z* from 0 to *x* to an integral over *ζ* from *−∞* to *x*, eliminates the *z^ν^* part from *J*, brings out (combining with the *x^−ν^* from *J*) an overall factor of *x^ν^/y^ν^*, replaces *x − z* and *y − z* with *x − ζ* and *y − ζ*, and cancels all factors of *x* + *y − ζ* thereby making the integrand the simple standard form (*x − ζ*)*^ν−^*^1^(*ζ − y*)*^−^*^1^*^−^*^2^*^ν^* and the integrals over the different parts of the range of *ζ* and for *x > y* and *x < y* all simply expressible in terms of Beta functions. The structure is now exactly the same as it would be if the limit *|x − y| ≪ x* were taken initially and Fourier and Wiener-Hopf analysis used. Note, however, that for *µ*(*y*) and *s*(*y*) powers of *y*, as we find, the contributions from the kernels are dominated by intermediate range of *y/x* and thus the approximation *x − y ≪ x* is incorrect by multiplicative constants, but gives the correct scalings with *x*.

The theoretically predicted scaling forms match up well with our numerics. One way to see this is to plot *J* and *K* as functions of (*x − y*)*/x*, for various walk lengths *x*. Theory predicts that if *J* is scaled by *x*^1^*^−ν^* and *K* by *x*^1+^*^ν^*, the response kernels should collapse onto a single, universal scaling form. Figure 14 shows these rescaled response kernels. We can see that both *J* and *K* collapse as expected. The shape of the theoretical curves at large *x* (dashed lines) matches the actual universal scaling form inferred from the numerics.

**Figure 14:**
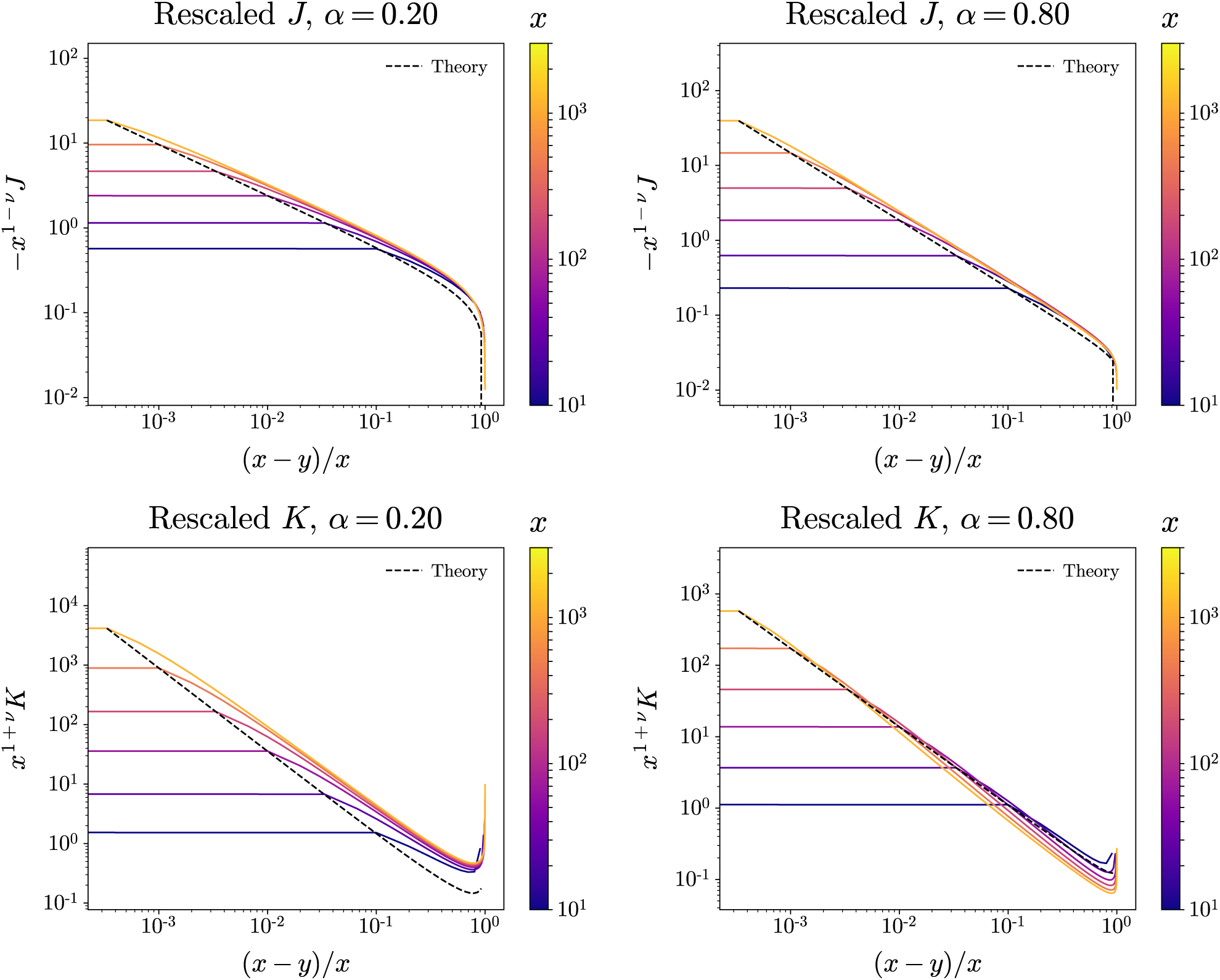
Rescaled *J* (*x, y*) and *K*(*x, y*) for various walk lengths, *x*. Genetic distance *x – y* is rescaled to range [0, 1] by dividing by *x*, and *J* and *K* are rescaled by predicted powers of *x* to compensate. Rescaling collapses response kernels. Theoretical scaling form for large *x* (dashed line) captures power law regime well. Largest deviations from the theoretical asymptotic forms are: *J* for small *α*, and *K* for *α* close to 1.

The largest deviations are for *J* with *α* near 0, and for *K* with *α* near 1. This is to be expected; these are when the power law regime falls off the most slowly for each, and the corrections are expected to be the largest. Regardless, the basic power law character of the response kernels at intermediate scales seems to hold; this intermediate range behavior is what dominates the dependence of evolutionary dynamics on past evolution. These numerical results support our use of the theoretical scaling forms to compute the typical evolutionary trajectories.

#### B.3 Adaptive walks on exponentially correlated landscapes

Here we analyze adaptive walks on landscapes with divergence function

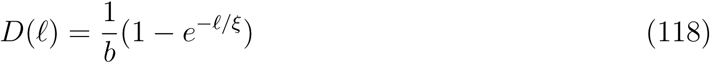

where *b* = 1 *− e^−^*^1^*^/ξ^* normalizes so that *D*(1) = 1 for all *ξ*. That is, the fitness of genomes at a distance 𝓁 ≪ *ξ* are highly correlated, and the fitness of genomes at long-distance 𝓁 ≫ *ξ* are close to statistically independent. In particular we are interested in the case where *ξ ≫* 1, which arises from the *NK* model.

The exponential correlations give the advantage that the response kernels can be computed exactly. We want to know the conditional distribution of the fitness at a distance *x* along the walk. It is easiest to compute in terms of *F* (*x*) along the path, where *F* is the absolute fitness function (well-defined since *D*(𝓁) is bounded). For arbitrary genomes *g* and *g′* we have

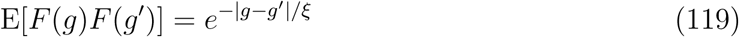

where for convenience we have normalized the landscape so that the absolute fitness at any point has 0 mean and variance 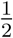 (instead of with *D*(1) = 1 as in the main text). On any spath, we can compute the conditional mean of the fitness at the end of the path *F* (*x*) in terms of the previously observed fitnesses as:

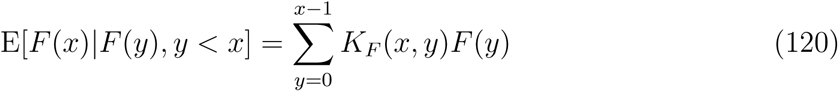

where *K_F_* (*x*) obeys the equation

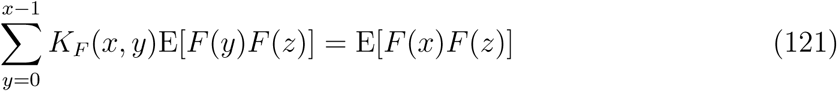

for all *y* on the path. We can solve for *K_F_*directly. Substitution gives

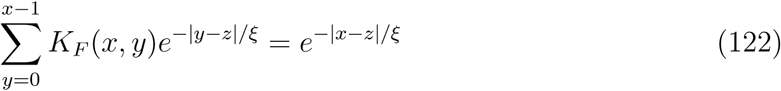

Solving for *K_F_* (*x, y*) for all *z < x* we get

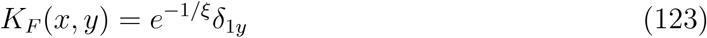

That is, the final fitness *F* (*x*) is correlated directly only with the previous fitness *F* (*x −* 1) with correlation *e^−^*^1^*^/ξ^*.

Evolution in an exponentially correlated landscape is in some sense a Markov process; only the last fitness value reached contributes to the statistics of the current neighborhood. The correlation coefficient *e^−^*^1^*^/ξ^* has the right limits; when *ξ ≫* 1, the landscape is effectively additive on short distances and the correlations vanish. When *ξ ≪* 1 the landscape is of the independent type and the average step available is exactly *−F* (*x*).

Though we have computed correlations in terns of the absolute fitnesses *F* (*x*), in exponentially correlated landscapes this gives complete information about *µ*(*x*). We can see this by computing the simple conditional variance Var[*F* (*x*)*|µ*(*x*)]:

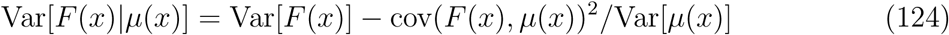

We can compute 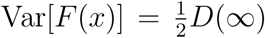 (since long range correlations vanish), 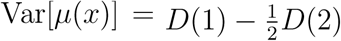, and 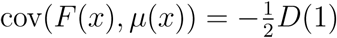. Combining, we have

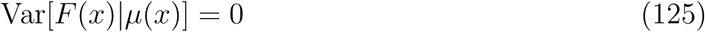

Therefore we have:

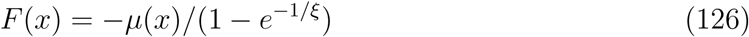

One can think of this as an extension of the situation in the independent fitnesses case. There, since 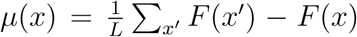. where *x’* are all one-step neighbors of *x*, independence and the central limit theorem gives 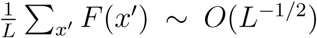, and we have, for large *L*, *µ*(*x*) = *− F* (*x*).

The combination of Equation 126 as well as *s*(*x*) = *F* (*x*) *− F* (*x −* 1) cause knowledge of the *F* (*x*) to be equivalent to knowledge of the *s*(*x*) *and µ*(*x*). Therefore Equation 123 can be used to compute the response kernels *J* and *K*. However, there is an additional complication: there are 2*x* total *s* and *µ* for a walk of length *x*, but only *x* total values of *F*. This redundancy leads to zero-modes in the *s − µ* covariance matrix, and a degeneracy of degree *x* in the response kernel.

Using this degeneracy, we can write the response kernel in a variety of ways. The two most interpretable are:

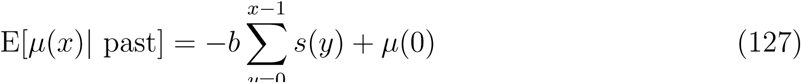

and

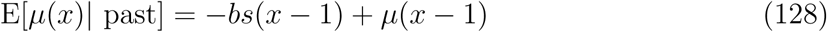

where *b* = 1 *− e^−^*^1^*^/ξ^*, and we normalize so that *D*(1) = 1 as in the main text. The first form suggests a flat weighting over past *s*(*x*) with an additional dependence on *µ*(0) — likely not important for long walks for which the first term continues to grow. The second form shows dependence only on the immediately preceding properties of the adaptive walk. The factor *b* accounts for the fact that the landscape is somewhere between additive and independent. For *ξ ≫* 1 (so that have almost additive effects of fitness steps), *b* → 0 and the dependence is only on *µ*(*x*), which is also 0 in that limit, giving E[*µ*(*x*)*|* past] = 0 as expected. For *ξ ≪* 1 (effectively independent fitnesses), *b* → 1 and we recover the response kernel of the behavior of the independent landscape.

### C Statistics of random and evolution-conditioned maxima

#### C.1 Local maxima

The expected number of maxima on a random fitness function on the hypercube depends on the local correlation properties only. We can compute this expectation in the following manner. Suppose that the fitnesses in some neighborhood of a genotype (ie that genotype and all single mutations away) are covariate Gaussian. Let *s_i_* be the fitness benefit of a mutant at site *i*. We assume that the *s_i_* have 0 average, variance 1, and identical correlation *r*. In the limit of large *L*, the {*s_i_*} are independent given their average *µ*(*g*). The distribution of *µ*(*g*) is a Gaussian random variable with mean 0 and variance *r*.

In the limit of large *L* the empirical distribution of the {*s_i_*} is also Gaussian with average *µ* and variance 1 *− r*. Note that *g* is a local max if all *s_i_* are negative, which happens with probability

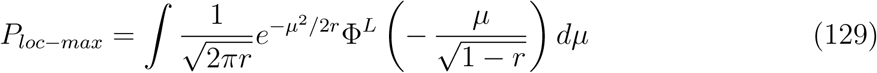

with Φ the error function. We use a saddle-point approximation to compute *P*. The log of the integrand *φ*(*µ*)

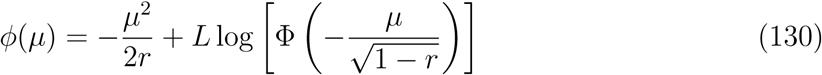

Differentiating gives

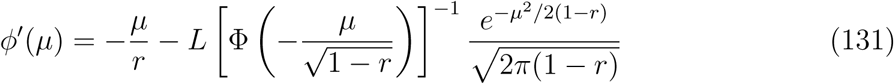

The peak is at *µ* = *µ^∗^* determined by:

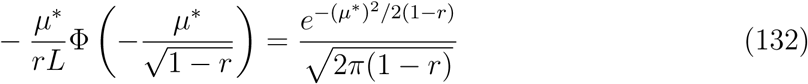

Clearly, *µ^∗^ <* 0. If 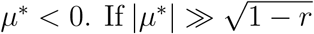, then 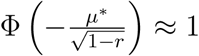 (leading correction 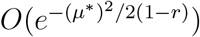). In this approximation, we have And then, approximately

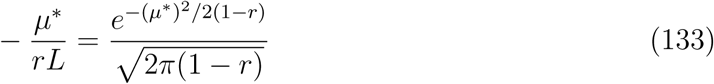

And then, approximately

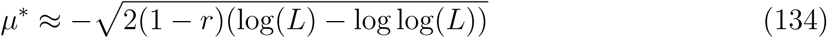

We can use this approximation to calculate 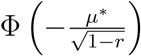. We need this calculation to be accurate to 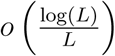 in order to compute *P* to logarithmic accuracy in large *L*. Error function asymptotics give

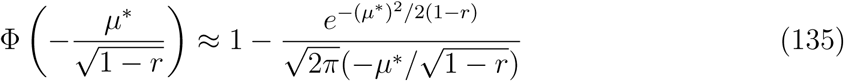

Substitution gives

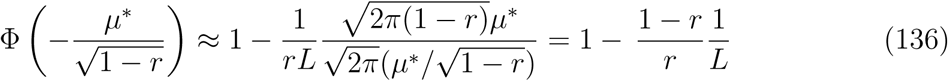

The second derivative of the log integrand *φ* is

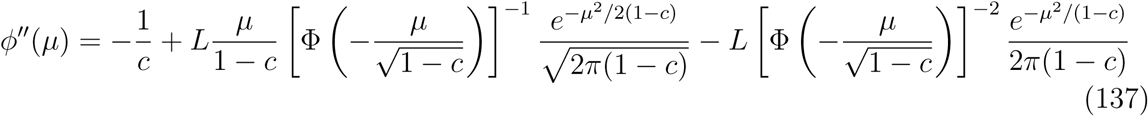

The value at *µ^∗^* is

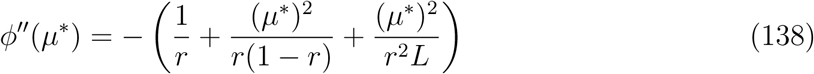

which is dominated by the middle term and is approximately

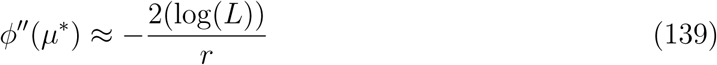

where we drop the log log(*L*) term as it doesn’t contribute to leading order in log(*L*).

The saddle point approximation then gives us

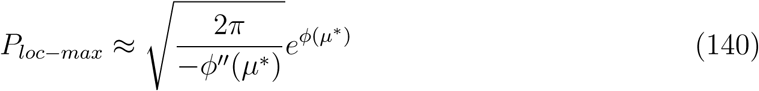

We can evaluate this to get

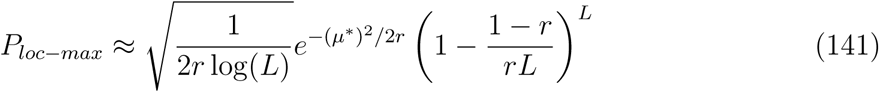

For large *L*, substituting in *µ^∗^* gives us

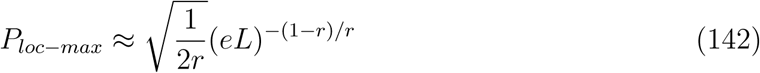

up to poly-log factors. Note that for the independent fitnesses model, 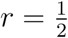, and 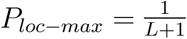, matching the approximate calculation up to an *O*(1) factor.

The result for *P_loc−max_* implies that if *r* has some limiting form for large *L*, the probability of a single genome being at a local max is some power of *L*. The total number of maxima thus scales roughly like *L^−^*^(1^*^−r^*^)^*^/r^*2*^L^*. As the correlation between step sizes decreases, the number of maxima does as well; the limit *r* → 0 corresponds to the additive model and there is a single local max on the whole fitness landscape (so *P_loc−max_* = 2*^−L^*).

Note that for power-law correlated fitness landscapes, *r* does not determine the long-distance dynamics of random adaptive walks. There are fitness landscapes with very different local structures which nonetheless have similar long-term evolutionary dynamics. In the limit of large genome size, *L*, the dynamics is dominated by the long-distance behavior of *D*(𝓁), which gives rise to the tails of the response kernels. Classifying the “ruggedness” of landscapes via expected properties of maxima is therefore not very useful for understanding evolution on epistatic landscapes.

#### C.2 Maxima reached by uphill random walks

We can estimate the max fitness increase reachable by an uphill-only random walk. First consider the independent fitnesses landscape. The cumulative distribution function, CDF, of an uphill walk of length *X* is given by Φ(*F*)*^X^*. Let the number of uphill paths of length *X* be given by *N_U_* (*X*). If the uphill paths are statistically independent, the maximum fitness attainable, *f*_max,up_, could be approximated by solving

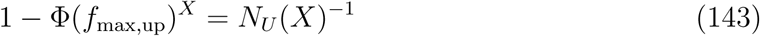

Correlations due to shared subpaths will cause this approximation to overestimate *f*_max,up_. On the hypercube, we have 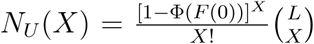. Solving for the CDF approximately, we get

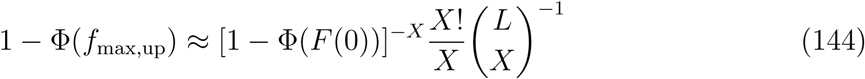

The maximum is dominated by the combinatorics of the number of paths, rather than the structure of the paths themselves. If Φ(*F* (0)) is not too close to 1, the right hand side is minimized at 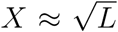, giving us the optimal CDF value 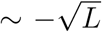, which gives a max fitness gain of

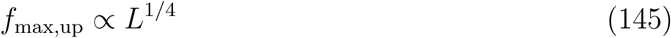

in the independent Gaussian case, compared to the global max of *O*(*L*^1^*^/^*^2^). Uphill paths have trouble reaching the global max, and random adaptive walks do not go nearly as high. The only way to go higher would be to allow some deleterious mutations as well (which we will not analyze here).

Performing calculations as above for correlated landscapes is more difficult and we will not attempt it here. However, we expect interpolation between the independent fitnesses and additive cases. The maximum absolute fitness 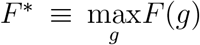 on the landscape can be crudely estimated as

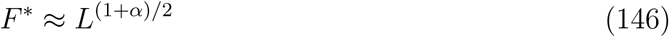

using the observation that the fitness difference between any pair of points at a distance *x* is *∼ x^α/^*^2^, and there are roughly exp(*cL*) “independent” pairs of points we can choose. We expect something similar to the independent fitnesses case, where *f*_max,up_ scales as a power of *L*, but not as much as *F ^∗^*. As before, to reach the global maximum one must have an extremely fortuitous choice of starting location. However we will see in the next section that small perturbations of the long range statistics can drastically increase chances of reaching points near the global maximum fitness.

#### C.3 Mixed landscapes with narrowly distributed additive piece

A small additive piece of the fitness function can have large effects on the global dynamics of evolution. The net effect on the dynamics is also sensitive to the distribution of the additive part. To show this concretely, we work with mixtures of a power law and an additive piece and consider different tails for the additive piece.

Let *s_C_* be the steps in the correlated part of the landscape, and let *s_A_* be the steps in the additive part so the total fitness gain is *s* = *s_A_*+ *s_C_*. We first consider an the additive part normally distributed with variance 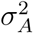. The distribution of *s_C_* and *s_A_*is jointly Gaussian. The conditional means can be computed directly:

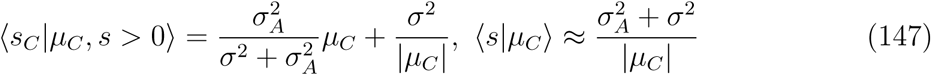

for 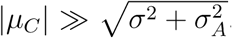. For 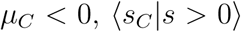 is reduced compared to what it would be in the absence of the additive piece. This change in the relationship between *s* and *µ_C_*changes the dynamics, making it easier to find uphill steps.

We can use the above equations to compute the dynamics on a power law plus additive landscape. From Equation 29 we have, schematically

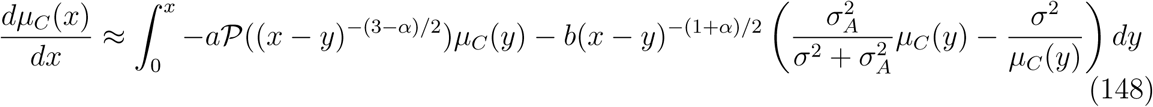

If *µ_C_* (*x*) and *s_C_* (*x*) saturate, then we know *s*(*x*) must saturate to 0 (since *J* (*x*) has a divergent integral). We solve for the equilibrium 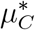 by setting the 2nd and 3rd terms equal to each other. If E[*µ_C_* (*x*)] is constant for all *x*, those two terms will be much larger than the 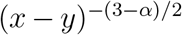 term. Define 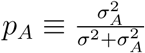 to be the fraction of the landscape that is additive. This gives us the saturating value 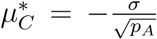. We can find the approximate length *X* at which saturation occurs for small values of *p_A_* by using the relationship E[*µ_C_* (*x*)] *∝ −σx*^(1^*^−α^*^)^*^/^*^2^ for the power law walk. We have

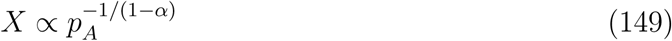

After the walk reaches saturation, we have E[*s*] = 0 and the saturated total step *s_sat_* goes as 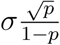 Thus E[*F* (*x*))] goes like a power law for roughly *p_*A*_^−^*^1^*^/^*^(1^*^−α^*^)^ steps, after which it goes roughly linearly at a rate 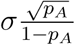 per step.

We can also compute how far up the walk is expected to go for finite *L*. We can find the minimal value of *p_A_* for which the walk length is *O*(*L*) for fixed *L*. This occurs when 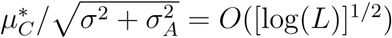, which occurs when

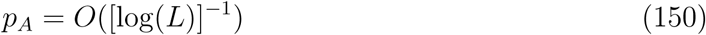

For large enough *p* the population gains fitness linearly with step size; for *p_A_* smaller than the critical value, the power law correlations perform better as they still have *f* (*x*) *∼ x*^(1+^*^α^*^)^*^/^*^2^. This critical value goes to zero as *L* increases (albeit slowly). The dynamics goes similarly for more narrowly distributed *s_A_* as well.

For Gaussian and sub-Gaussian distributions, the primary effect of the additive piece is to let *s* relax to 0. The adaptive walk “chooses” to use the additive component of the landscape as the primary source of its adaptive steps, while not moving much in the correlated landscape. A balance can be found where *µ_C_*does not change; this means that in the correlated landscape, the trajectory goes along a path of constant curvature. This small additive piece can have a large global effect on evolutionary dynamics.

The transition to *O*(*L*) fitness gains due to the additive piece has been previously seen in the case of RMF models (additive mixture with *α* = 0). Previous work has shown phase transitions from linear to sublinear fitness gains of random adaptive walks as the amount of additive piece is changed in the limit of infinite *L*. [49, 56] The phase transition occurs at *p* = 0 for sub-exponential tails of the random piece if the additive piece is constant. These new results explicitly show the crossover for finite but large *L*, showing the scale of the amount of additive piece needed to go long distances decays slowly with *L* if *s_A_* is narrowly distributed.

Even in cases where the additive component does not change the “typical” dynamics, it may change extremal ones. Consider the case of an additive, narrowly distributed component plus a correlated component. Now suppose we wished to find the “best” uphill path starting at a particular node. If we had full knowledge of the landscape, one simple strategy would be the following: for every step, find a direction where *s ≈* 0 and *s_A_≈ σ_A_*. If we can easily find such a step, is has *s >* 0 and also ensures that *µ_C_* will not increase. Then, at the next step we should be able to easily find such a step again.

So long as *σ_A_ ≫ L^−^*^1^, we should be able to find a step in the correlated landscape such that *s <* 0 and *|s| < σ_A_*. This means that the critical *p* for the existence of at least one path which gets close to the global maximum is at most *L^−^*^1^, as opposed to the [log(*L*)]*^−^*^1^

#### C.4 Mixed landscapes with broadly distributed additive piece

In the main text, we considered the case where large steps were relatively plentiful; however the regime where the probability of a large step is *O*(*L^−^*^1^) has some additional structure as well.

Consider a simplified model of a correlated plus additive landscape, where with probability *p_add_* any single uphill direction has an arbitrarily large uphill part. When *p_add_* = *n_add_L^−^*^1^ for some *n_add_* = *O*(1), there is some waiting time to find the uphill direction. If the probability of any uphill step on the correlated landscape alone is *p_cor_*, then an uphill step using the additive piece occurs with probability 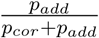, or every 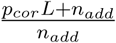 steps.

The dynamics then proceeds as follows: the population follows its normal trajectory on the correlated landscape until it finds and picks a large uphill direction. It then takes the large uphill direction, and takes a step *µ* in the correlated landscape. On average then, this leads to the following equilibrium equation for *µ*:

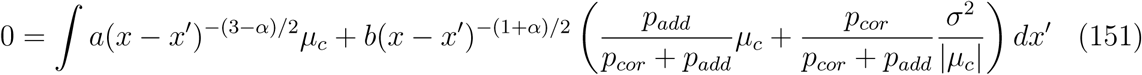

where 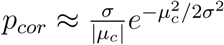. The equilibrium condition is given when the second and 3rd terms cancel; we have

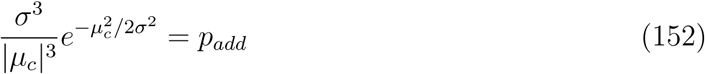

which gives us 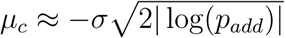, or *p_cor_*= 2*|* log(*p_add_*)*|p_add_*. For small *p_add_* (as we expect in the crossover regime), *p_cor_≫ p_add_*.

The walks are of extensive (linear in *L*) length when *Lp_cor_* = *O*(1) (more accurately, when 1 *− Lp_cor_* = *o*(*L^−^*^1^)). Therefore, to have dynamics of this type that nonetheless go a long-distance, we need *|* log(*p_add_*)*|* = *O*(1) - guaranteed when *p_add_* = *O*(*L^−^*^1^). A necessary condition for this ansatz to hold is that the additive piece gets used before the correlated piece would normally get stuck; that is we need 𝓁_cor_ ≫ *l_add_* where

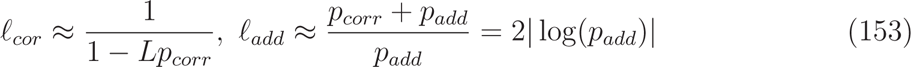

which means that walks in the correlated bit must go at least length log(*L*) for this method of equilibration to work.

This again shows us that the global and sometimes extremal statistics of the landscape matter far more than the local ones. Different extremal statistics lead not only to different crossovers in behavior, but potentially to different phases as well.

#### C.5 “Very abstemious” walks

To better understand the geometry of the fitness landscape, it is useful to know what uphill paths exist (or are likely to exist), and how these might be found by local “decisions”. In Section 5.1, we considered walks with large *L* and fixed *q*. What happens when we choose very small *q* so that the smaller the step the more likely it is to be chosen? More precisely, what happens when *q* scales with *L*? As there are at most *L* possible steps, *q* canot be smaller than *∼ L^−^*^1^; to understand the behavior we focus on the case where *q ∼ L^−ζ^*, for some positive *ζ <* 1.

If, for a slow, steady walk the number of steps, *x*, starts to become of order *L*, the tree approximation we have been using breaks down. A major complication is then that some fraction of possible steps — of order *x* out of *L* — would reverse mutations that have already occurred. This changes the correlations, and the response kernel forms we have derived no longer obtain. But there is a simple way out: if we only consider mutations that have not already occurred, then the symmetry under permutation still implies that the *L−x* allowable next steps will still have a distribution well characterized by the now-restricted *µ*(*x*) as long as *L − x* is large. Thus we should be able to take a total number of steps, *X*, at least some fraction of *L*.

For very small *q*, *µ*(*x*) will remain small until large *x*. The distribution of available *s* is thus not far from symmetric and a positive *s* can be chosen with typical value of order *q* (as long as *q ≫* 1*/L*). The contribution from the *s*(*y*) history to the conditional mean, *µ*(*x*), is then simply *−qx^ν^*. For Equation 29 to be true the effects of the *s*(*y*) history must cancel the effects of the *µ*(*y*) history, so Equations 31 and 32 require that *−µ ∼ qx*^2^*^ν^*. The walk can continue in this way until *−µ*(*x* = *X*_1_) *∼ σ* which occurs for *X*_1_ *∼* 1*/q*^1^*^/^*^2^*^ν^* at which point the fitness gain has already reached

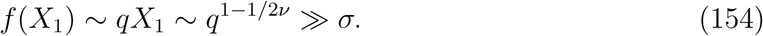

We can now choose *q* just small enough that *X*_1_ = *O*(*L*) by choosing *q ∼* 1*/L*^2^*^ν^* = *L^−^*^1+^*^α^* (which is larger than *O*(*L^−^*^1^) as required). Then the walk has already reached *f* (*X*_1_) *∼ L*^1^*^−^*^2^*^ν^* = *L^α^*— a remarkably good outcome!

This walk can be continued into the regime where *−µ ≫ σ*. To do this while keeping the total length *X* of order *L*, one needs

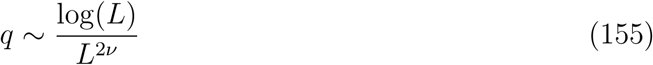

and the walk then reaches a height

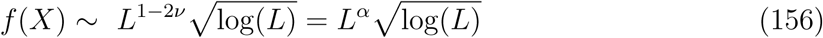

as quoted in the main text. Note that the logarithmic factor should really be log(*qL*) as *qL* is the size of the pool of neighbors with *s* in the bottom *q* fraction of the positive ones. But as log(*qL*) *≈* (1 *−* 2*ν*) log(*L*) this only decreases the maximum fitness reached by an order one factor, and the walk could keep going a bit longer by loosening the restriction of *q*. However it will still be highly restricted by conditioning on the past and we expect that not much extra can be gained by gradually relaxing the *q* restriction: this could be analyzed by our methods but as it would at best change the coefficient of the bound, we do not do so.

### D Dynamics on static and time-varying landscapes

#### D.1 Time-dependence of power law walks (fixed landscape)

We assume a model where mutations arise from a Poisson process, with characteristic time *τ_M_*= 1 between mutations. Deleterious mutations are purged instantly; adaptive mutations fix instantly.

Under this model, the time *τ_U_* (*x*) to take the *x* + 1 th step is distributed exponentially with mean time 1*/p_U_*(*x*), where *p_U_* (*x*) is the probability of any given step being uphill. Note that in the large *L* limit, *p_U_* (*x*) only depends on the current average available fitness gain *µ*(*x*), as the variance of available steps (given *µ*(*x*)) is constant. For *|µ|/σ ≫* 1, we have approximately

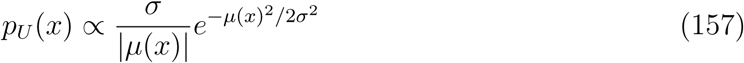

This means that *τ_U_* (*x*) is super-exponential in *µ*(*x*) — the adaptive walk is very slow.

Since *µ*(*x*) has variation, *τ_U_* (*x*) is broadly distributed across different walks. In fact, log(*τ_U_* (*x*)) is roughly normal. We can decompose its variance as

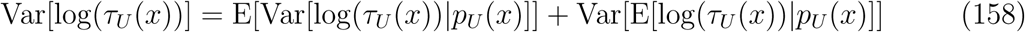

where the average is taken over all uphill walks of length *x*. From the convexity of log, we have approximately

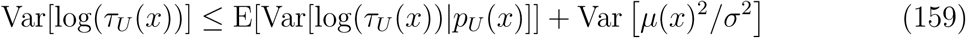

for large *µ*(*x*). Since *τ_U_* (*x*)*|p_U_* (*x*) is exponentially distributed, Var[log(*τ_U_* (*x*))*|p_U_* (*x*)] is small compared to the variability from *µ*(*x*) and the second term dominates. We expect, then, that log(*τ_U_*(*x*)) is distributed similarly to *µ*(*x*)^2^; its mean and variance should both scale as *x*^1^*^−α^*. (Note that we expect a similar result when considering the distribution of *τ_U_* (*x*) conditioned on the path itself; this is relevant for the analysis of time-dependent landscapes in Section 7.2.)

Figure 15 shows the exponents of power law fits to the mean and standard deviations of E[log(*t*(*x*))], averaged over different walks. These exponents show good agreement for low *α* where the approximate arguments above are expected to hold. They deviate at high *α* where *µ*(*x*) grows slowly enough that neglected terms matter more (at least for the range of *x* explored).

**Figure 15:**
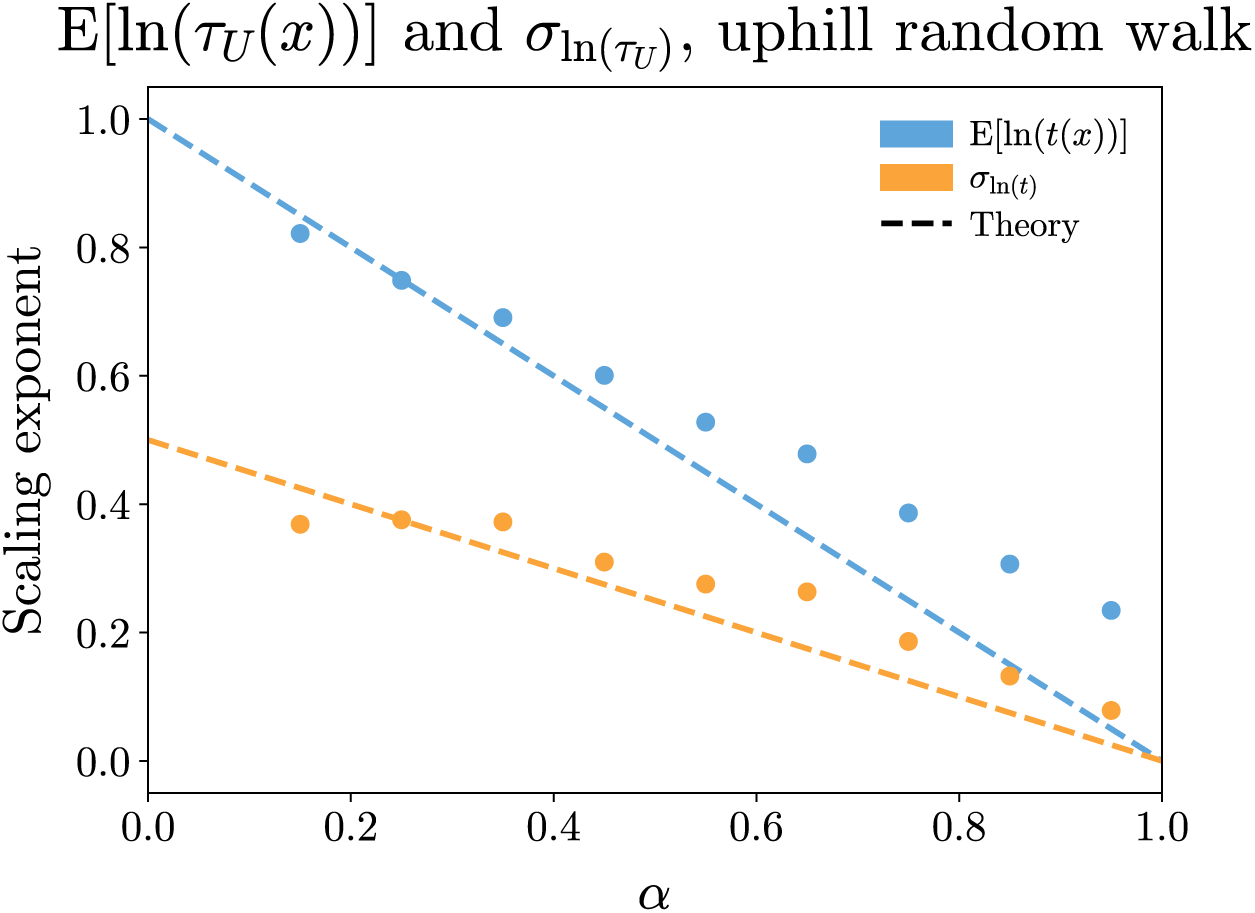
Average log derivatives of mean and standard deviation of log(*τ_U_* (*x*)). For low *α*, mean and variance scale like *µ*(*x*)^2^.

This suggests then that the time taken to reach fitness gain *f* (*x*) is dominated by the last step. We can write log(*τ_U_* (*f*)) as

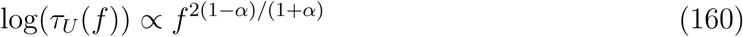

for the time taken to reach a fitness gain *f*. This is too slow to be a reasonable model of evolution in most scenarios. However, this result is very sensitive to the Gaussian tails of our random fitness functions. It may be possible to find distributions with alternate extremal statistics that give similar *f* (*x*) but faster *τ_U_*: we leave this for future research.

#### D.2 Exponential time correlation

In this section we develop the results needed to analyze exponentially time-correlated landscapes. Consider the general case of a set of mean-zero Gaussian random variables {*Z*(*y, t*)}, with correlations given by

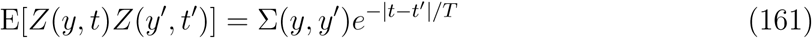

Here *t* and *t′* denote times, while *y* and *y′* denote some other type of coordinate (spatial, genetic, etc.).

The exponential structure simplifies the process of making a “time update” across some set of *y*. More concretely, suppose that the values of the *Z*(*y, t*_1_) are known at the time *t*_1_ for all *y* in some set *Y*. Suppose we now want to draw *Z*(*y, t*_2_) conditioned on *Z*(*y^′^, t*_1_) (across all *y^′^* in *Y* including *y*), for some *t*_2_ *> t*_1_. The conditional mean of *Z*(*y, t*_2_) is given by:

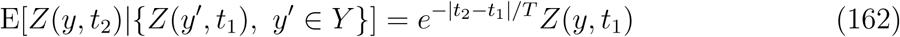

The expected value in the future depends only on the previously observed value at that location.

Equation 162 can be verified by using Equation 17, with *Z* in Equation 17 corresponding to *Z*(*y, t*_2_) (the unknown random variables) in the notation of this section, and similarly *W* corresponding to *Z*(*y, t*_1_) (the known random variables). Note that Equation 17 is satisfied if

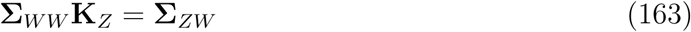

With our identifications we have

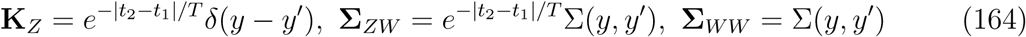

for *y, y^′^ ∈ Y*, which trivially satisfies the condition. Then Equation 162 comes from matrix multiplying (convolving) **K***_Z_* with the vector of known values *Z*(*y, t*_1_).

The conditional covariance across a pair *y* and *y′* in *Y* at time *t*_2_ can be written as:

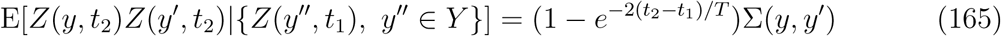

using Equation 18 with

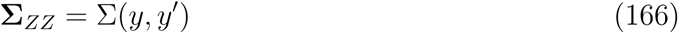

for *y, y′ ∈ Y* (which also comes from our correspondence).

This means that we can write *Z*(*y, t*_2_) as

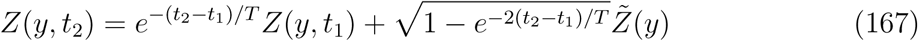

where the 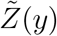 are Gaussian random variables defined over the *y* in *Y*, independent of all of the *Z*(*y, t*_1_), with covariance Σ(*y, y′*). In other words, the effect of the passage of time is to add an independently drawn equal-time slice to the last observed equal-time slice (with appropriate weighting). (This decomposition is possible since the sum of Gaussian random variables is also Gaussian, and the decomposition has the same mean and correlation as *Z*(*y, t*_2_).)

Equation 167 holds even if other values of *Z*(*y, t′*) are known for time or times *t^′^ < t*_1_ (including if an unequal number of observations were made for different *y*). This means that a “space update” is easy as well: if we need the value *Z*(*x, t*_1_) for some *x* not in *Y*, given the observations *Z*(*y, t*_1_) for all *y* in *Y* (before performing the time update to *t*_2_), we can draw the new value the way we would without time-dependence (that is, conditioning only on the *Z*(*y, t*_1_) via the time-independent covariance function Σ).

The above analysis gives an efficient algorithm to simulate evolutionary dynamics on a landscape with exponential time correlations. Given a timestep Δ*t* and the function *p_U_* (*µ*) for the probability of an uphill step given the current mean available step *µ*, with current mutation number *x* and time coordinate *t*, we proceed as follows:

1. Check if evolution finds an uphill step in time Δt (probability *p_U_ (µ(x, t))*Δ*t*).
2. If an uphill step is found: generate the new step, *s(x, t)*, and new mean available step, *µ(x+1, t)*, as in time-independent case (conditioned on *µ(y, t)* and *s(y, t)* for previously observed *y*).
3. Generate new, random *µ̄(y)* and *s̄(y)* according to the time-independent divergence, D(l), independent of all previous observations (generated for all observed y, including *s(x, t)* and *µ(x + 1, t)* if uphill step was taken).
4. Set 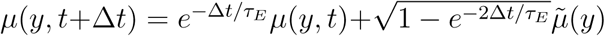 for all observed y (and equivalently for *s(y, t + Δt)*).
5. Return to step 1.

As Δ*t* goes to 0, the algorithm is exact. Note that the landscape update steps 3 *−* 5 are exact due to the analysis above; the only approximation is that the mutation occurs at the start of the Δ*t* interval. In practice, for Δ*t ≪ τ_E_* and Δ*t ≪ τ_U_*, where *τ_U_*is the typical time to an uphill step, the algorithm is quite accurate.

It is also efficient. Every uphill step with exponential time correlations takes the same amount of computation as the time-independent case does. The additional cost is the updates to the landscape when uphill steps are not taken. For average time 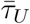 for an uphill step, this introduces an extra cost of 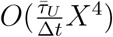 to simulate an evolution with *X* uphill steps.

This simple algorithm is specific to the case of exponential time correlations with no extra spatial structure; for general time correlations all past observations will play a role and the computational complexity increases rapidly with time.

#### D.3 Power law landscapes with exponential time correlations

After a transient period, dynamics in a landscape with exponentially time-decaying correlations leads to an equilibration of the DFE average *µ* around some critical value *µ_c_*. If the landscape has power law correlations, we must solve an integral equation to find *µ_c_*.

For slowly varying landscapes, *m ≡ −µ_c_/σ ≫* 1 and the average values of *s* and *µ* will control the overall dynamics. We assume that the time for each uphill step is not too broadly distributed near the steady state. Using the response kernel asymptotics, we have the approximate differential equation

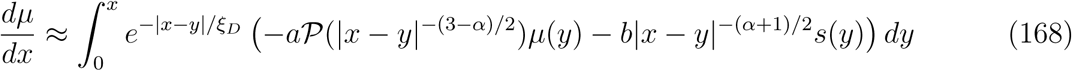

with

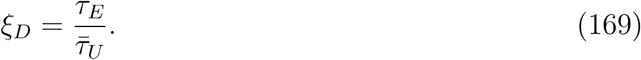

Here we average over the new random part of the landscape that came from the time-dependence of the landscape. The derivative vanishes at steady state. If we replace *µ*(*y*) with *µ_c_*and *s*(*y*) with 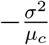, and take the limit of large *x*, the integrals are well approximated by Γ functions and we get

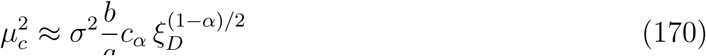

for some constant *c_α_*.

#### D.4 Heterogeneity in mutation waiting times

For power-law correlated seascapes with exponential time correlations, mutations can be roughly divided into two classes: “fast” mutations whose waiting time is much less than *τ_E_* (so the landscape is effectively constant between mutation events), and “slow” mutations which occur after the landscape has changed significantly (but before total decorrelation over time *τ_E_*).

In order to understand the two classes we define *r*(*t*) to be the probability, per time, of a beneficial mutation occurring a time *t* after the previous mutation. We define *µ*(*x*) as the average-available fitness step, at the time just after the previous step was taken. For convenience, we also define the scaled minus-average (over the steady state) *m ≡ −µ_c_/σ ≫* 1 and its scaled variance *W ≡* Var[*µ*(*x*)]*/σ*^2^, so that the quantity *h*(*x*) *≡* (*µ_c_ − µ*(*x*))*/σ* has mean zero and variance *W*. For very slowly varying landscapes, *τ_E_≫* 1, so that *m* will be large (although only logarithmically so with 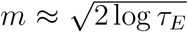, due to the Gaussian nature of the seascape.) We expect that *W* will be of order unity with contributions both from *V_η_*(*x*) and the effects from conditioning on the past as in Equation 92. (Note that *h* is not exactly Gaussian due to the distributions of the earlier *s*(*x*), but the deviations will not change the overall conclusions.)

For *t ≪ τ_E_*, from 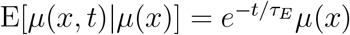 we have 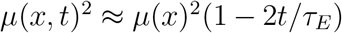 where we have dropped the stochastic part of the change 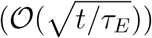 which can be self-consistently shown to be negligible from the analysis below. We thus have

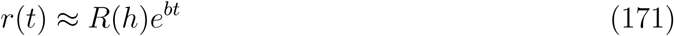

with

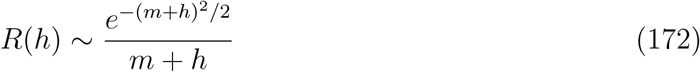

and

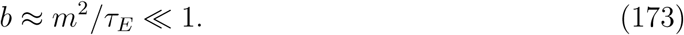

The first beneficial mutation typically occurs after a time *τ_U_* where the integrated probability of a mutation occurring is 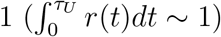. This gets us 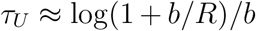. For not too large −*µ*(*x*) (that is, if *R*(*h*) ≫ *b*), the time-dependence of the landscape does not matter and *τ_U_* ∼ 1/*R*(*h*). The next mutation is very likely to occur before the environment changes by enough to make a difference.

For anomalously large *−µ*(*x*), a beneficial mutation is likely to occur only after the environment has started to change, although still well before it is decorrelated. This regime occurs for *R*(*h*) *≪ b* corresponding to *h > H* with *R*(*H*) = *b*: *h* even slightly above *H* leads to large *b/R* since *R* depends exponentially on (*h* + *m*)^2^. The waiting time is proportional to how much *−µ*(*x, t*) has to decrease for the mutation to occur. It is likely to occur after a waiting time near

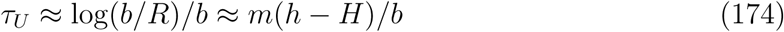

(the *−*[*h − H*]^2^*/*2 in log(*R*) can be neglected). We s*_√_*hall show that such waiting only occurs for a very small fraction of cases — i.e. that 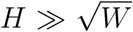. Thus the probability that *h > H* is of order 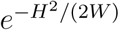 and its distribution has an exponential tail: *e^−^*^(^*^h−H^*^)^*^H/W^*. The average waiting time 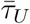 is thus dominated by the rare “slightly-stuck” situations with *h − H ∼ W/H* so that

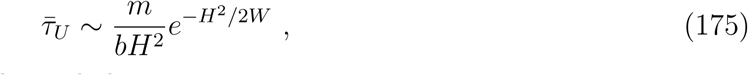

where we have used *H ≪ m* as shown below.

For power-law correlated landscapes, the effects of past history (Equation 58) gives us *τ*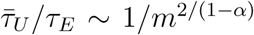. Thus for consistency we must have 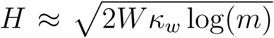 with *k_w_* = (1 + α) (1 − α). This justifies both *H ≪ m* and 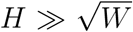. The typical *τ_U_* (*x*) are broadly distributed with

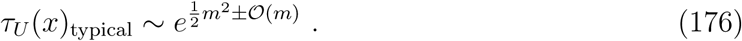

But since *H* is given by exp[(*m* + *H*)^2^*/*2] = *b* = *m*^2^*/τ_E_*, *τ_U_*(*x*)_typical_ is much less than *τ_E_*by a factor of

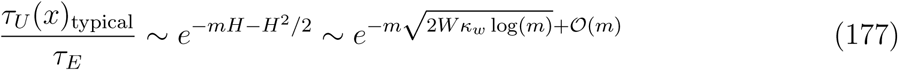

(ignoring multiplicative powers of *m*). Moreover, even the delay times for the rare slightly-stuck subset of steps that wait for the environment to change and dominate the average 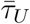, are much smaller than *τ_E_*

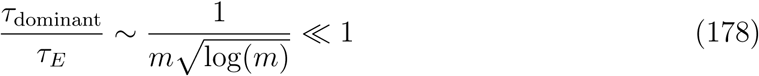

showing that during the waiting time the environment has changed but is only slightly decorrelated. This means that in one decorrelation time, *τ_E_*, the total number of steps taken is well approximated by 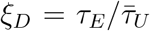 with smaller variations around this. The dynamics is thus quite smooth on time scales of *τ_E_* and length scales of *ξ_D_*. This justifies the analysis in Appendix D.3, which assumed such smoothness.

On shorter timescales the dynamics is very heterogenous, with the effects of time-dependence negligible except for a small (*∼* 1*/m^κw^*) fraction of steps. The dynamics alternates between rapid adaptation in a static environment, and a process where the population “waits” for more new directions to open up due to changes in the environment.

## Footnotes

1 This is, up to an additive constant, the negative of the *γ*-measure from [17].

## Bibliography

[1] Claudia Bank, Sebastian Matuszewski, Ryan T. Hietpas, and Jeffrey D. Jensen. On the (un)predictability of a large intragenic fitness landscape. Proceedings of the National Academy of Sciences, 113(49):14085–14090, December 2016. ISSN 0027-8424, 1091-6490. doi: 10.1073/pnas.1612676113.

[2] Jeffrey E. Barrick, Dong Su Yu, Sung Ho Yoon, Haeyoung Jeong, Tae Kwang Oh, Dominique Schneider, Richard E. Lenski, and Jihyun F. Kim. Genome evolution and adaptation in a long-term experiment with Escherichia coli. Nature, 461(7268):1243–1247, October 2009. ISSN 0028-0836. doi: 10.1038/nature08480.

[3] Jesús Blazquez, María-Isabel Morosini, María-Cristina Negri, and Fernando Baquero. Selection of Naturally Occurring Extended-Spectrum TEM *β*-Lactamase Variants by Fluctuating *β*-Lactam Pressure. Antimicrobial Agents and Chemotherapy, 44(8):2182–2184, January 2000. ISSN 0066-4804, 1098-6596. doi: 10.1128/AAC.44.8.2182-2184.2000.

[4] Joshua S. Bloom, Ian M. Ehrenreich, Wesley T. Loo, Thuy-Lan Vo Lite, and Leonid Kruglyak. Finding the sources of missing heritability in a yeast cross. Nature, 494 (7436):234–237, February 2013. ISSN 0028-0836. doi: 10.1038/nature11867.

[5] Joshua S. Bloom, Iulia Kotenko, Meru J. Sadhu, Sebastian Treusch, Frank W. Albert, and Leonid Kruglyak. Genetic interactions contribute less than additive effects to quantitative trait variation in yeast. Nature communications, 6, 2015.

[6] Zachary D. Blount, Christina Z. Borland, and Richard E. Lenski. Historical contingency and the evolution of a key innovation in an experimental population of Escherichia coli. Proceedings of the National Academy of Sciences, 105(23):7899–7906, June 2008. ISSN 0027-8424, 1091-6490. doi: 10.1073/pnas.0803151105.

[7] Alan J. Bray and David S. Dean. Statistics of Critical Points of Gaussian Fields on Large-Dimensional Spaces. Physical Review Letters, 98(15):150201, April 2007. doi: 10.1103/PhysRevLett.98.150201.

[8] L. F. Cugliandolo and J. Kurchan. Analytical solution of the off-equilibrium dynamics of a long-range spin-glass model. Physical Review Letters, 71(1):173–176, July 1993. doi: 10.1103/PhysRevLett.71.173.

[9] Leticia F. Cugliandolo and Pierre Le Doussal. Large time nonequilibrium dynamics of a particle in a random potential. Physical Review E, 53(2):1525–1552, February 1996. doi: 10.1103/PhysRevE.53.1525.

[10] J. Arjan GM de Visser and Joachim Krug. Empirical fitness landscapes and the predictability of evolution. Nature Reviews Genetics, 15(7):480–490, 2014.

[11] J. Arjan GM de Visser, Su-Chan Park, and Joachim Krug. Exploring the effect of sex on empirical fitness landscapes. The American Naturalist, 174(S1):S15–S30, 2009.

[12] Michael M. Desai and Daniel S. Fisher. Beneficial mutation–selection balance and the effect of linkage on positive selection. Genetics, 176(3):1759–1798, 2007.

[13] Michael M. Desai, Aleksandra M. Walczak, and Daniel S. Fisher. Genetic diversity and the structure of genealogies in rapidly adapting populations. Genetics, 193(2):565–585, 2013.

[14] David Deutscher, Isaac Meilijson, Martin Kupiec, and Eytan Ruppin. Multiple knockout analysis of genetic robustness in the yeast metabolic network. Nature Genetics, 38(9): 993–998, September 2006. ISSN 1546-1718. doi: 10.1038/ng1856.

[15] Júlia Domingo, Guillaume Diss, and Ben Lehner. Pairwise and higher-order genetic interactions during the evolution of a tRNA. Nature, 558(7708):117–121, June 2018. ISSN 1476-4687. doi: 10.1038/s41586-018-0170-7.

[16] Jeremy A. Draghi and Joshua B. Plotkin. Selection Biases the Prevalence and Type of Epistasis Along Adaptive Trajectories. Evolution, 67(11):3120–3131, 2013. ISSN 1558-5646. doi: 10.1111/evo.12192.

[17] Luca Ferretti, Benjamin Schmiegelt, Daniel Weinreich, Atsushi Yamauchi, Yutaka Kobayashi, Fumio Tajima, and Guillaume Achaz. Measuring epistasis in fitness landscapes: The correlation of fitness effects of mutations. Journal of Theoretical Biology, 396:132–143, May 2016. ISSN 0022-5193. doi: 10.1016/j.jtbi.2016.01.037.

[18] Daniel S. Fisher. Asexual evolution waves: Fluctuations and universality. Journal of Statistical Mechanics: Theory and Experiment, 2013(01):P01011, 2013.

[19] S. A. Frank. Generative models versus underlying symmetries to explain biological pattern. Journal of Evolutionary Biology, 27(6):1172–1178, June 2014. ISSN 1420-9101. doi: 10.1111/Jeb.12388.

[20] Lizhi Ian Gong, Marc A Suchard, and Jesse D Bloom. Stability-mediated epistasis constrains the evolution of an influenza protein. eLife, 2:e00631, May 2013. ISSN 2050-084X. doi: 10.7554/eLife.00631.

[21] Benjamin H. Good and Michael M. Desai. The impact of macroscopic epistasis on long-term evolutionary dynamics. Genetics, 199(1):177–190, 2015. ISSN 0016-6731. doi: 10.1534/genetics.114.172460.

[22] Benjamin H. Good, Igor M. Rouzine, Daniel J. Balick, Oskar Hallatschek, and Michael M. Desai. Distribution of fixed beneficial mutations and the rate of adaptation in asexual populations. Proceedings of the National Academy of Sciences, February 2012. ISSN 0027-8424, 1091-6490. doi: 10.1073/pnas.1119910109.

[23] Benjamin H. Good, Jeffrey E. Barrick, Michael J. McDonald, Michael M. Desai, and Richard E. Lenski. The dynamics of molecular evolution over 60,000 generations. Nature, 551(7678):45, October 2017. ISSN 1476-4687. doi: 10.1038/nature24287.

[24] Devin Greene and Kristina Crona. The Changing Geometry of a Fitness Landscape Along an Adaptive Walk. PLOS Computational Biology, 10(5):e1003520, May 2014. ISSN 1553-7358. doi: 10.1371/journal.pcbi.1003520.

[25] Nelson G. Hairston and Theresa A. Dillon. Fluctuating Selection and Response in a Population of Freshwater Copepods. Evolution, 44(7):1796–1805, November 1990. ISSN 1558-5646. doi: 10.1111/j.1558-5646.1990.tb05250.x.

[26] Alex R. Hall, Pauline D. Scanlan, Andrew D. Morgan, and Angus Buckling. Host–parasite coevolutionary arms races give way to fluctuating selection. Ecology Letters, 14(7):635–642, July 2011. ISSN 1461-0248. doi: 10.1111/j.1461-0248.2011.01624.x.

[27] David W. Hall, Matthew Agan, and Sara C. Pope. Fitness Epistasis among 6 Biosynthetic Loci in the Budding Yeast Saccharomyces cerevisiae. Journal of Heredity, 101 (suppl 1):S75–S84, January 2010. ISSN 0022-1503, 1465-7333. doi: 10.1093/jhered/esq007.

[28] Peter Hegarty and Anders Martinsson. On the existence of accessible paths in various models of fitness landscapes. The Annals of Applied Probability, 24(4):1375–1395, 2014. ISSN 1050-5164.

[29] Sungmin Hwang, Benjamin Schmiegelt, Luca Ferretti, and Joachim Krug. Universality Classes of Interaction Structures for NK Fitness Landscapes. Journal of Statistical Physics, 172(1):226–278, July 2018. ISSN 1572-9613. doi: 10.1007/s10955-018-1979-z.

[30] Mark Isalan, Caroline Lemerle, Konstantinos Michalodimitrakis, Carsten Horn, Pedro Beltrao, Emanuele Raineri, Mireia Garriga-Canut, and Luis Serrano. Evolvability and hierarchy in rewired bacterial gene networks. Nature, 452(7189):840–845, April 2008. ISSN 0028-0836. doi: 10.1038/nature06847.

[31] Kavita Jain, Joachim Krug, and Su-Chan Park. Evolutionary Advantage of Small Populations on Complex Fitness Landscapes. Evolution, 65(7):1945–1955, 2011. ISSN 1558-5646. doi: 10.1111/j.1558-5646.2011.01280.x.

[32] Elizabeth R. Jerison, Sergey Kryazhimskiy, James Kameron Mitchell, Joshua S. Bloom, Leonid Kruglyak, and Michael M. Desai. Genetic variation in adaptability and pleiotropy in budding yeast. eLife, 6:e27167, 2017.

[33] José I. Jiménez, Ramon Xulvi-Brunet, Gregory W. Campbell, Rebecca Turk-MacLeod, and Irene A. Chen. Comprehensive experimental fitness landscape and evolutionary network for small RNA. Proceedings of the National Academy of Sciences, 110(37): 14984–14989, October 2013. ISSN 0027-8424, 1091-6490. doi: 10.1073/pnas.1307604110.

[34] Rees Kassen and Thomas Bataillon. Distribution of fitness effects among beneficial mutations before selection in experimental populations of bacteria. Nature Genetics, 38 (4):484–488, April 2006. ISSN 1061-4036. doi: 10.1038/ng1751.

[35] Stuart A. Kauffman and Edward D. Weinberger. The NK model of rugged fitness landscapes and its application to maturation of the immune response. Journal of Theoretical Biology, 141(2):211–245, November 1989. ISSN 0022-5193. doi: 10.1016/S0022-5193(89)80019-0.

[36] Jorge Kurchan and Laurent Laloux. Phase space geometry and slow dynamics. Journal of Physics A: Mathematical and General, 29(9):1929, 1996. ISSN 0305-4470. doi: 10.1088/0305-4470/29/9/009.

[37] Richard E. Lenski, Michael R. Rose, Suzanne C. Simpson, and Scott C. Tadler. Long-term experimental evolution in Escherichia coli. I. Adaptation and divergence during 2,000 generations. The American Naturalist, pages 1315–1341, 1991.

[38] Sasha F Levy, Jamie R Blundell, Sandeep Venkataram, Dmitri A Petrov, Daniel S Fisher, and Gavin Sherlock. Quantitative evolutionary dynamics using high-resolution lineage tracking. Nature, 519(7542):181–186, 2015. doi: 10.1038/nature14279.

[39] Chuan Li, Wenfeng Qian, Calum J. Maclean, and Jianzhi Zhang. The fitness landscape of a tRNA gene. Science, 352(6287):837–840, May 2016. ISSN 0036-8075, 1095-9203. doi: 10.1126/science.aae0568.

[40] Yuping Li, Sandeep Venkataram, Atish Agarwala, Barbara Dunn, Dmitri A. Petrov, Gavin Sherlock, and Daniel S. Fisher. Hidden Complexity of Yeast Adaptation under Simple Evolutionary Conditions. Current Biology, 28(4):515–525.e6, February 2018. ISSN 0960-9822. doi: 10.1016/j.cub.2018.01.009.

[41] Raymond H. Y. Louie, Kevin J. Kaczorowski, John P. Barton, Arup K. Chakraborty, and Matthew R. McKay. Fitness landscape of the human immunodeficiency virus envelope protein that is targeted by antibodies. Proceedings of the National Academy of Sciences, 115(4):E564–E573, January 2018. ISSN 0027-8424, 1091-6490. doi: 10.1073/pnas.1717765115.

[42] Debora S. Marks, Lucy J. Colwell, Robert Sheridan, Thomas A. Hopf, Andrea Pagnani, Riccardo Zecchina, and Chris Sander. Protein 3D Structure Computed from Evolutionary Sequence Variation. PLOS ONE, 6(12):e28766, December 2011. ISSN 1932-6203. doi: 10.1371/journal.pone.0028766.

[43] Debora S. Marks, Thomas A. Hopf, and Chris Sander. Protein structure prediction from sequence variation. Nature Biotechnology, 30(11):1072–1080, November 2012. ISSN 1546-1696. doi: 10.1038/nbt.2419.

[44] Joshua K. Michener and Christina D. Smolke. High-throughput enzyme evolution in Saccharomyces cerevisiae using a synthetic RNA switch. Metabolic Engineering, 14(4): 306–316, July 2012. ISSN 1096-7176. doi: 10.1016/j.ymben.2012.04.004.

[45] Ville Mustonen and Michael Lässig. Fitness flux and ubiquity of adaptive evolution. Proceedings of the National Academy of Sciences, 107(9):4248–4253, March 2010. ISSN 0027-8424, 1091-6490. doi: 10.1073/pnas.0907953107.

[46] Richard A. Neher and Oskar Hallatschek. Genealogies of rapidly adapting populations. Proceedings of the National Academy of Sciences, 110(2):437–442, January 2013. ISSN 0027-8424, 1091-6490. doi: 10.1073/pnas.1213113110.

[47] Richard A. Neher and Boris I. Shraiman. Statistical genetics and evolution of quantitative traits. Reviews of Modern Physics, 83(4):1283, 2011.

[48] Johannes Neidhart, Ivan G. Szendro, and Joachim Krug. Exact results for amplitude spectra of fitness landscapes. Journal of Theoretical Biology, 332:218–227, September 2013. ISSN 0022-5193. doi: 10.1016/j.jtbi.2013.05.002.

[49] Johannes Neidhart, Ivan G. Szendro, and Joachim Krug. Adaptation in Tunably Rugged Fitness Landscapes: The Rough Mount Fuji Model. Genetics, 198(2):699–721, October 2014. ISSN 0016-6731, 1943-2631. doi: 10.1534/genetics.114.167668.

[50] S. Nowak and J. Krug. Accessibility percolation on n-trees. EPL (Europhysics Letters), 101(6):66004, 2013. ISSN 0295-5075. doi: 10.1209/0295-5075/101/66004.

[51] Stefan Nowak and Joachim Krug. Analysis of adaptive walks on NK fitness landscapes with different interaction schemes. Journal of Statistical Mechanics: Theory and Experiment, 2015(6):P06014, 2015.

[52] Timothy Nugent and David T. Jones. Accurate de novo structure prediction of large transmembrane protein domains using fragment-assembly and correlated mutation analysis. Proceedings of the National Academy of Sciences, 109(24):E1540–E1547, June 2012. ISSN 0027-8424, 1091-6490. doi: 10.1073/pnas.1120036109.

[53] H. Allen Orr. The Distribution of Fitness Effects Among Beneficial Mutations. Genetics, 163(4):1519–1526, April 2003. ISSN 0016-6731, 1943-2631.

[54] Giorgio Parisi. On the statistical properties of the large time zero temperature dynamics of the SK model. In Scaling And Disordered Systems, pages 161–171. World Scientific, 2002.

[55] Su-Chan Park and Joachim Krug. *δ*-exceedance records and random adaptive walks. Journal of Physics A: Mathematical and Theoretical, 49(31):315601, 2016.

[56] Su-Chan Park, Ivan G. Szendro, Johannes Neidhart, and Joachim Krug. Phase transition in random adaptive walks on correlated fitness landscapes. Physical Review E, 91 (4):042707, 2015.

[57] Su-Chan Park, Johannes Neidhart, and Joachim Krug. Greedy adaptive walks on a correlated fitness landscape. Journal of theoretical biology, 397:89–102, 2016.

[58] Patrick C. Phillips. Epistasis—the essential role of gene interactions in the structure and evolution of genetic systems. Nature Reviews Genetics, 9(11):855–867, 2008.

[59] Frank J. Poelwijk, Vinod Krishna, and Rama Ranganathan. The Context-Dependence of Mutations: A Linkage of Formalisms. PLOS Computational Biology, 12(6):e1004771, June 2016. ISSN 1553-7358. doi: 10.1371/journal.pcbi.1004771.

[60] Victoria O. Pokusaeva, Dinara R. Usmanova, Ekaterina V. Putintseva, Lorena Espinar, Karen S. Sarkisyan, Alexander S. Mishin, Natalya S. Bogatyreva, Dmitry N. Ivankov, Arseniy V. Akopyan, Sergey Ya Avvakumov, Inna S. Povolotskaya, Guillaume J. Filion, Lucas B. Carey, and Fyodor A. Kondrashov. An experimental assay of the interactions of amino acids from orthologous sequences shaping a complex fitness landscape. PLOS Genetics, 15(4):e1008079, April 2019. ISSN 1553-7404. doi: 10.1371/journal.pgen.1008079.

[61] Esteban Real, Sherry Moore, Andrew Selle, Saurabh Saxena, Yutaka Leon Suematsu, Quoc Le, and Alex Kurakin. Large-scale evolution of image classifiers. arXiv preprint arXiv:1703.01041, 2017.

[62] Daniel E. Rozen, Michelle G. J. L. Habets, Andreas Handel, and J. Arjan G. M. de Visser. Heterogeneous Adaptive Trajectories of Small Populations on Complex Fitness Landscapes. PLOS ONE, 3(3):e1715, March 2008. ISSN 1932-6203. doi: 10.1371/journal.pone.0001715.

[63] Zachary R. Sailer and Michael J. Harms. High-order epistasis shapes evolutionary trajectories. PLOS Computational Biology, 13(5):e1005541, May 2017. ISSN 1553-7358. doi: 10.1371/journal.pcbi.1005541.

[64] David Sherrington and Scott Kirkpatrick. Solvable model of a spin-glass. Phys. Rev. Lett., 35:1792–1796, Dec 1975. doi: 10.1103/PhysRevLett.35.1792.

[65] Tong Si, Jiazhang Lian, and Huimin Zhao. Strain Development by Whole-Cell Directed Evolution. In Directed Enzyme Evolution: Advances and Applications, pages 173–200. Springer, Cham, 2017. ISBN 978-3-319-50411-7 978-3-319-50413-1. doi: 10.1007/978-3-319-50413-17.

[66] Peter F. Stadler and Robert Happel. Random field models for fitness landscapes. Journal of Mathematical Biology, 38(5):435–478, 1999.

[67] Ivan G. Szendro, Martijn F. Schenk, Jasper Franke, Joachim Krug, and J. Arjan GM De Visser. Quantitative analyses of empirical fitness landscapes. Journal of Statistical Mechanics: Theory and Experiment, 2013(01):P01005, 2013.

[68] Athanasios Typas, Robert J. Nichols, Deborah A. Siegele, Michael Shales, Sean R. Collins, Bentley Lim, Hannes Braberg, Natsuko Yamamoto, Rikiya Takeuchi, Barry L. Wanner, Hirotada Mori, Jonathan S. Weissman, Nevan J. Krogan, and Carol A. Gross. High-throughput, quantitative analyses of genetic interactions in *E. coli*. Nature Methods, 5(9):781–787, September 2008. ISSN 1548-7105. doi: 10.1038/nmeth.1240.

[69] Sandeep Venkataram, Barbara Dunn, Yuping Li, Atish Agarwala, Jessica Chang, Emily R. Ebel, Kerry Geiler-Samerotte, Lucas Herissant, Jamie R. Blundell, Sasha F. Levy, Daniel S. Fisher, Gavin Sherlock, and Dmitri A. Petrov. Development of a Comprehensive Genotype-to-Fitness Map of Adaptation-Driving Mutations in Yeast. Cell, 166(6):1585–1596.e22, September 2016. ISSN 0092-8674, 1097-4172. doi: 10.1016/j.cell.2016.08.002.

[70] E. D. Weinberger. Fourier and Taylor series on fitness landscapes. Biological Cybernetics, 65(5):321–330, September 1991. ISSN 1432-0770. doi: 10.1007/BF00216965.

[71] Daniel M. Weinreich, Nigel F. Delaney, Mark A. Depristo, and Daniel L. Hartl. Darwinian evolution can follow only very few mutational paths to fitter proteins. Science (New York, N.Y.), 312(5770):111–114, April 2006. ISSN 1095-9203. doi: 10.1126/science.1123539.

[72] Daniel B. Weissman, Michael M. Desai, Daniel S. Fisher, and Marcus W. Feldman. The rate at which asexual populations cross fitness valleys. Theoretical population biology, 75(4):286–300, 2009.

[73] Michael J. Wiser, Noah Ribeck, and Richard E. Lenski. Long-Term Dynamics of Adaptation in Asexual Populations. Science, 342(6164):1364–1367, December 2013. ISSN 0036-8075, 1095-9203. doi: 10.1126/science.1243357.

[74] Alex Wong. Epistasis and the Evolution of Antimicrobial Resistance. Frontiers in Microbiology, 8, February 2017. ISSN 1664-302X. doi: 10.3389/fmicb.2017.00246.

[75] Nicholas C Wu, Lei Dai, C Anders Olson, James O Lloyd-Smith, and Ren Sun. Adaptation in protein fitness landscapes is facilitated by indirect paths. eLife, 5:e16965, July 2016. ISSN 2050-084X. doi: 10.7554/eLife.16965.

